# Repeated gain and loss of a single gene modulates the evolution of vascular pathogen lifestyles

**DOI:** 10.1101/2020.04.24.058529

**Authors:** Emile Gluck-Thaler, Aude Cerutti, Alvaro Perez-Quintero, Jules Butchacas, Verónica Roman-Reyna, Vishnu Narayanan Madhaven, Deepak Shantharaj, Marcus V. Merfa, Céline Pesce, Alain Jauneau, Taca Vancheva, Jillian M. Lang, Caitilyn Allen, Valerie Verdier, Lionel Gagnevin, Boris Szurek, Sébastien Cunnac, Gregg Beckham, Leonardo de la Fuente, Hitendra Kumar Patel, Ramesh V Sonti, Claude Bragard, Jan E. Leach, Laurent D. Noël, Jason C. Slot, Ralf Koebnik, Jonathan M. Jacobs

## Abstract

Vascular pathogens travel long distances through host veins leading to life-threatening, systemic infections. In contrast, non-vascular pathogens remain restricted to infection sites, triggering localized symptom development. The contrasting features of vascular and non-vascular diseases suggest distinct etiologies, but the basis for each remains unclear. Here, we show that the hydrolase CbsA acts as a phenotypic switch between vascular and non-vascular plant pathogenesis. *cbsA* was enriched in genomes of vascular phytopathogenic bacteria in the Xanthomonadaceae family and absent in most non-vascular species. CbsA expression allowed non-vascular *Xanthomonas* to cause vascular blight while *cbsA* mutagenesis resulted in reduction of vascular or enhanced non-vascular symptom development. Phylogenetic hypothesis testing further revealed that *cbsA* was lost in multiple non-vascular lineages and more recently gained by some vascular subgroups, suggesting that vascular pathogenesis is ancestral. Our results overall demonstrate how the gain and loss of single loci can facilitate the evolution of complex ecological traits.

## Introduction

Pathogenic microorganisms cause diseases of animals and plants. Some pathogenic species colonize the host vasculature, which leads to systemic infection, while others remain localized to non-vascular tissues. Complex structural and biochemical differences between vascular and non-vascular tissues suggest that pathogens have multiple distinct adaptations to either environment, yet the genetic and evolutionary bases of such adaptations are largely unknown.

Adaptations often occur through wholesale gain and loss of specific genes, resulting in more rapid evolution compared with incremental changes at the DNA sequence level alone (*1*). In bacteria, gene gain occurs primarily through horizontal gene transfer while gene loss or pseudogenization occurs through multiple mechanisms, including transposon-mediated insertions and sequence deletions in open reading frames (*2*–*4*). Especially for loci encoding ecologically relevant traits, gene gain and loss effectively act as phenotypic switches, enabling rapid shifts between what otherwise seem like complex lifestyles (*3*). For example, transitions between plant pathogenic and commensal *Pseudomonas* (*5*), transitions between mutualist and parasitic phenotypes in nitrogen-fixing bacteria (*6*, *7*) and transitions between mutualistic and plant pathogenic *Rhodococcus* (*8*) have all been shown to reproducibly occur through the gain and loss of genomic islands containing multiple genes all contributing to the same phenotype. Such rapid evolutionary dynamics have profound implications for our understanding of disease ecology and disease management strategies.

In plants, vascular xylem and non-vascular parenchyma tissues represent distinct niches. Xylem is comprised of dead cells with highly reinforced walls organized into cylinders that provide plants with structural integrity and a means of long-distance fluid transport. In contrast, parenchyma tissues are composed of living cells and gas filled intercellular spaces. Xylem fluid consists primarily of water and mineral nutrients, and is thought to be nutrient limiting, although many vascular pathogens can use it to reach high densities (*9*). Xylem tissue runs throughout the plant, enabling the distribution of water from roots to leaves, but also serving as a potential pathway for rapid, systemic transport of pathogens.

*Xanthomonas* (Gammaproteobacteria) is diverse genus of plant-associated Gram-negative bacteria that cause vascular and non-vascular diseases of over 200 monocot and dicot plant hosts (*10*). *Xanthomonas* species are separated into subgroups called pathovars (pv.) based on their phenotypic behavior such as symptom development (e.g. vascular or non-vascular) or host range (*10*). Vascular xanthomonads invade the water transporting xylem; non-vascular *Xanthomonas* species cause localized symptoms by colonizing the mesophyll. Although often closely related, the genetic determinants distinguishing vascular from non-vascular *Xanthomonas* lineages at the intraspecific level are not clear.

Here, we used *Xanthomonas* as a model to study the etiology of plant vascular pathogenesis because this genus contains multiple independent pairs of strains from the same species that cause either vascular or non-vascular diseases. This enabled us to disentangle genetic features that are shared due to ancestry and those that may be shared due to common tissue-specific lifestyles. Given the tendency of bacteria to evolve through the gain and loss of genes organized into clusters or genomic islands, we hypothesized that vascular and non-vascular pathogenesis emerge through the gain and loss of small numbers of linked loci. Surprisingly, we found evidence supporting the most extreme version of this hypothesis, where transitions between vascular and non-vascular lifestyles are mediated by the repeated gain and loss of a single gene that acts as a phenotypic switch.

## Results

### *cbsA* is significantly associated with vascular pathogenesis

We first identified high priority candidate genes associated with transitions to vascular and non-vascular lifestyles. We classified predicted proteins from 59 publicly available whole genome sequences of *Xanthomonas* and *Xylella* species into ortholog groups (OGs). We then conducted an analysis of trait evolution across a SNP-based phylogeny where for each OG we tested the hypothesis that transitions to vascular or non-vascular lifestyles were dependent on that OG’s presence or absence (Figure 1). The phylogenetic relationships between vascular and non-vascular pathovars indicated that xylem pathogenesis is paraphyletic, i.e., not limited to a single clade, an individual Xanthomonas sp., or host plant genus (Figure 1; Supplemental Figure 1&2). Instead, vascular diseases of many host plant families are caused by different pathovars across the *Xanthomonas* genus. We identified two OGs whose presence was strongly associated (Log Bayes Factor >10) with the distribution of tissue-specific lifestyles (Figure 1; Supplemental Figure 1, Supplemental Table 1&2). One OG (OG0003492) was highly associated with vascular pathogenesis, while the other (OG0002818) was associated with non-vascular pathogenesis. For this study, we focused on vascular pathogen-enriched OG0003492, which encodes a cell wall degrading cellobiohydrolase (EC 3.2.1.4, glycosylhydrolase family GH6) called CbsA (*11*, *12*). Next, phylogenetic analysis of CbsA sequences revealed that distinct monophyletic lineages within this gene family are alternatively found in either vascular or non-vascular pathogen genomes (Figure 1B; Supplemental Figure 3). Within *Xanthomonas*, CbsA sequences form two distinct clades: the first contains sequences found in both vascular and non-vascular pathogen genomes, and the second contains sequences found exclusively in vascular pathogen genomes. All vascular pathogens with a CbsA homolog found in the first clade also possess a CbsA homolog found in the second clade, effectively possessing two copies of the CbsA gene (Figure 1B, Supplemental Figure 3). The association of specific CbsA clades with specific pathogen lifestyles, in addition to its occasional presence in multiple copies in vascular pathogen genomes, suggests that CbsA sequences found in either of these two clades have distinct biological functions, with sequences that are exclusive to vascular pathogens likely contributing to vascular pathogenesis.

**Figure 1.**
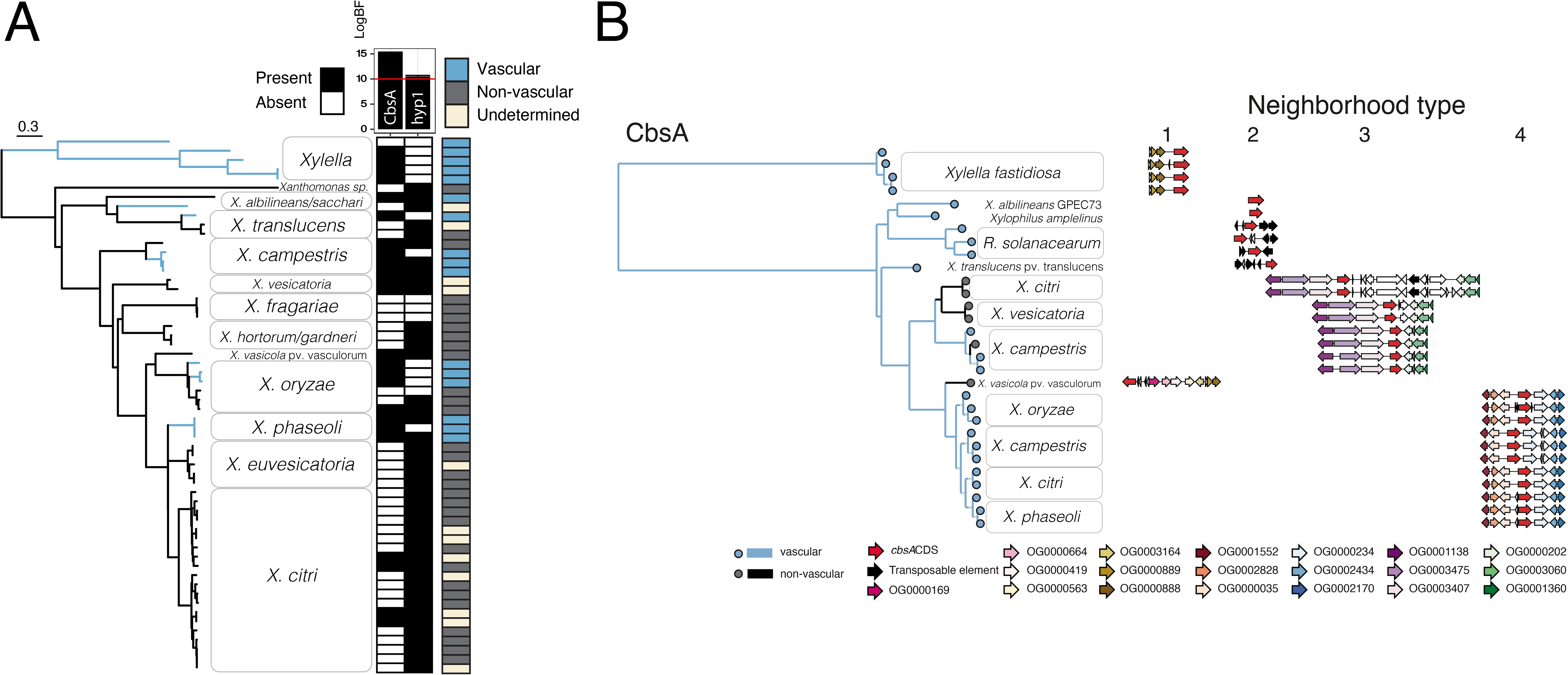
The cellobiohydrolase CbsA is associated with transitions to vascular pathogenic lifestyles in Gram-negative pathogens. A) Highest-ranking associations between ortholog group presence/absences and evolutionary transitions between vascular and non-vascular lifestyles in the *Xanthomonadaceae*. A genome-based SNP phylogeny is shown to the left, with strains from the same species condensed into clades. Classifications of each strain as vascular (blue), non-vascular (yellow) or unknown (gray) are depicted to the right of each tip, followed by a heatmap summarizing, for each strain, the presence (black) or absence (white) of the two gene ortholog groups whose distributions are most strongly supported to be dependent on vascular lifestyle status (determined by model testing through the ranking of log Bayes Factors; Methods). Additional figure details can be found in Supplemental Figures 1&5. B) A phylogenetic tree based on CbsA amino acid sequences from strains with whole genome sequences found in (A), where branches on the tree are color coded according to pathogenic lifestyle. To the right of each tip is a schematic depicting the neighborhood type in which that particular *cbsA* sequence is found, where the four possible neighborhood types are defined based on conserved synteny (indicated by color-coded gene models corresponding to specific ortholog groups). Vascular bacteria possess *cbsA* homologs located in type 1, 2, and 4 neighborhoods, while non-vascular bacteria possess *cbsA* homologs found primarily in type 3 neighborhoods. Note that strains of the vascular pathogen *X. campestris* pv. campestris have two copies of *cbsA* located in either type 3 or type 4 neighborhoods.

### Heterologous expression of *cbsA* bestows vascular pathogenesis to a non-vascular pathogen

Because *cbsA* was present in vascular and largely absent from non-vascular *Xanthomonas* species, we hypothesized that *cbsA* was either: A) gained by vascular *Xanthomonas* species or B) lost by non-vascular *Xanthomonas* species. To experimentally test the alternate models, we examined the effects of manipulating *cbsA* on the contrasting tissue-specific behavior of two closely related barley pathogens from the same species: vascular *Xanthomonas translucens* pvs. translucens (Xtt) and non-vascular undulosa (Xtu).

Xtt and Xtu both cause non-vascular bacterial leaf streak (BLS) disease of barley (*13*). However, only Xtt can colonize the xylem which leads to long distance bacterial blight (BB) symptom development (Figure 2A-C) (*13*, *14*). Upon leaf clipping, only Xtt produces distant vascular BB; meanwhile Xtu symptoms remain near the site of inoculation (Figure 2A). Moreover, Xtt strains contain an intact copy of *cbsA*, while *X. translucens* pv. undulosa contains a copy of *cbsA* that is disrupted in the 5’ region by a transposase (Supplemental Figure 4).

**Figure 2.**
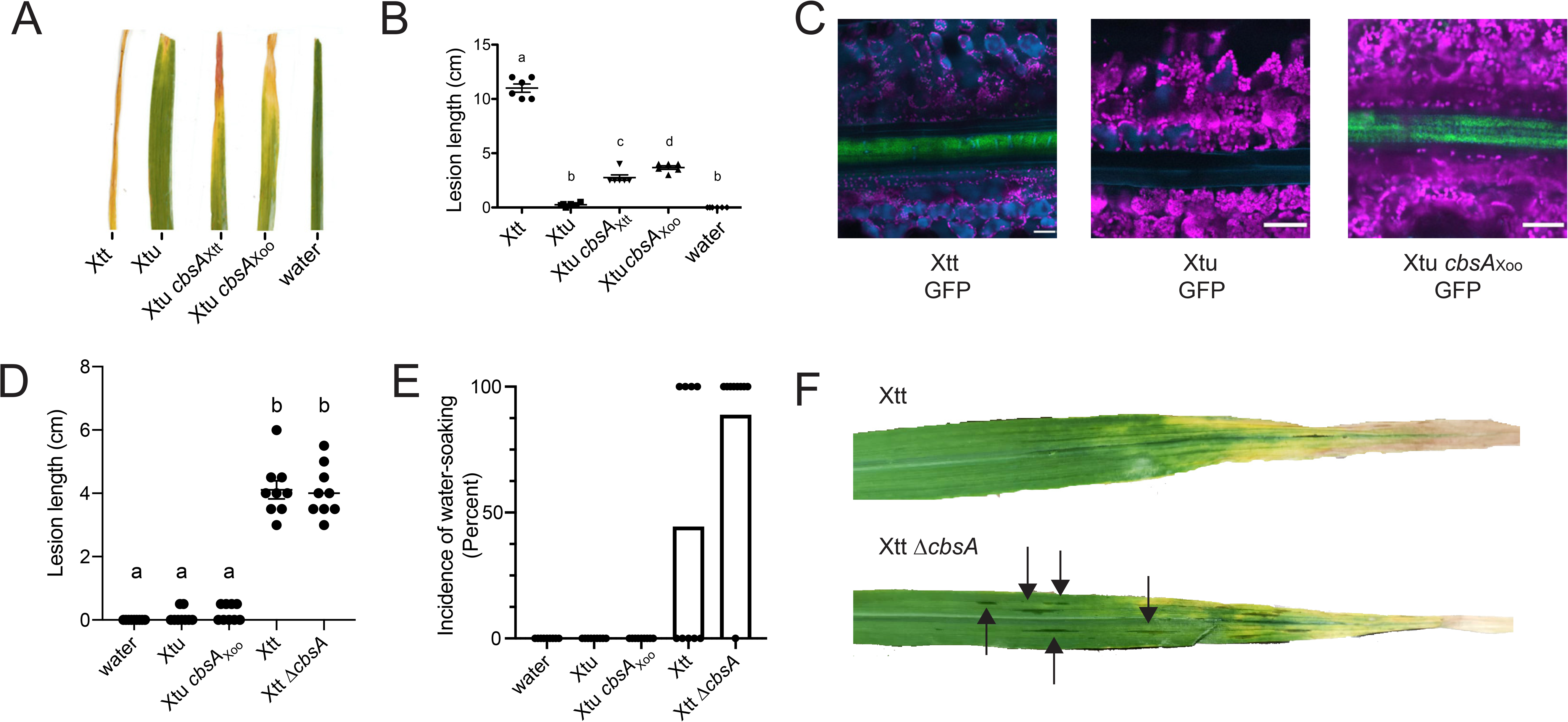
Experimental gain and loss of CbsA facilitates transitions between vascular and non-vascular pathogenic lifestyles. A) Addition of either *cbsA* from vascular *X. translucens* pv. translucens (Xtt) or *cbsA* from vascular *Xanthomonas oryzae* pv. oryzae (Xoo) to non-vascular *X. translucens* pv. undulosa (Xtu) permits development of chlorotic lesions indicative of vascular disease on barley 21 days post-inoculation (dpi) B) Corresponding vascular lesion lengths, with significant differences among treatments indicated by a-d (n = 6, *P*<0.02) C) Representative confocal images of vascular bundles downstream of leaf lesions on barley 12 dpi with green fluorescent protein (GFP) transformed strains demonstrate gain of vascular colonization by Xtu *cbsA*_Xoo_. Green indicates bacterial cells expressing GFP; magenta indicates chlorophyll autofluorescence outlining non-vascular mesophyll cells; cyan indicates autofluorescence outlining xylem cell walls or phenylpropanoid accumulation in mesophyll cells. D-E) Lesion lengths or incidence of non-vascular water soaked lesions were quantified after barley leaf clipping 14 dpi with Xtt Δ*cbsA*. Bars in E) represent percent leaves showing symptoms with dots included to display individual leaf lesions incidence. F) Images of symptomatic barley leaves infected with Xtt and Xtt Δ*cbsA*, where water soaked lesions are indicated with black arrows indicating non-vascular symptom development.

As Xtt possesses *cbsA* while Xtu lacks an intact copy, we tested if the expression of CbsA promotes vascular symptom development in Xtu. Xtu miniTn*7*::*cbsA*_Xtt_, a single insertion variant with an intact copy of *cbsA* from Xtt, caused distant leaf lesions of approximately 4.5 cm (Figure 2A-B). Moreover, expression of the characterized CbsA ortholog from the vascular rice pathogen Xoo (Xtu miniTn*7*::*cbsA*_Xoo_) also permitted Xtu to cause distant symptom development consistent with a vascular pathogenic lifestyle. Using GFP-expressing strains, we reproducibly observed Xtu miniTn7::*cbsA*_Xoo_ inside the xylem similar to Xtt (Figure 2C). Wild-type Xtu did not produce vascular symptoms and was not detected in distant xylem vessels (Figure 2C). Therefore, the gain of *cbsA* from either of two different vascular pathogens is sufficient to promote xylem-mediated colonization and distant infection of leaves by non-vascular Xtu.

### Impact of *cbsA* mutagenesis on vascular pathogenesis is dependent on genetic background

We found that the Xtt Δ*cbsA* mutant was still capable of causing vascular leaf blight, suggesting other unknown factors support vascular pathogenesis beyond CbsA alone (Figure 2D&F). However, while Xtt Δ*cbsA* could still cause systemic symptom development, the mutation of this cellulase altered this strain’s pathogenic behavior by promoting the development of non-vascular, water-soaked lesions downstream of the xylem blight on 90% of infected leaves compared with only 10% of leaves on plants infected with wild-type vascular Xtt (Figure 2E&F). These water-soaked symptoms are typical of non-vascular disease development in Xtt and Xtu (*13*, *14*). Therefore, while vascular disease development is not completely abolished by *cbsA* mutagenesis, the absence of *cbsA* increased the development of non-vascular disease symptoms.

These results did not match previous reports that *cbsA* deletion mutants in *X. oryzae* pv. oryzae and *R. solanacearum* have reduced systemic virulence and vascular pathogenesis (*15*, *16*). We therefore replicated and expanded upon these previous findings by mutating *cbsA* in *Xanthomonas oryzae* pv. oryzae and *Xylella fastidiosa* (Xanthomonadaceae). *X. oryzae* pv. oryzae causes bacterial blight of rice with systemic symptoms similar to Xtt on barley. *Xylella fastidiosa*, an insect-vectored, xylem pathogen, is the causal agent of Pierce’s disease of grape and the emerging olive quick decline disease. *X. oryzae* pv. oyzae and *X. fastidiosa* deletion mutants were severely reduced in vascular symptom development, confirming and building upon previous reports (Supplemental Figure 5)(*15*). The variable effects of mutagenizing *cbsA* in Xtt versus *X. oryzae* pv. oyzae and *X. fastidiosa* indicate that the robustness of vascular phenotypes is lineage dependent within *Xanthomonas*, with certain species likely possessing multiple determinants in addition to *cbsA* that contribute to vascular pathogenesis.

### The genomic location of *cbsA* alternates between four distinct neighborhoods

Across all examined genomes, *cbsA* is found embedded in one of four genomic neighborhood types with conserved gene synteny (Figure 1B&3). The localization of *Xylella fastidiosa*’s and *X. vasicola’s cbsA* in type 1 neighborhoods, combined with a lack of evidence suggesting horizontal gene transfer between these two species (Figure 1B), provides support that *cbsA* was present and organized in a type 1 context in the last common ancestor of *Xanthomonas* and *Xylella*. Based on this inference, it is likely that *cbsA* was then re-located into type 2, type 3 and type 4 neighborhoods through separate cis-transposition events as *Xanthomonas spp*. diversified. The timing of transposition events 3 and 4 are uncertain due to lack of resolution in species-level relationships, but likely occur near to where indicated on the species tree (Figure 3). Within the gamma-proteobacteria, all known vascular pathogens in our dataset have a copy of *cbsA* localized in the context of type 1, 2, or 4 neighborhoods. Within *Xanthomonas*, sequences from the clade of CbsA homologs found in both vascular and non-vascular pathogens are located in type 3 neighborhoods, while sequences from the clade of CbsA homologs found exclusively in vascular pathogen genomes are located in type 4 neighborhoods, further supporting the hypothesis that sequences belonging to either of these two clades have separate functions (Figure 2).

**Figure 3.**
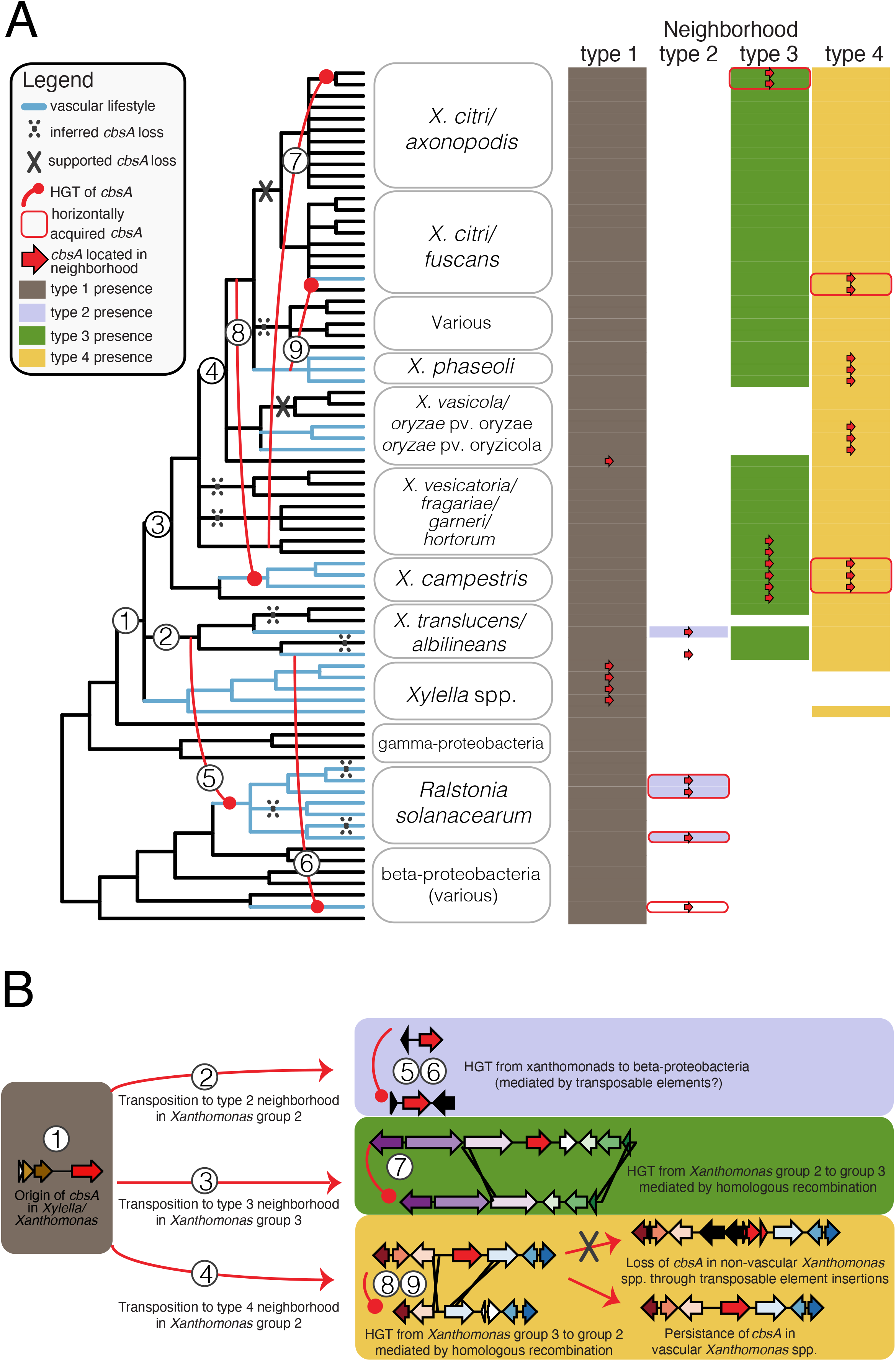
Repeated horizontal transfer, transposition, and gene loss events drive the distribution of *cbsA* in gram-negative bacteria. A) A 50% majority rule consensus tree summarizing 81 conserved single copy ortholog trees is shown to the left, with the names of the 75 individual isolates consolidated into relevant taxonomic groupings. Inferred horizontal gene transfer (HGT), transposition, and loss events are drawn and numbered on the tree, and further described in B). The matrix to the right of tree indicates the presence/absence of one of four distinct genomic neighborhood types (shaded/unshaded cells) in which *cbsA* homologs are found within a given genome (presence of *cbsA* indicated by an overlaid red arrow). Note that in many cases, all of the constituent genes making up a specific neighborhood are present in a given genome save for *cbsA* (indicated by the absence of an overlaid red arrow). This tree has been lightly edited for viewing purposes by removing several taxa from outside the Xanthamonadales, and can be viewed in its entirety in Supplemental Figure 3. B) The sequence of inferred evolutionary events drawn onto the tree in A). Genomic neighborhood types are represented by schematics where gene models are color-coded according to ortholog group. The color-coding of neighborhood types is consistent across both panels.

### *cbsA* has been independently gained by lineages now displaying vascular lifestyles

*cbsA* and varying lengths of adjacent sequence experienced three horizontal transfers in the *Xanthomonas* genus mediated by homologous recombination events in flanking gene neighborhoods (events 7,8,9 in Figure 3, Supplemental Figure 6-8). Two transfers from what was likely the ancestor of the vascular pathogen *X. phaseoli* are coincident with the emergence of vascular lifestyles in xylem-adapted *X. campestris* pv. campestris and *X. citri* pv. phaseoli, and occurred within the context of type 4 neighborhoods (events 8 and 9 Figure 3; Supplemental Figure 6-8). The third transfer occurred in the context of a type 3 neighborhood, where neither the donor lineage of *X. vesicatoria* nor the recipient lineage of *X. citri* have been reported to be capable of vascular pathogenesis.

### *cbsA* was horizontally transferred from vascular gamma-to beta-proteobacteria

We found additional evidence that *cbsA* was horizontally transferred from gamma-proteobacterial Xanthomonadaceae to the beta-proteobacterial xylem plant pathogens *R. solanacearum* and *Xylophilus ampelinus* (Figure 3). *cbsA* sequences in both *X. transluscens* pv. transluscens and *R. solanacearum* are flanked on one or both sides by transposable elements (Figure 1B), providing a plausible mechanism for mediating horizontal transfer through transposition between these distant lineages. However, we could not test this specific hypothesis with confidence because the phylogenies of the transposable elements in question are complex and contain signatures of extensive horizontal transfer between strains.

### *cbsA* has been repeatedly lost from lineages now displaying non-vascular lifestyles

At least 10 losses of *cbsA* are required to parsimoniously explain its distribution across the beta- and gamma-proteobacteria when taking into account all HGT events supported by phylogenetic hypothesis testing (Figure 3; Supplemental Tables 4-6). While the majority of losses are inferred using parsimony criteria (e.g. losses in non-vascular strains of *X. hortorum* and *X. fragariae;* Methods), several *cbsA* pseudogenes present in extant species directly support the hypothesis of repeated, independent losses through distinct inactivation mechanisms. For example, *cbsA* was independently pseudogenized in the non-vascular *X. translucens* pv. undulosa and *X. sacchari* through sequence deletions in its 5’ coding region (Supplemental Figure 4&6). In contrast, transposable elements have disrupted the 5’ region of *cbsA* in non-vascular *X. oryzae* pv. oryzicola, and are present in the type 4 neighborhoods of certain non-vascular *X. citri* subsp. citri and *X. fuscans* subsp. aurantifolii isolates that lack a copy of *cbsA* (Supplemental Figure 6). These examples of multiple, independent disruptions to *cbsA* in lineages displaying non-vascular lifestyles suggest that non-vascular pathogenesis convergently evolved through repeated gene loss.

## Discussion

Systemic pathogens traverse host veins to move long distances, leading to life-threatening systemic infections. In contrast, non-vascular pathogens remain restricted to the site of infection, triggering localized symptom development with far fewer implications for host health. Although complex differences between these modes of infection suggest they have radically different origins, the results we present here suggest that vascular and non-vascular pathogenesis are two points on an evolutionary continuum, a finding with important implications for understanding and predicting pathogen evolution (Figure 4). By integrating comparative genomic, phylogenetic, and functional genetic analyses, we found evidence that vascular and non-vascular plant pathogenic lifestyles emerge from the repeated gain and loss of a single gene that can act as a phenotypic switch.

**Figure 4.**
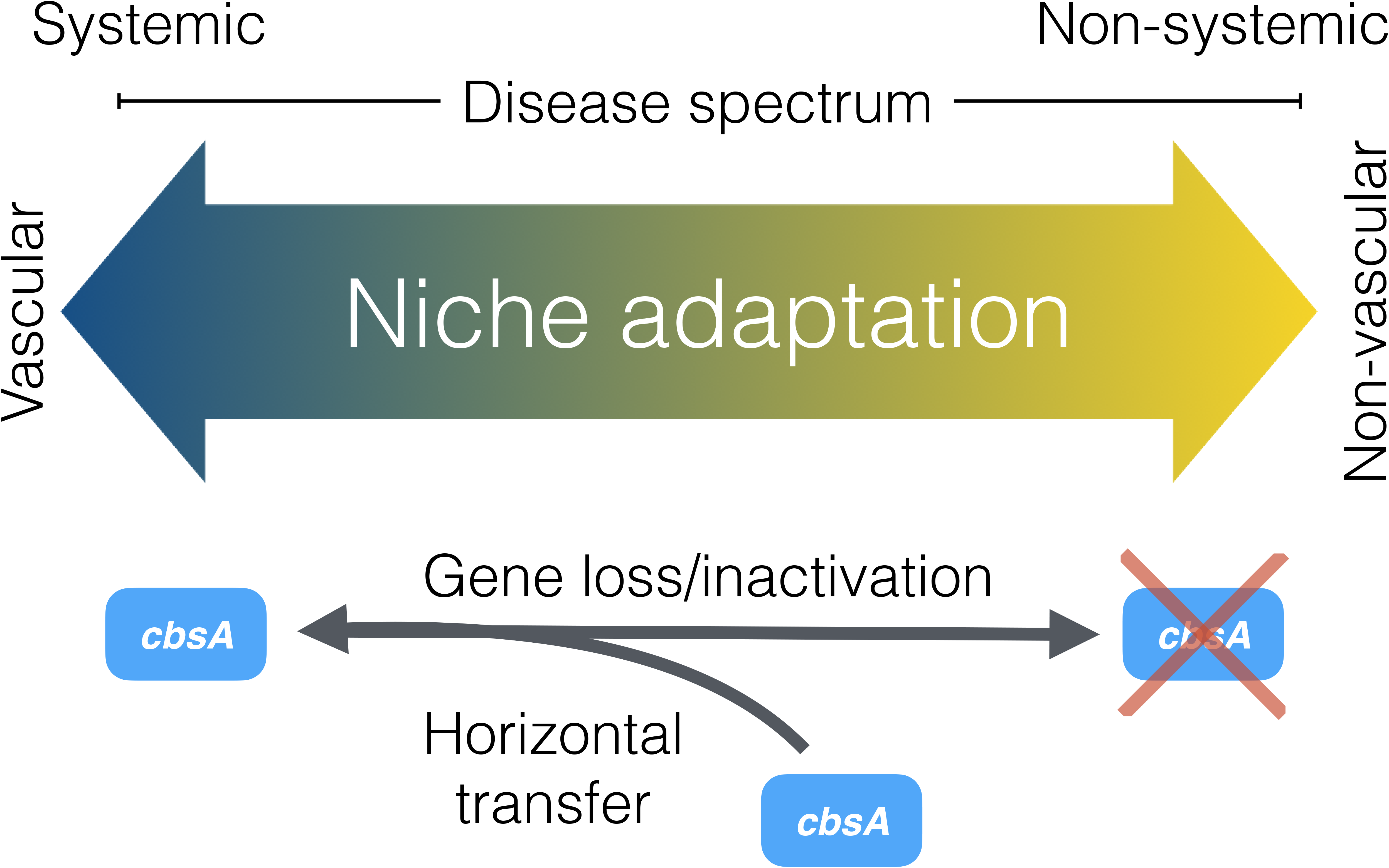
The evolution of vascular and non-vascular pathogenesis in plant-associated *Xanthomonas* bacteria is driven by the gain and loss of *cbsA*. Our combined phenotypic and phylogenetic analyses support a model where vascular and non-vascular pathogenesis exist as two points on the same evolutionary continuum that is traversed by either the acquisition or loss of a single cellobiohydrolase, *cbsA*.

Our functional and phylogenetic results suggest that *cbsA* contributes to the evolution of *Xanthomonas* vascular pathogenicity, but to varying extent depending on the species considered. Xylem-specific pathogens, including *X. fastidiosa, X. oryzae* pv. oryzae and *R. solanacearum*, require CbsA for vascular pathogenesis, whereas Xtt, which induces both vascular and non-vascular disease symptoms, appears to use other factors beyond CbsA to colonize xylem vasculature. That the phenotypic outcomes of CbsA acquisition are dependent on genetic background suggests that there exist multiple evolutionary routes to vascular pathogenesis, and highlights the particularities of specific host-pathogen interactions. Nevertheless, the preponderance of phenotypic and phylogenetic evidence supports the hypothesis that *cbsA* was present in the last common ancestor of *Xanthomonas* and *Xylella*, has since played not only a historical but possibly a contemporary role in driving the emergence and re-emergence of tissue-specific behavior in the Xanthomonadaceae.

While we document repeated gains and losses of *cbsA*, the conditions that favor phenotypes resulting from either its presence or absence remain to be determined. Although *cbsA* homologs are among the highest expressed genes during xylem pathogenesis (*9*, *17*), and are required for vascular pathogenesis in several species (Supplemental Figure 5), the contributions of CbsA to pathogen fitness remain unclear. Current theory suggests that there may be a fitness cost to retaining this gene and the vascular lifestyle it enables, given that CbsA induces immune responses and can prime the plant against *Xanthomonas* infection (*15*). Furthermore, cell wall degradation products, such as the CbsA enzymatic biproduct cellobiose, could act as a danger-associated molecular pattern in the plant mesophyll and may induce plant defenses through WRKY transcription factors (*18*). We therefore speculate that *cbsA’s* absence may be selected for to dampen recognition by the host and/or the elicitation of host immunity; however, these hypotheses remain to be tested.

Gene loss is a fundamental mechanism of adaptation (*19*). Especially for loci with large effects such as *cbsA*, only a minimal number of loss events are required to incur drastic changes to phenotype. Adaptive phenotypes arising through loss of function may emerge over shorter timescales compared with adaptive phenotypes arising through gains in function, as genes typically have more mutational opportunities for losing functions than for gaining functions (*20*). Even within our own limited dataset, we observed multiple mutational routes in the form of sequence deletions and transposable element insertions that led to the convergent loss of *cbsA* in different non-vascular pathogen lineages, which suggests that non-vascular phenotypes readily emerge in the Xanthomonadaceae.

Although there may be fewer mutational routes for gaining gene functions compared with losing them, our phylogenetic analyses revealed that rates of gain and loss may be balanced by latent patterns in genome architecture, such as the conservation of synteny. Homologous recombination in bacteria is typically studied within species, and is considered to be important for maintaining genetic diversity in what would otherwise be clonal lineages (*21*). Less considered are the impacts of homologous recombination across species. Our results add to a growing body of literature suggesting that, while perhaps less common than intraspecific homologous recombination (*22*, *23*), interspecific gene exchange facilitated by homologous recombination at syntenic loci is an important mechanism of adaptation (*24*). All three *cbsA* HGT events within *Xanthomonas* occurred through homologous recombination in syntenic neighborhoods flanking *cbsA* presence/absence polymorphisms, and two of these resulted in the reversal of an ancestral loss event (Figure 2), suggesting that synteny conservation potentiates not only gene gain but the reversal of lineage-specific gene loss. By effectively increasing an individual strain’s ability to access cross-species pan-genomic material, the conservation of synteny is likely to be an important accelerator of ecological adaptation.

Overall, our study provides an integrated evolutionary and functional framework for studying the genetic bases of transitions between vascular and non-vascular pathogen lifestyles (Figure 4). Our experiments demonstrate that the acquisition of *cbsA* is sufficient for long-distance systemic pathogenesis in specific *Xanthomonas* pathogens. Conversely, the loss of *cbsA*, while not necessary to abolish vascular disease development, is sufficient for the development of non-vascular disease symptoms. We add to a growing body of literature that suggests that transitions between distinct bacterial ecotypes may be mediated by the recurrent gain and loss of few loci (*5*, *8*). Although it remains to be determined how the processes of rapid gene gain and loss impact vascular and non-vascular evolution in other pathogenic microbes, our work suggests that these evolutionary events play an important role in shaping bacterial adaptation to specific host tissues.

## Materials and Methods

### Comparative genomics for identification of vascular pathogen-specific genes

Using Orthofinder v2.2.3 (*25*), we first created ortholog groups (OGs) from all predicted amino acid sequences derived from 171 complete and 8 partially complete publicly available assemblies from the Xanthomonadaceae and representative lineages across the beta- and gamma-proteobacteria in order to obtain a comprehensive comparative genomic dataset (Table S1). Consensus functional annotations for each OG were obtained by determining the most frequent protein family domain present among the members of the OG using InterProScan version 5.25-64.0 (*26*). Predicted proteins across all genomes were classified into 36,905 OGs using Orthofinder (Supplemental Table 2) (*25*)

Genomes were classified as vascular, non-vascular or unknown based on available information in the literature (Supplemental Table 1). The *Xanthomonas* species included xylem and parenchyma pathogens that infect diverse dicot and monocot crops such as rice, wheat, barley, cabbage, tomato, citrus and common bean. A distant vascular grape and citrus Xanthomonadaceae bacterium, *Xylella fastidiosa*, was also analyzed.

For analyses limited to the Xanthamonadaceae, we built a more resolved SNP-based parsimony tree using kSNP3 (*27*) from a set of publicly available complete and annotated genomes from different species in the Xanthomonadaceae family (optimum kmer size = 21; Supplemental Table 1). Using the kSNP3 as a reference, associations were identified between the presence/absence of each ortholog group in the analyzed genomes and the vascular/non-vascular trait using BayesTraitsV3 (*28*). The likelihood that both traits (vascularity vs. gene presence) evolved independently was compared to the likelihood they evolved dependently. Evidence of dependent evolution was assessed as Log Bayes Factors = 2(log marginal likelihood dependent model – log marginal likelihood independent model), and genes were considered to have strong evidence of dependent evolution with a Log Bayes Factor >10.

### Bacterial strains and growth conditions

The bacterial strains used in this study are listed in Table S7. *Escherichia coli* strains were grown at 37°C in Luria-Bertani medium. *X. translucens* or *X. oryzae* cells were grown at 28°C on solid or liquid nutrient broth or peptone-sucrose rich media (*14*). When necessary, media were supplemented with gentamicin (15 μg/ml), kanamycin (25 μg/ml) or spectinomycin (50 μg/ml). See Table S7 for specific strains used in this study.

### Recombinant DNA techniques

Total genomic and plasmid DNA were isolated by standard methods. *E. coli* and *Xanthomons* species were transformed as previously described (*14*). To construct complementation vectors of *cbsA*_Xtt_ and *cbsA*_Xoo_, the gene regions including the native promoters were PCR-amplified from *X. translucens* pv. tranlsucens str. UPB886. Each were cloned into pUC18miniTn7T to create pUC18miniTn7T::*cbsA*_Xtt_ and pUC18miniTn7T::*cbsA4*_Xoo_ (*29*). For gene expression, *X. translucens* pv. undulosa strains were transformed with miniTn*7* plasmids and pTNS1 to promote transposition and single gene insertion, and each was confirmed as described (*29*). We were unable to insert *cbsA* via miniTn*7 X. translucens* pv. translucens strain UPB886. We therefore sequenced *X. translucens* pv. translucens Δ*cbsA* with long read PacBio SMRT sequencing (Supplemental Figure 9). There were no notable differences in sequence between wild-type UPB886 and the Δ*cbsA* mutant. For visualization of bacteria by fluorescence microscopy, *Xanthomonas* bacteria (Table S7) were transformed with vectors for GFP expression (pNEO-GFP) (*30*). See Supplemental Tables 7 & 8 for specific strains and primers, respectively, used in this study.

### Plant growth conditions, inoculation methods and live imaging with confocal microscopy

Barley (*Hordeum vulgare* L. cv. Morex or Betzes) were grown in growth chambers with cycles of 16 hours of light per day at 22-24°C. Rice (*Oryza sativa* cv. Nipponbare) were grown in growth chambers with 16 hours of light per day at 28°C 70% relative humidity or in the greenhouse. Plant seeds were directly germinated in potting mix. For either barley or rice, one leaf per plant was inoculated by leaf-clipping 7-10 days after seeds were planted with a water-based inoculum (OD_600_=0.1) or water as a control as previously described (*14*). Disease symptoms were assessed using at least n = 5 replications per condition. Statistical differences were evaluated using the one-way ANOVA with Tukey’s multiple comparison test or Student’s *t*-test when appropriate. Symptom development was evaluated 21 days post-inoculation.

For bacterial localization, barley plant leaves were inoculated as above. Whole leaf tissue was imaged 5-14 days post inoculation with a Leica SP2 AOBS (Wetzlar, Germany) laser scanning confocal microscope with 40X oil objective. Barley leaves were cut directly adjacent to the inoculation zone for asymptomatic plants and immediately downstream of symptoms for symptomatic plants. Plant tissue was mounted onto a glass slide with water and covered with a glass coverslip. A 488 nm laser was used for GFP excitation and emitted fluorescence was collected between 505 and 540 nm. A 405 nm and 633 nm lasers were used for autofluorescence and emitted fluorescence was collected between 410 and 460 nm to define plant cell structures and between 650 and 700 nm for chlorophyll. Three to six plants were examined per biological replicate per treatment over three total biological replicates. Representative confocal images represent maximal projections calculated from 15 to 25 confocal planes acquired in the z dimension (increments of at least 0.5 mm).

### Phylogenetic analyses

To decrease redundancy among strain- or species-specific genomes in our dataset while maintaining sample power, we built a preliminary 50% majority-rule consensus tree based on the maximum likelihood (ML) phylogenies of 139 amino acid alignments of single copy orthologs. We used this tree to guide our selection of at most 3 representative genomes from each *Xanthomonas* pathovar, ultimately arriving at a final dataset of 86 genomes (Table S1). Using this de-replicated genomic dataset, we then built a final 50% majority-rule consensus tree based on 81 amino acid-based ML phylogenies of single copy orthologs that had greater than 60% average bootstrap support, our rationale being that consolidating multiple gene trees with high support increases the robustness of species-level phylogenetic inference (*31*). We rooted the final consensus tree at the bifurcation between the beta- and gamma-proteobacteria.

All nucleotide and amino acid alignments were generated using MAFFT v7.047 with options ‘--auto’ for automatic selection of best alignment strategy (*32*) and trimmed using trimAL v1.4 with options ‘-automated1’ for heuristic method selection and ‘-gt 0.25’ for removing all sites with gaps in ≥75% of sequences (*33*). Sequences with gaps in ≥30% of sites were removed. All ML trees were built using IQTREE v1.6.9 with option ‘-m MFP’ to find the best-fitting model of sequence evolution (*34*). Majority-rule consensus trees were built using RAxML v8.2.11 (*35*).

### Analysis of *cbsA* homologs and neighboring genomic regions

In order to determine the precise mechanisms and relative order through which *cbsA* was gained and lost from the genomes in our dataset, we analyzed the evolutionary history and structural features of all gene neighborhoods that flank *cbsA*. Using custom scripts, we first explored the gene neighborhoods surrounding *cbsA* homologs (+/- 15kb) in the 86-genome dataset for conserved synteny, as defined by orthogroup content conservation. For each of the four conserved neighborhood types that we identified, we then re-searched all genomes for regions composed of these genes, thus identifying all instances of each neighborhood in each genome, regardless of whether *cbsA* was present or not (Table S3). In doing so, we could then leverage phylogenetic evidence from flanking genes to support or reject competing hypotheses of gene duplications, horizontal gene transfers and losses that may have resulted in *cbsA’s* extant distribution.

We built nucleotide-based ML phylogenies of *cbsA* and the genes from each neighborhood type, and manually reconciled their evolutionary histories with the consensus species tree using a combination of parsimony-based gene tree-species tree reconciliation and likelihood-based phylogenetic testing (Supplemental Figures 6-8; Supplemental Tables 4-6). In order to robustly root the *cbsA* tree for reconciliation analysis, we first retrieved the top 1000 hits in the NCBI nr protein database (last accessed: 03/09/18) to the *cbsA* sequence in *X. campestris* (accession: WP_076057318) and used them to build a midpoint rooted ML tree (available on the Figshare repository). This tree was then used as a reference to root subsequent ML trees that focused only on this study’s clade of *cbsA* sequences of interest. We additionally built a ML tree with *cbsA* sequences from the full 179 genome dataset in order to verify the final topology of the *cbsA* tree built with the 86 genome de-replicated dataset (available on the Figshare repository). All other gene trees were midpoint rooted.

All genomic regions were further annotated for transposable elements with BLAST using the ISFinder database in order to ensure a comprehensive structural annotation of mobile elements (*36*). Nucleotide sequences of the genomic regions that were missing *cbsA* were searched using BLASTn with a *cbsA* query to ensure any missing or incomplete *cbsA* coding regions were identified. The Mixture Model and Hidden Markov Model from the PhyML package were used to detect homologous recombination breakpoints in the untrimmed nucleotide alignments that were then manually inspected and refined if necessary (*37*).

### Phylogenetic hypothesis testing

In each tree with a topology that suggested HGT, we compared the likelihood of the most likely tree obtained through a standard ML search (representing the hypothesis of HGT) with the likelihood of a constrained tree where sequences were forced to adhere to a topology that would be expected under a scenario of vertical inheritance (representing the hypothesis of no HGT). In this way, we could probabilistically assess whether a scenario of vertical inheritance or HGT best explained the observed sequence data. We used the approximately unbiased (AU) test with 100,000 re-samplings using the RELL method (*38*) as implemented in IQTREE v1.6.9 (*34*) to identify the most likely tree among a set of constrained and optimal trees. The null hypothesis that the constrained tree had the largest observed likelihood was rejected at α ≤ 0.05. Practically, this meant that we inferred HGT by showing that the constrained ML tree was significantly worse (smaller log likelihood) than the optimal ML tree. Constrained ML tree searches were conducted using IQTREE v1.6.9 (*34*) by supplying a trimmed nucleotide alignment and a non-comprehensive, multifurcating constraint tree specifying the monophyly of particular sequences of interest to which the resulting ML tree was forced to adhere to (Figures S4-6; see Tables S4-6 for all constraint criteria).

### Data visualization

All phylogenetic trees were visualized using ETE3 v3.0.0b32 (*39*). All genomic regions were visualized using Easyfig (*40*).

## Supporting information

Supplemental Text with Figures

## General

The authors are grateful to the French Xanthomonads Network, Jeffery Chang (Oregon State), Stephen Cohen (Ohio State) and Tiffany Lowe-Power (UC—Davis) for fruitful intellectual discussions.

## Funding

An NSF Postdoctoral Fellowship in Biology (1306196) to JMJ, a US Fulbright Scholar Award to Belgium to JMJ, a USDA-NIFA Postdoctoral Fellowship (2017-67012-26116) to JMJ, a COST SUSTAIN travel grant to JMJ and a NSF-NIFA joint PBI grant (2018-05040) to JMJ, JML and JEL. NSF (DEB-1638999) to JCS. A Fonds de Recherche du Quebec-Nature et Technologies Doctoral Research Scholarship to EGT. AJ and LDN are supported by the NEPHRON project (ANR-18-CE20-0020-01). This work was supported by a Ph.D. grant from the French Ministry of National Education and Research to AC. LIPM is part of the TULIP LabEx (ANR-10-LABX-41; ANR-11-IDEX-0002-02).

## Author contributions

JMJ conceptualized and JMJ and RK supervised the conducted research. EGT, AC and APQ equally conducted research and provided formal analysis. CP, TV and JML conducted additional research. JMJ, EGT, APQ, JS, LDN wrote the original draft, while RK, CA, LG, BS, SC, VV, JEL, CB and GB participated in reviewing and editing the manuscript.

## Competing interests

Authors declare no competing interests

## Data and materials availability

All data trimmed alignments, optimal and constrained maximum likelihood tree files, orthogroup assignments, and custom scripts are available on the Figshare data repository (DOI: 10.6084/m9.figshare.8218703).

## Notes

### Competing Interest Statement

The authors have declared no competing interest.

## References and Notes

1. J. Iranzo, Y. I. Wolf, E. V. Koonin, I. Sela, Gene gain and loss push prokaryotes beyond the homologous recombination barrier and accelerate genome sequence divergence. Nat. Commun. (2019), doi:10.1038/s41467-019-13429-2.

2. E. V. Koonin, Y. I. Wolf, Genomics of bacteria and archaea: The emerging dynamic view of the prokaryotic world. Nucleic Acids Res. (2008), doi:10.1093/nar/gkn668.

3. A. T. Maurelli, R. E. Fernández, C. A. Bloch, C. K. Rode, A. Fasano, “Black holes” and bacterial pathogenicity: A large genomic deletion that enhances the virulence of Shigella spp. and enteroinvasive Escherichia coli. Proc. Natl. Acad. Sci. U. S. A. 95, 3943–8 (1998).

4. C.-H. Kuo, H. Ochman, Deletional Bias across the Three Domains of Life. Genome Biol. Evol. 1, 145–62 (2009).

5. R. A. Melnyk, S. S. Hossain, C. H. Haney, Convergent gain and loss of genomic islands drive lifestyle changes in plant-associated Pseudomonas. ISME J. (2019), doi:10.1038/s41396-019-0372-5.

6. S. S. Porter, J. Faber-Hammond, A. P. Montoya, M. L. Friesen, C. Sackos, Dynamic genomic architecture of mutualistic cooperation in a wild population of Mesorhizobium. ISME J. (2019), doi:10.1038/s41396-018-0266-y.

7. K. G. Nandasena, G. W. O’Hara, R. P. Tiwari, J. G. Howieson, Rapid in situ evolution of nodulating strains for Biserrula pelecinus L. through lateral transfer of a symbiosis island from the original mesorhizobial inoculant. Appl. Environ. Microbiol. (2006), doi:10.1128/AEM.00889-06.

8. E. A. Savory, S. L. Fuller, A. J. Weisberg, W. J. Thomas, M. I. Gordon, D. M. Stevens, A. L. Creason, M. S. Belcher, M. Serdani, M. S. Wiseman, N. J. Grünwald, M. L. Putnam, J. H. Chang, Evolutionary transitions between beneficial and phytopathogenic rhodococcus challenge disease management. Elife (2017), doi:10.7554/eLife.30925.

9. J. M. Jacobs, L. Babujee, F. Meng, A. Milling, C. Allen, The in planta transcriptome of Ralstonia solanacearum: Conserved physiological and virulence strategies during bacterial wilt of tomato. MBio. 3 (2012), doi:10.1128/mBio.00114-12.

10. M.-A. Jacques, M. Arlat, A. Boulanger, T. Boureau, S. Carrère, S. Cesbron, N. W. G. Chen, S. Cociancich, A. Darrasse, N. Denancé, M. Fischer-Le Saux, L. Gagnevin, R. Koebnik, E. Lauber, L. D. Noël, I. Pieretti, P. Portier, O. Pruvost, A. Rieux, I. Robène, M. Royer, B. Szurek, V. Verdier, C. Vernière, Using Ecology, Physiology, and Genomics to Understand Host Specificity in Xanthomonas: French Network on Xanthomonads (FNX). Annu. Rev. Phytopathol. 54 (2016), doi:10.1146/annurev-phyto-080615-100147.

11. L. Tayi, S. Kumar, R. Nathawat, A. S. Haque, R. V. Maku, H. K. Patel, R. Sankaranarayanan, R. V. Sonti, A mutation in an exoglucanase of Xanthomonas oryzae pv. oryzae, which confers an endo mode of activity, affects bacterial virulence, but not the induction of immune responses, in rice. Mol. Plant Pathol. (2018), doi:10.1111/mpp.12620.

12. G. T. Beckham, J. Ståhlberg, B. C. Knott, M. E. Himmel, M. F. Crowley, M. Sandgren, M. Sørlie, C. M. Payne, Towards a molecular-level theory of carbohydrate processivity in glycoside hydrolases. Curr. Opin. Biotechnol. (2014),, doi:10.1016/j.copbio.2013.12.002.

13. C. Bragard, E. Singer, A. Alizadeh, L. Vauterin, H. Maraite, J. Swings, Xanthomonas translucens from Small Grains: Diversity and Phytopathological Relevance. Phytopathology. 87, 1111–1117 (1997).

14. C. Pesce, J. M. Jacobs, E. Berthelot, M. Perret, T. Vancheva, C. Bragard, R. Koebnik, Comparative Genomics Identifies a Novel Conserved Protein, HpaT, in Proteobacterial Type III Secretion Systems that Do Not Possess the Putative Translocon Protein HrpF. Front. Microbiol. 8, 1177 (2017).

15. G. Jha, R. Rajeshwari, R. V Sonti, Functional interplay between two Xanthomonas oryzae pv,. oryzae secretion systems in modulating virulence on rice. Mol. Plant. Microbe. Interact. 20, 31–40 (2007).

16. H. Liu, S. Zhang, M. a Schell, T. P. Denny, Pyramiding unmarked deletions in Ralstonia solanacearum shows that secreted proteins in addition to plant cell-wall-degrading enzymes contribute to virulence. Mol. Plant. Microbe. Interact. 18, 1296–1305 (2005).

17. J. F. González, G. Degrassi, G. Devescovi, D. De Vleesschauwer, M. Höfte, M. P. Myers, V. Venturi, A proteomic study of Xanthomonas oryzae pv. oryzae in rice xylem sap. J. Proteomics. 75, 5911–5919 (2012).

18. C. de A. Souza, S. Li, A. Z. Lin, F. Boutrot, G. Grossmann, C. Zipfel, S. C. Somerville, Cellulose-Derived Oligomers Act as Damage-Associated Molecular Patterns and Trigger Defense-Like Responses. Plant Physiol. (2017), doi:10.1104/pp.16.01680.

19. R. Albalat, C. Cañestro, Evolution by gene loss. Nat. Rev. Genet. (2016),, doi:10.1038/nrg.2016.39.

20. A. K. Hottes, P. L. Freddolino, A. Khare, Z. N. Donnell, J. C. Liu, S. Tavazoie, Bacterial Adaptation through loss of Function. PLoS Genet. (2013), doi:10.1371/journal.pgen.1003617.

21. B. J. Shapiro, J. Friedman, O. X. Cordero, S. P. Preheim, S. C. Timberlake, G. Szabó, M. F. Polz, E. J. Alm, Population genomics of early events in the ecological differentiation of bacteria. Science (80-.). (2012), doi:10.1126/science.1218198.

22. C. L. Huang, P. H. Pu, H. J. Huang, H. M. Sung, H. J. Liaw, Y. M. Chen, C. M. Chen, M. B. Huang, N. Osada, T. Gojobori, T. W. Pai, Y. T. Chen, C. C. Hwang, T. Y. Chiang, Ecological genomics in Xanthomonas: The nature of genetic adaptation with homologous recombination and host shifts. BMC Genomics (2015), doi:10.1186/s12864-015-1369-8.

23. N. Potnis, P. P. Kandel, M. V. Merfa, A. C. Retchless, J. K. Parker, D. C. Stenger, R. P. P. Almeida, M. Bergsma-Vlami, M. Westenberg, P. A. Cobine, L. De La Fuente, Patterns of inter- and intrasubspecific homologous recombination inform eco-evolutionary dynamics of Xylella fastidiosa. ISME J. (2019), doi:10.1038/s41396-019-0423-y.

24. E. A. Newberry, R. Bhandari, G. V. Minsavage, S. Timilsina, M. O. Jibrin, J. Kemble, E. J. Sikora, J. B. Jones, N. Potnis, Independent evolution with the gene flux originating from multiple Xanthomonas species explains genomic heterogeneity in Xanthomonas perforans. Appl. Environ. Microbiol. (2019), doi:10.1128/AEM.00885-19.

25. D. M. Emms, S. Kelly, OrthoFinder: solving fundamental biases in whole genome comparisons dramatically improves orthogroup inference accuracy. Genome Biol. (2015), doi:10.1186/s13059-015-0721-2.

26. P. Jones, D. Binns, H. Y. Chang, M. Fraser, W. Li, C. McAnulla, H. McWilliam, J. Maslen, A. Mitchell, G. Nuka, S. Pesseat, A. F. Quinn, A. Sangrador-Vegas, M. Scheremetjew, S. Y. Yong, R. Lopez, S. Hunter, InterProScan 5: Genome-scale protein function classification. Bioinformatics (2014), doi:10.1093/bioinformatics/btu031.

27. S. N. Gardner, T. Slezak, B. G. Hall, kSNP3.0: SNP detection and phylogenetic analysis of genomes without genome alignment or reference genome. Bioinformatics (2015), doi:10.1093/bioinformatics/btv271.

28. M. Pagel, A. Meade, BayesTraits. 2005 IEEE Comput. Syst. Bioinforma. Conf. Work. Poster Abstr. (2005), doi:10.1109/CSBW.2005.110.

29. K. H. Choi, H. P. Schweizer, mini-Tn7 insertion in bacteria with single attTn7 sites: Example Pseudomonas aeruginosa. Nat. Protoc. (2006), doi:10.1038/nprot.2006.24.

30. S.-W. Han, C.-J. Park, S.-W. Lee, P. C. Ronald, An efficient method for visualization and growth of fluorescent Xanthomonas oryzae pv. oryzae in planta. BMC Microbiol. 8, 164 (2008).

31. L. Salichos, A. Rokas, Inferring ancient divergences requires genes with strong phylogenetic signals. Nature (2013), doi:10.1038/nature12130.

32. K. Katoh, D. M. Standley, MAFFT multiple sequence alignment software version 7: improvements in performance and usability. Mol. Biol. Evol. (2013), doi:10.1093/molbev/mst010.

33. S. Capella-Gutiérrez, J. M. Silla-Martínez, T. Gabaldón, trimAl: A tool for automated alignment trimming in large-scale phylogenetic analyses. Bioinformatics (2009), doi:10.1093/bioinformatics/btp348.

34. L. T. Nguyen, H. A. Schmidt, A. Von Haeseler, B. Q. Minh, IQ-TREE: A fast and effective stochastic algorithm for estimating maximum-likelihood phylogenies. Mol. Biol. Evol. (2015), doi:10.1093/molbev/msu300.

35. A. Stamatakis, RAxML version 8: A tool for phylogenetic analysis and post-analysis of large phylogenies. Bioinformatics (2014), doi:10.1093/bioinformatics/btu033.

36. P. Siguier, ISfinder: the reference centre for bacterial insertion sequences. Nucleic Acids Res. (2006), doi:10.1093/nar/gkj014.

37. B. Boussau, L. Guéguen, M. Gouy, A mixture model and a Hidden Markov Model to simultaneously detect recombination breakpoints and reconstruct phylogenies. Evol. Bioinforma. (2009).

38. H. Shimodaira, An approximately unbiased test of phylogenetic tree selection. Syst. Biol. (2002), doi:10.1080/10635150290069913.

39. J. Huerta-Cepas, F. Serra, P. Bork, ETE 3: Reconstruction, Analysis, and Visualization of Phylogenomic Data. Mol. Biol. Evol. (2016), doi:10.1093/molbev/msw046.

40. M. J. Sullivan, N. K. Petty, S. A. Beatson, Easyfig: A genome comparison visualizer. Bioinformatics (2011), doi:10.1093/bioinformatics/btr039.

