## Supplementary figures and images for "Repeated gain and loss of a single gene modulates the evolution of vascular pathogen lifestyles"

### Supplemental Figure 1

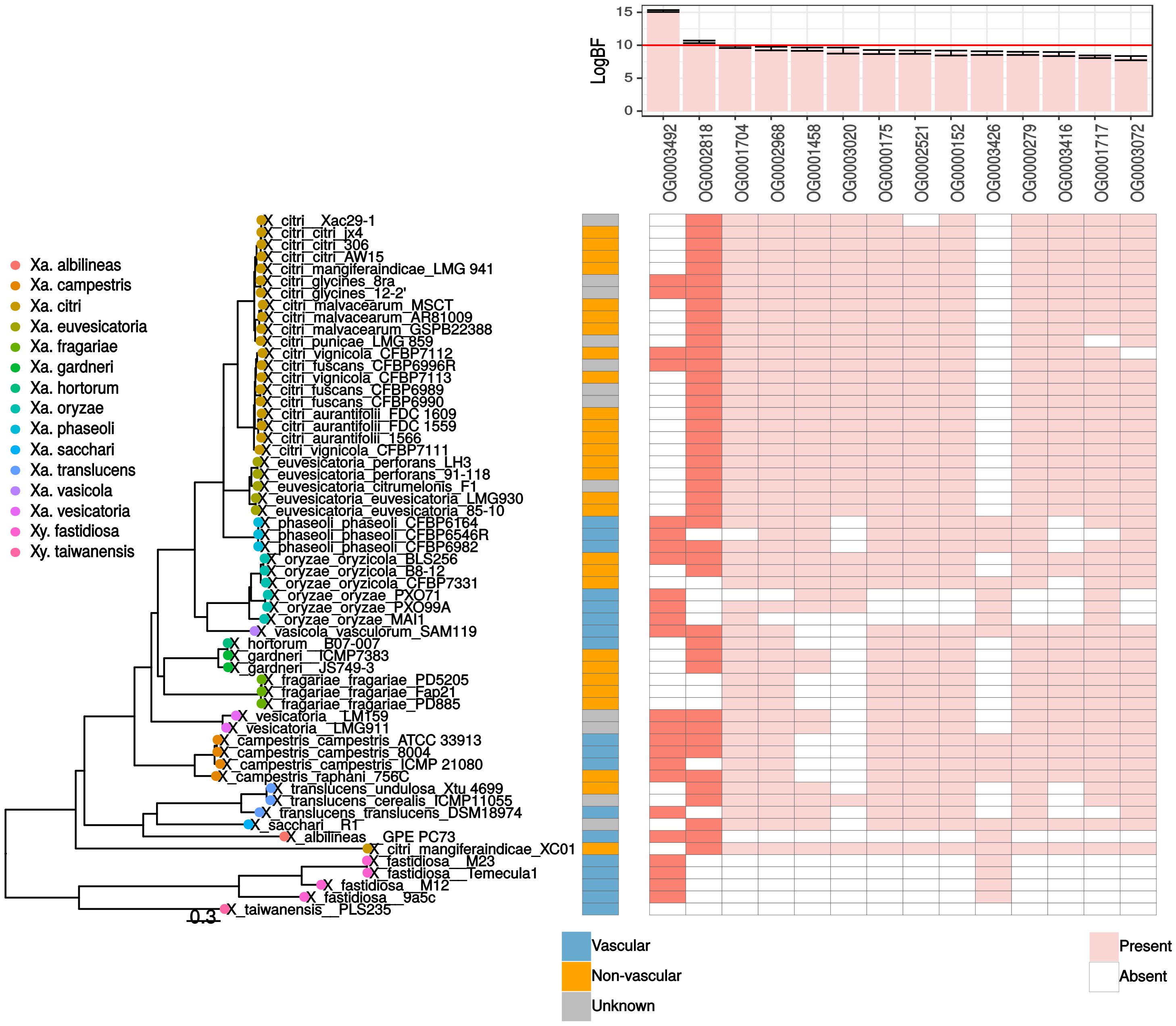

### Supplemental Figure 2

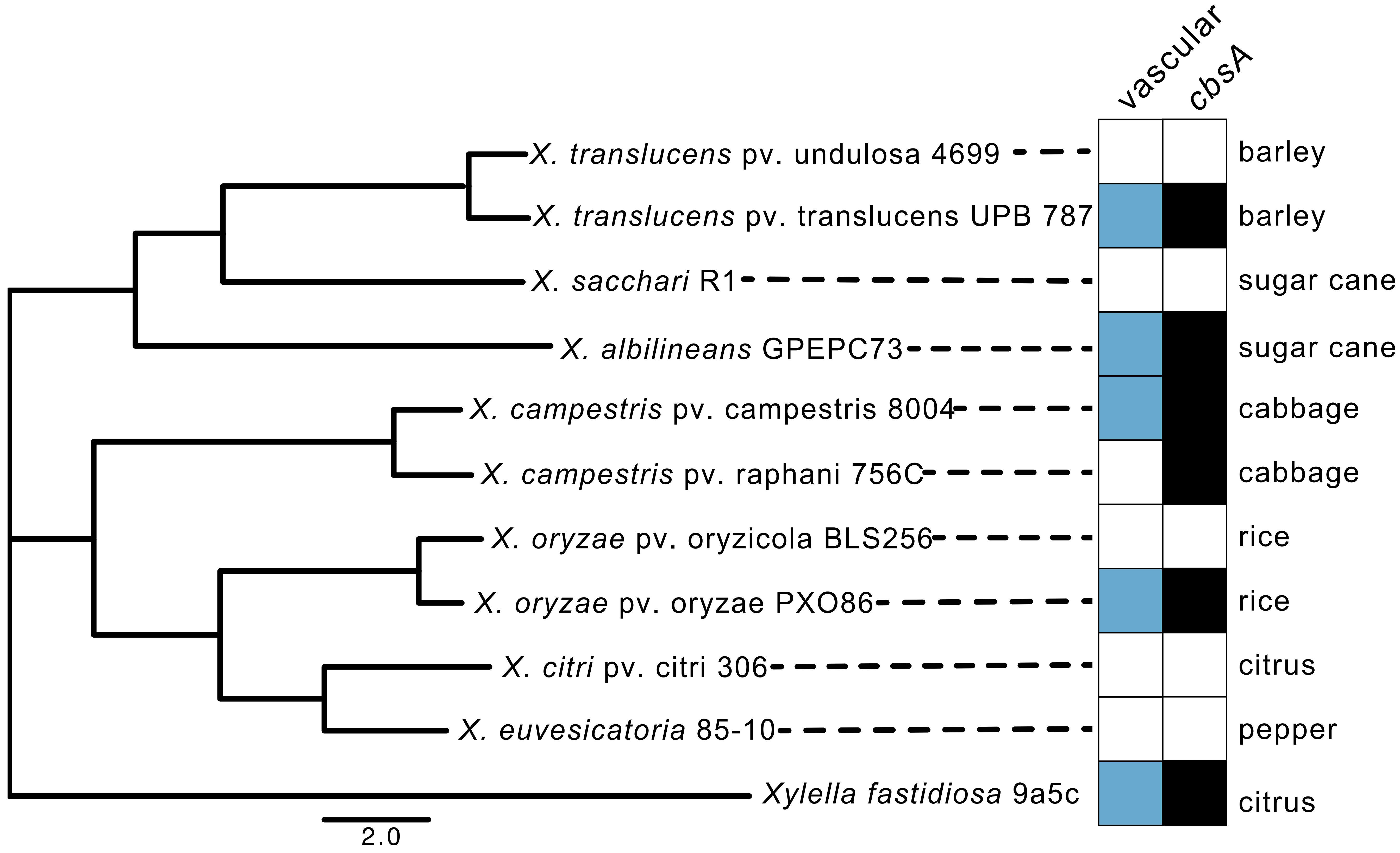

### Supplemental Figure 3

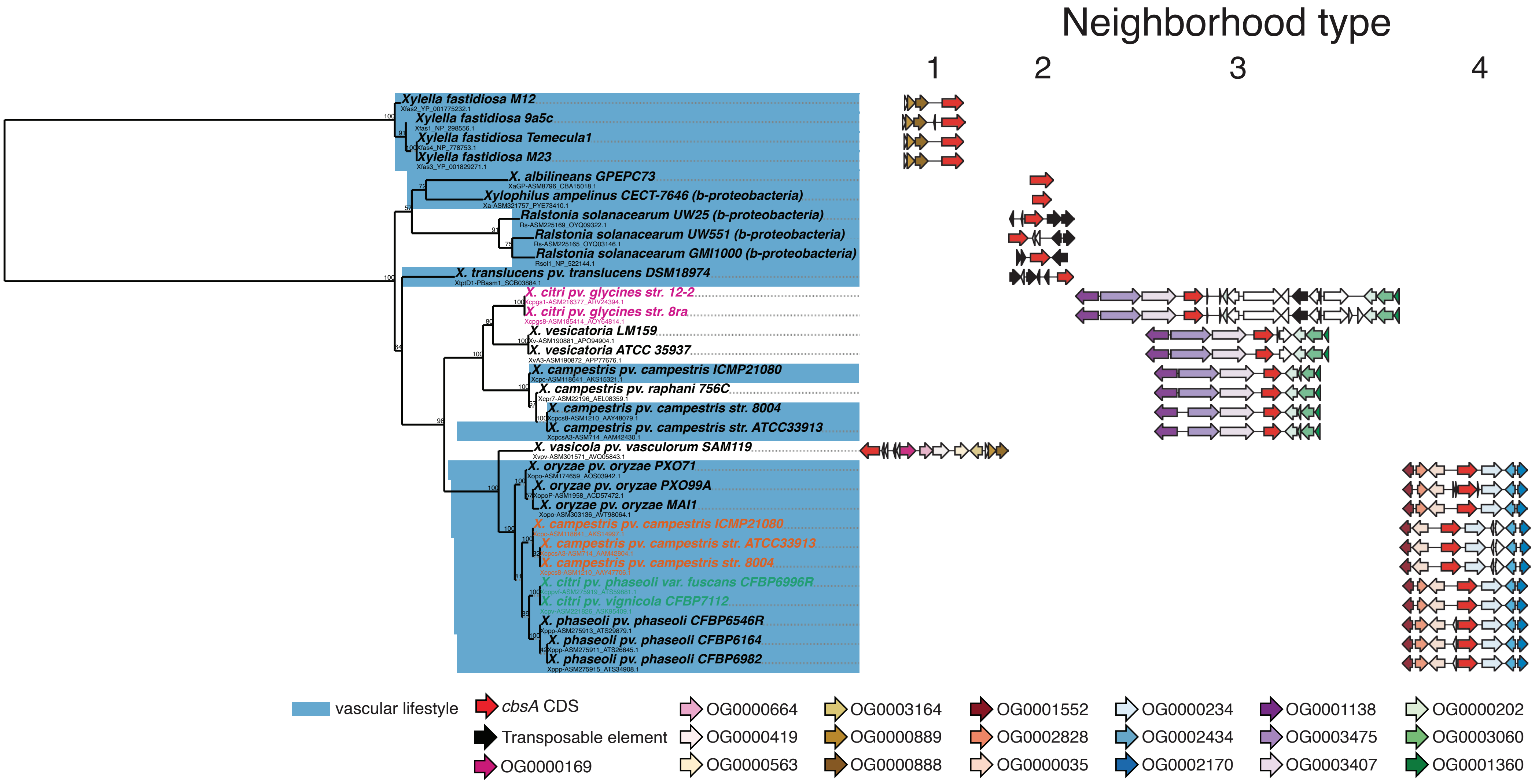

### Supplemental Figure 4

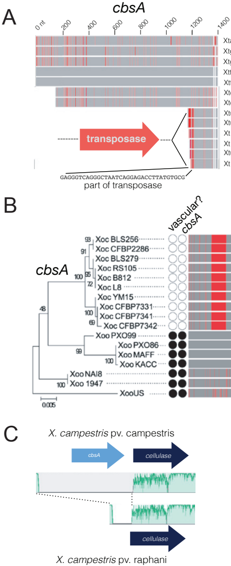

### Supplemental Figure 5

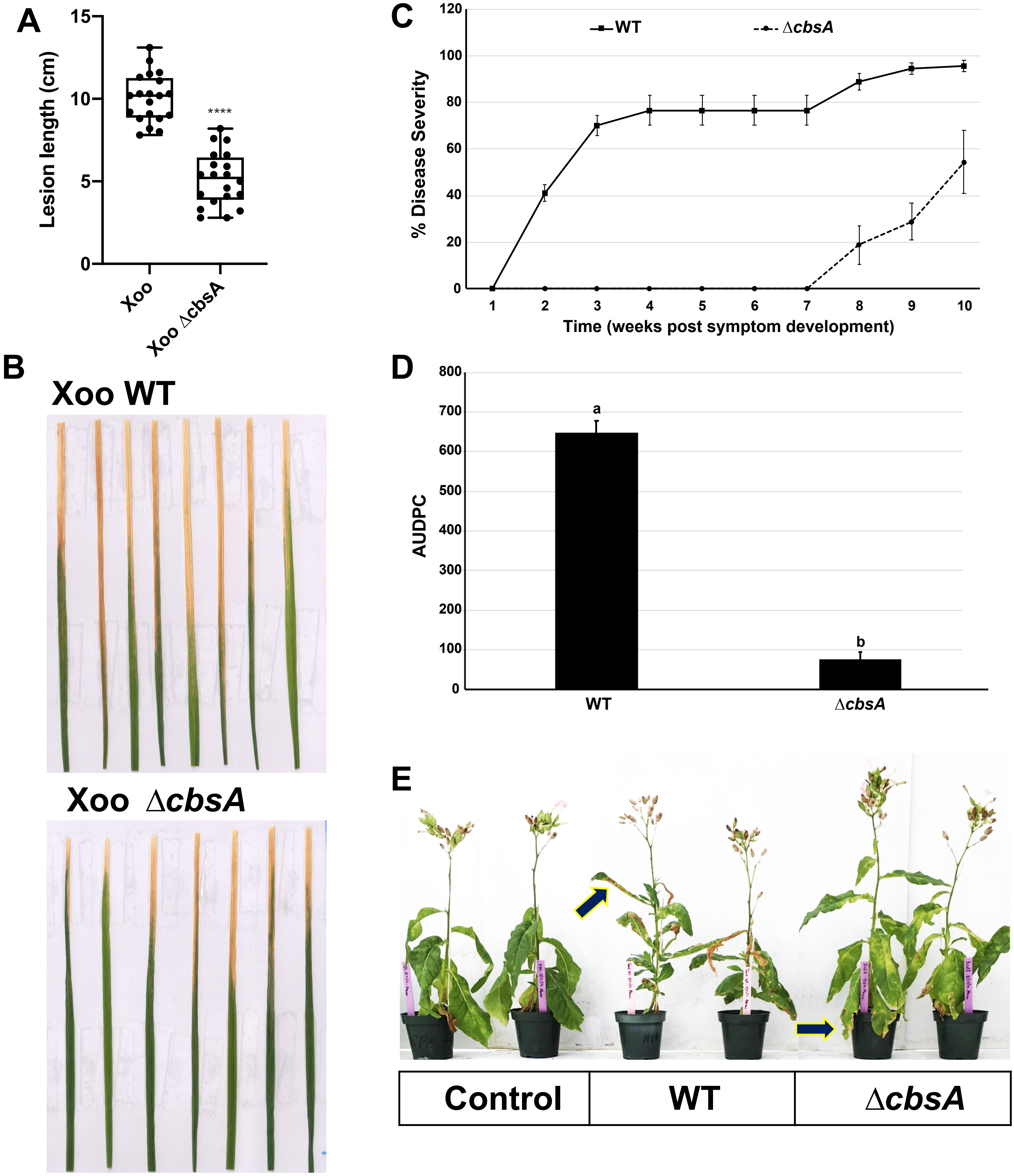

### Supplemental Figure 6

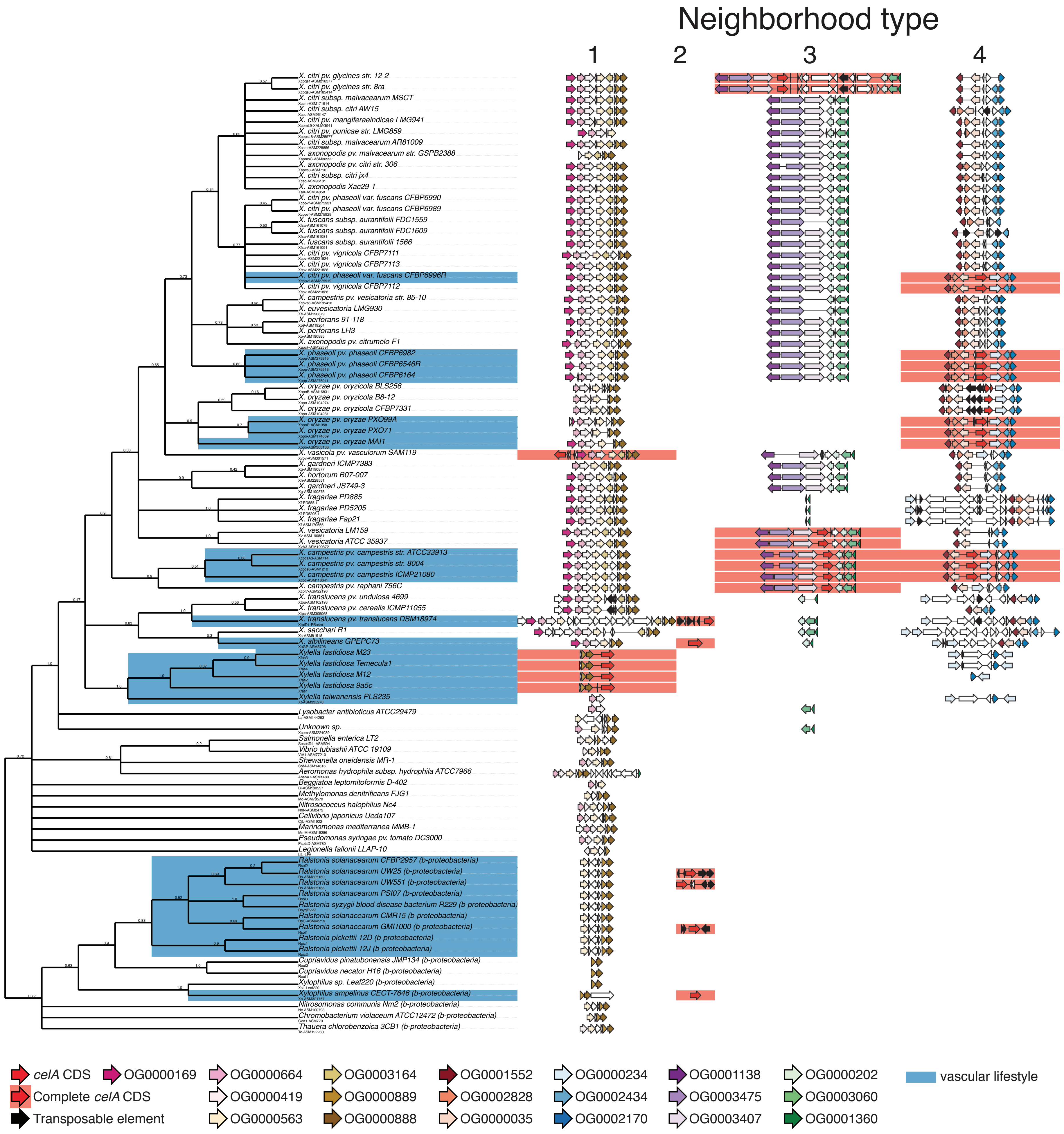

### Supplemental figure 9

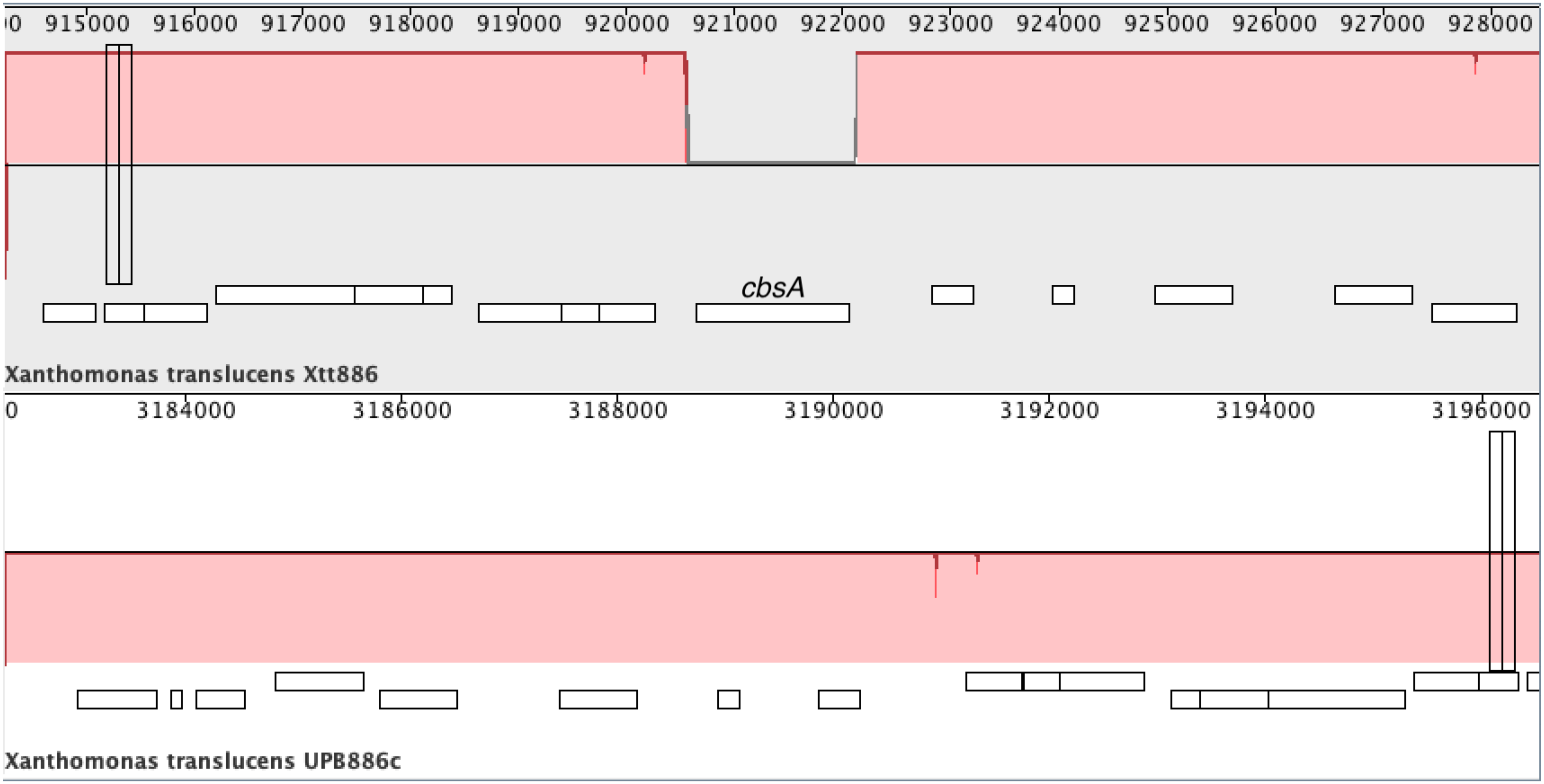
