## Supplemental Figure 7 for "Repeated gain and loss of a single gene modulates the evolution of vascular pathogen lifestyles"

a) OG0003189 (carboxylesterase)

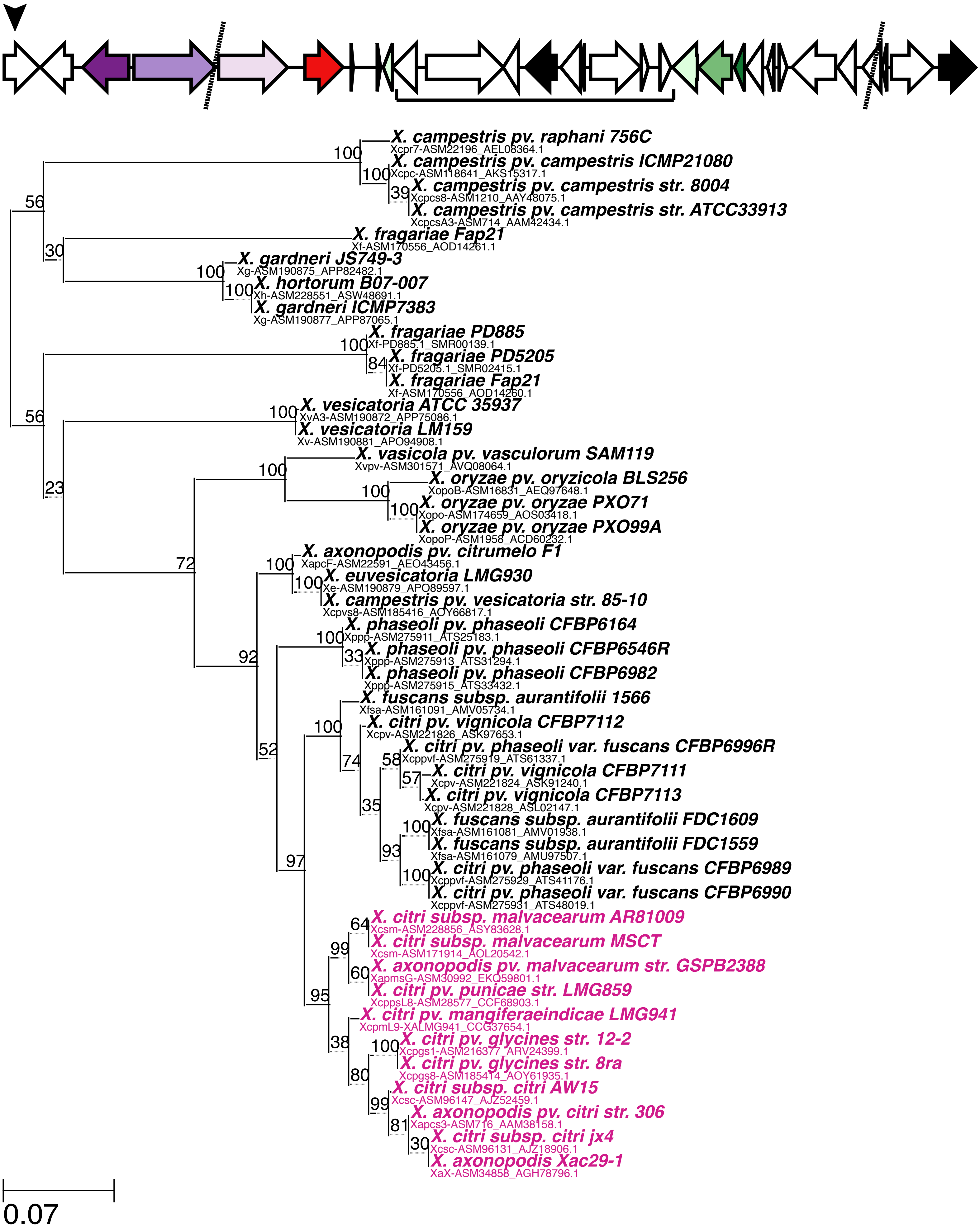

b) OG0003864 (function unknown)

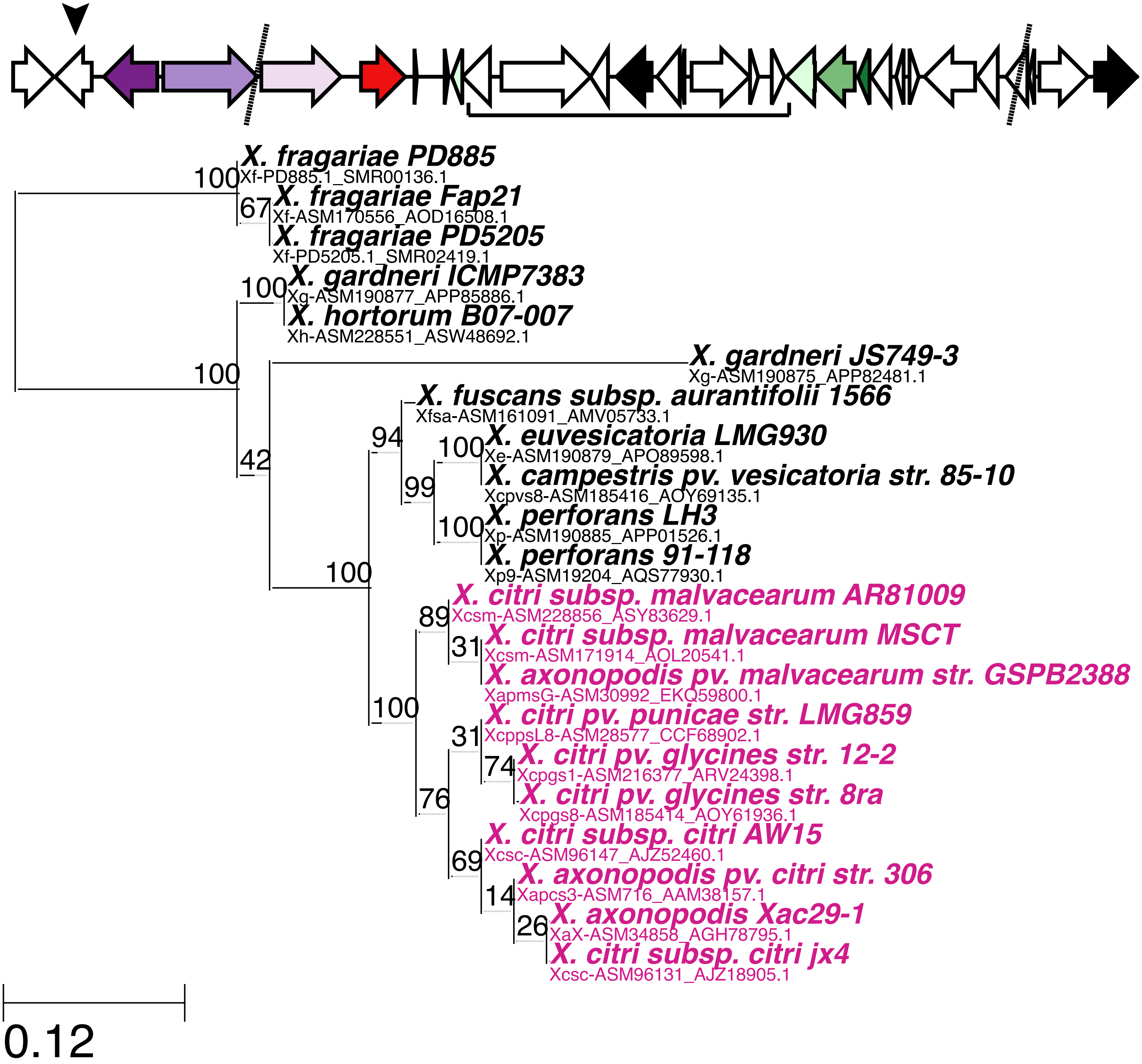

c) OG0001138 (alpha-glucosidase)

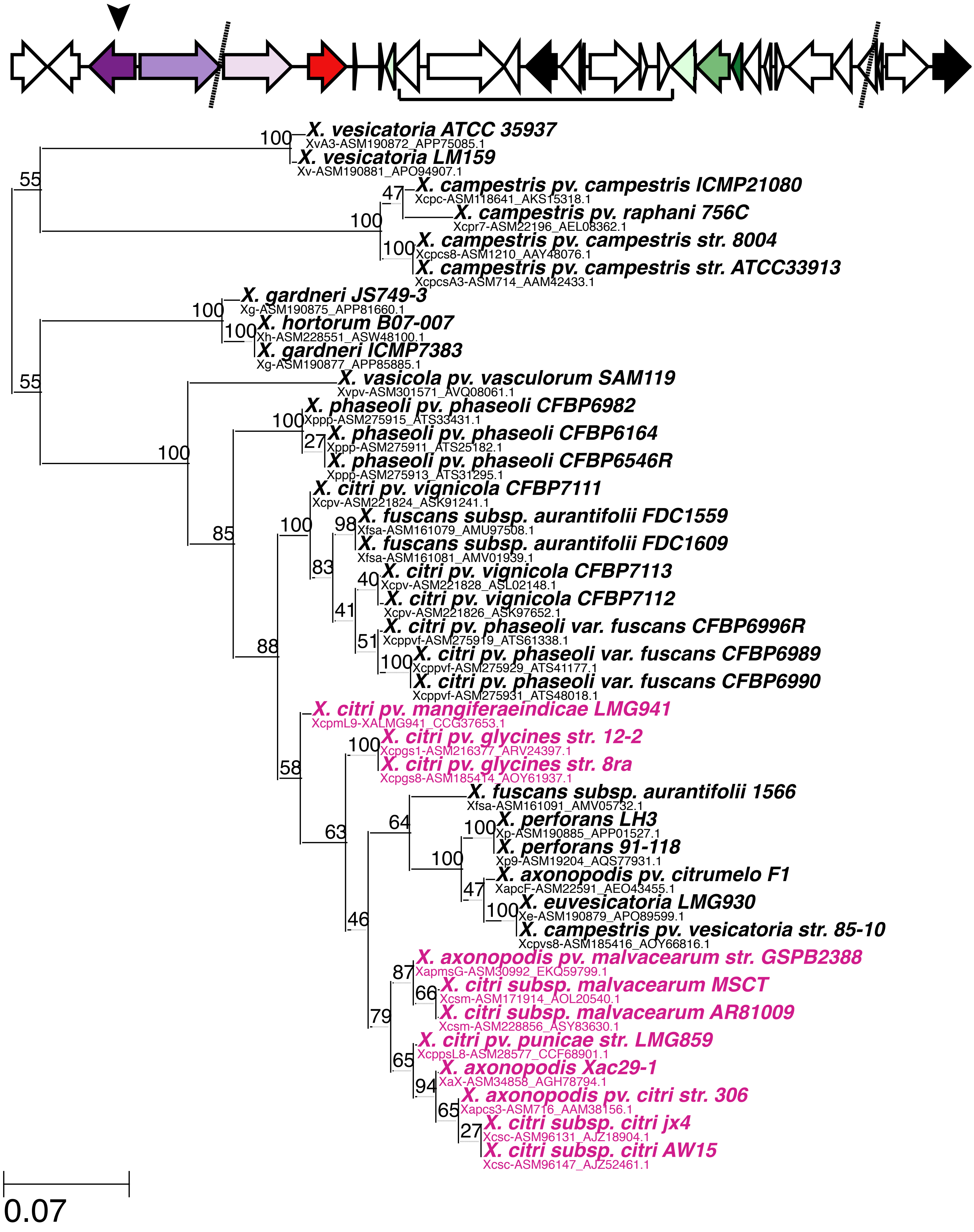

d) OG0003475 (hydrolase family 2, sugar binding)

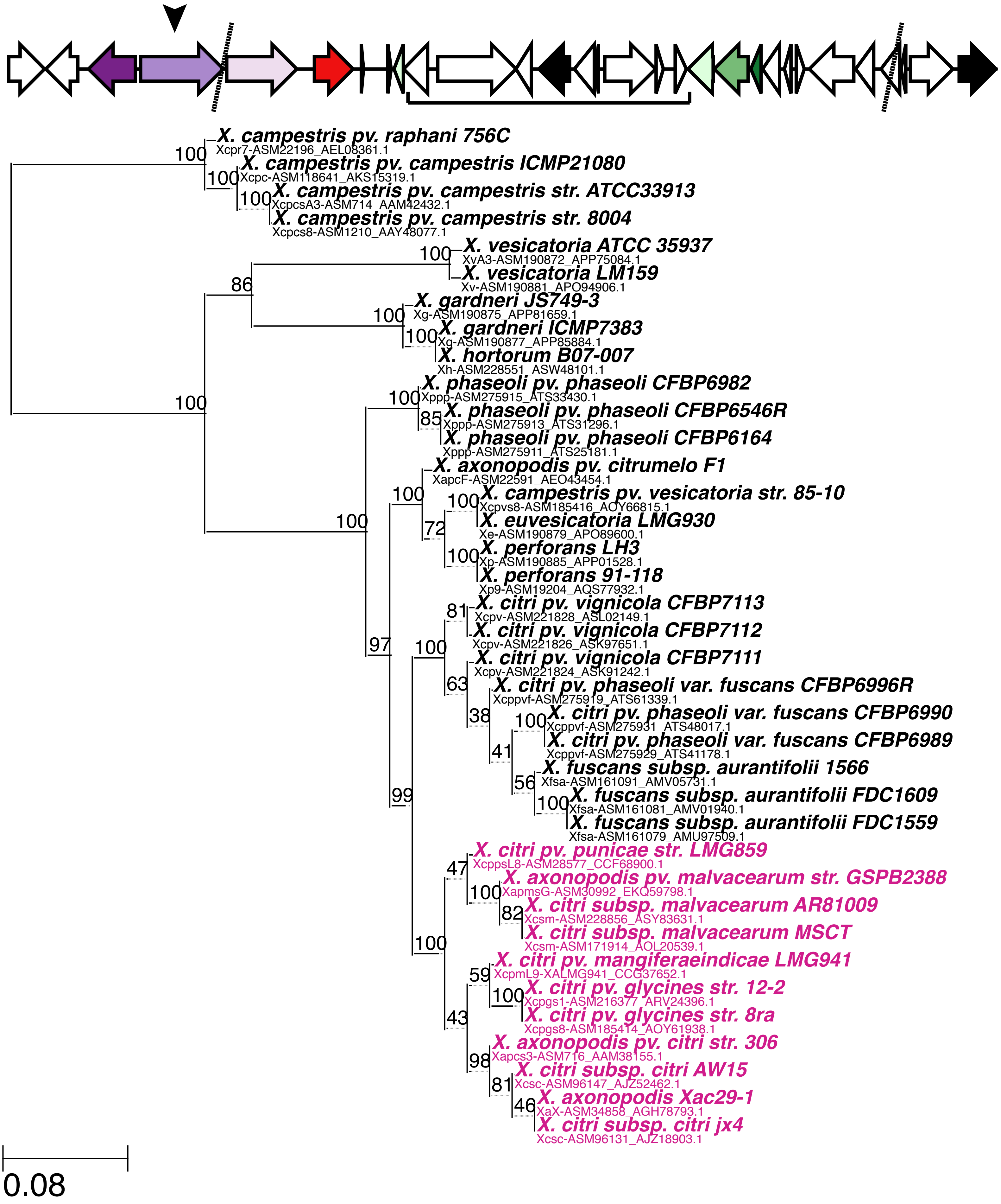

e) OG0003407 (tonB-dependent receptor)

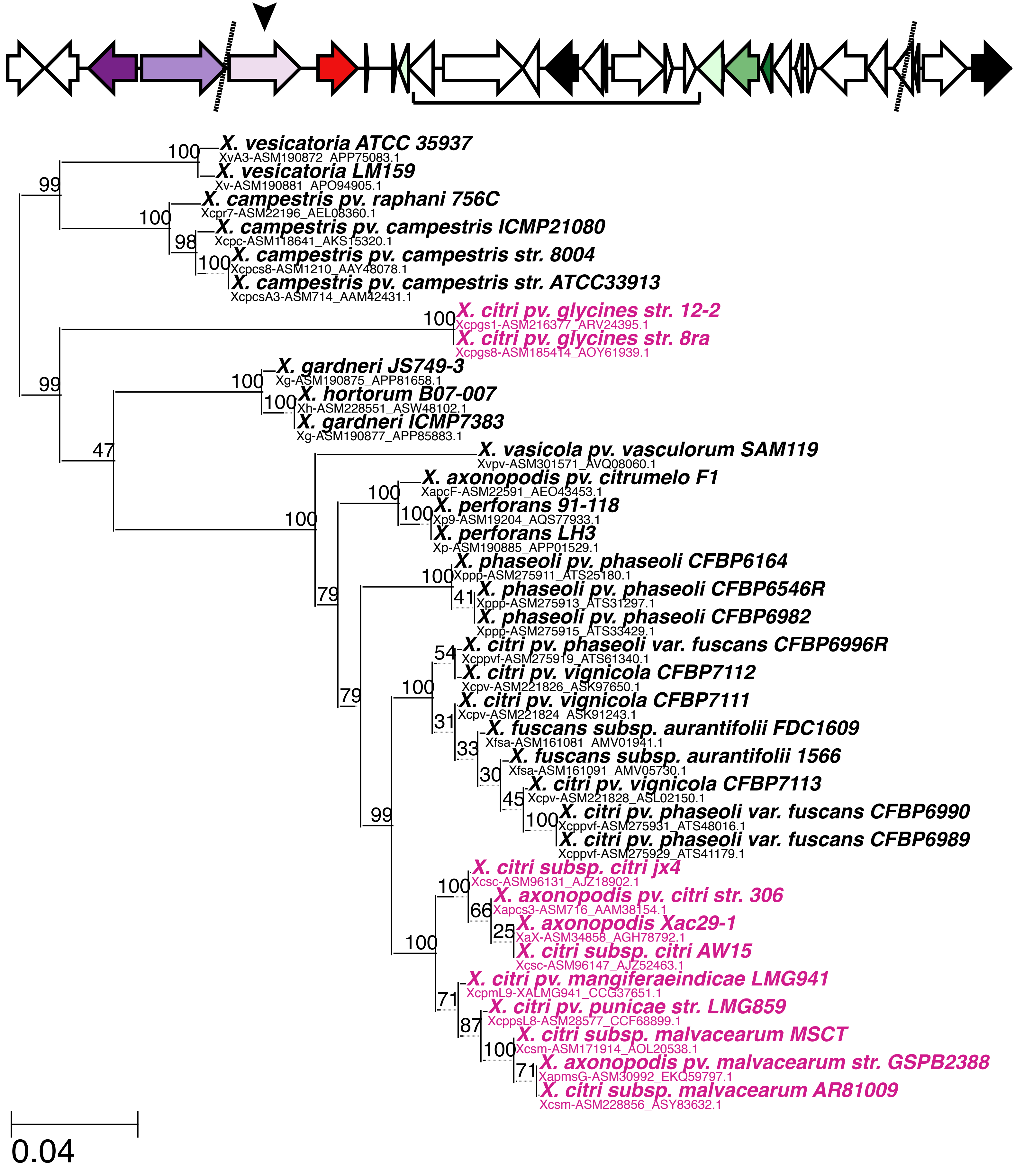

f) OG0006923 (cytochrome p450 domain-containing protein)

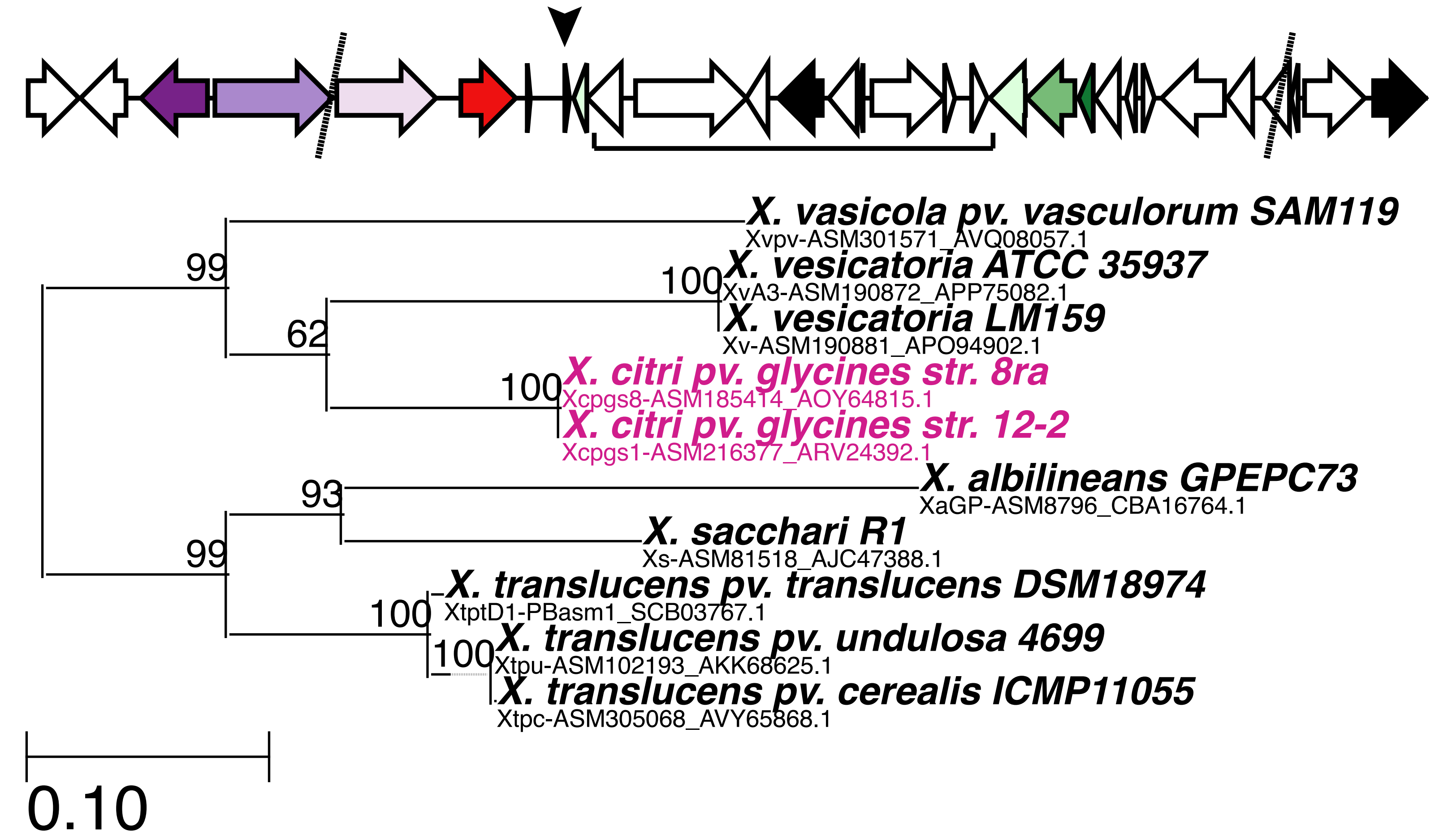

g) OG0000202 (transcription regulator, LacI family)

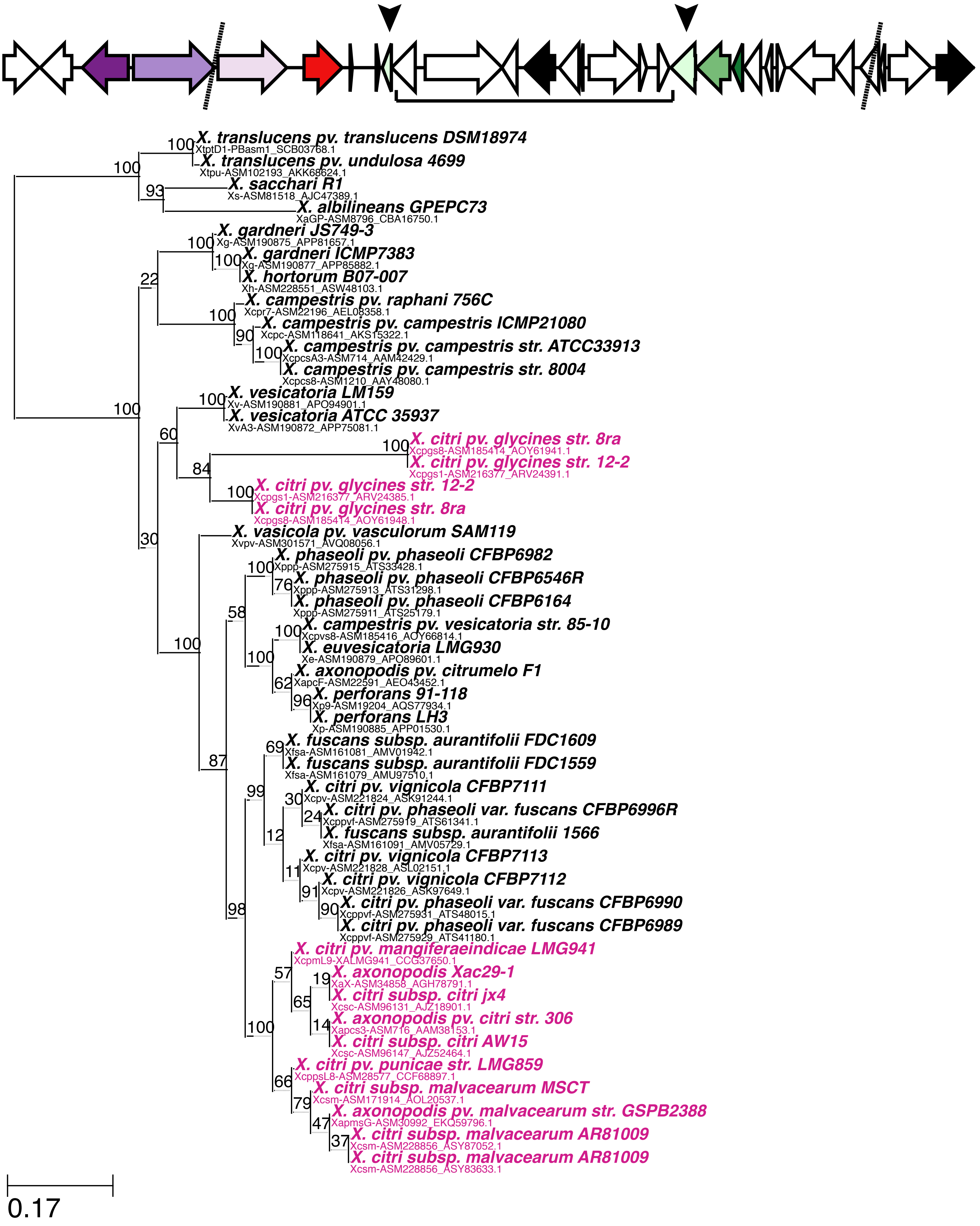

h) OG0012116 (response regulator)

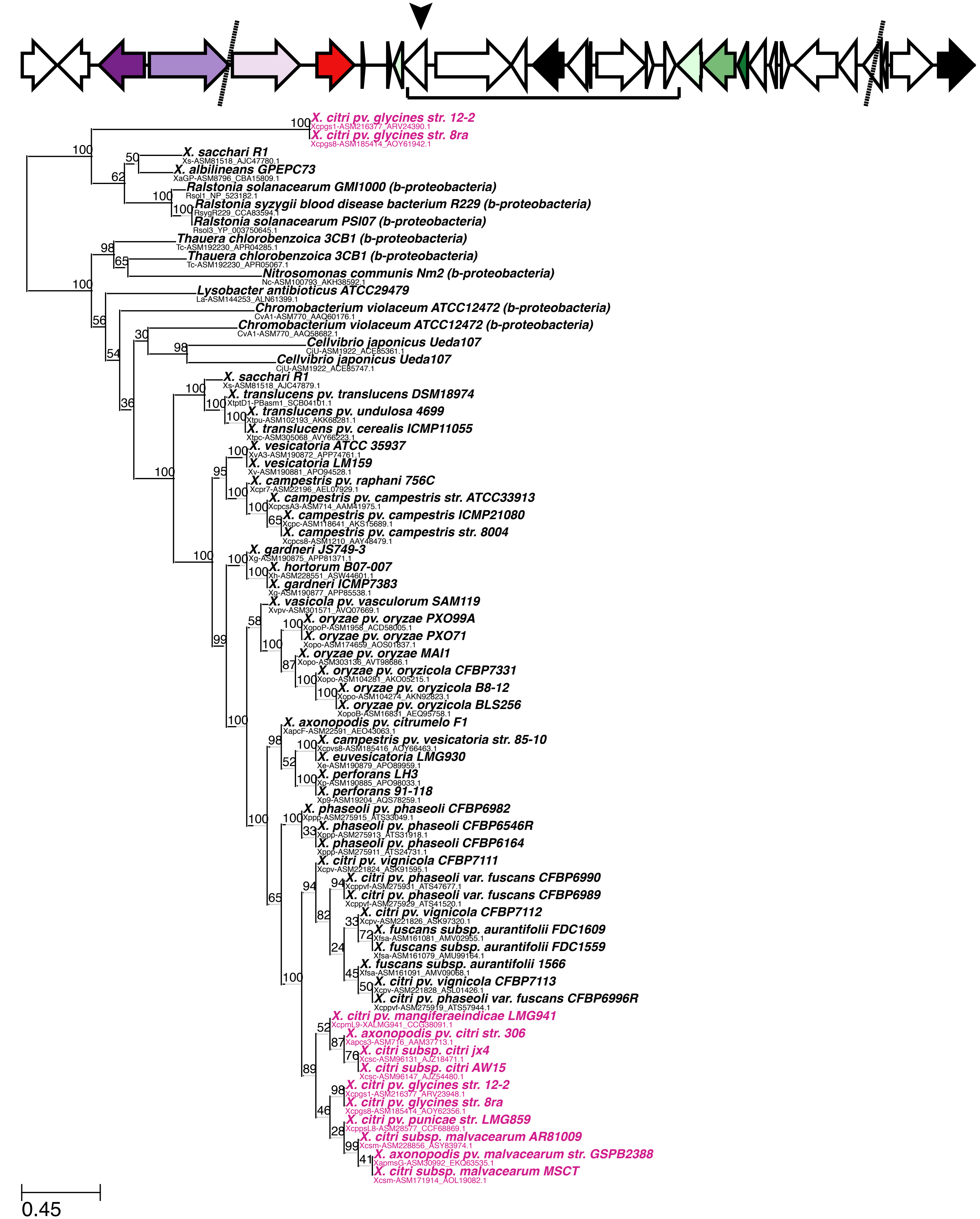

i) OG0006653 (histidine kinase)

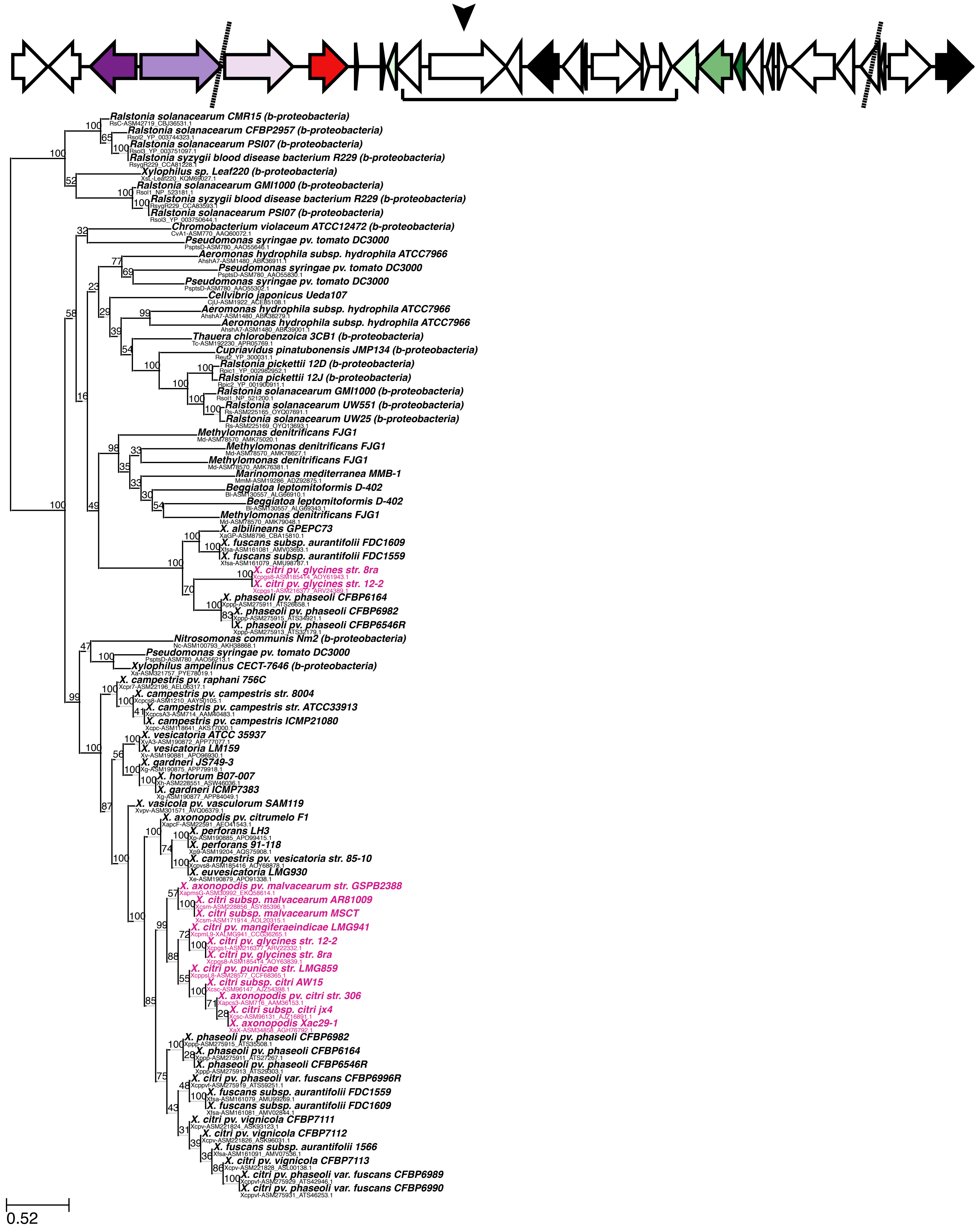

j) OG0004064 (response regulator)

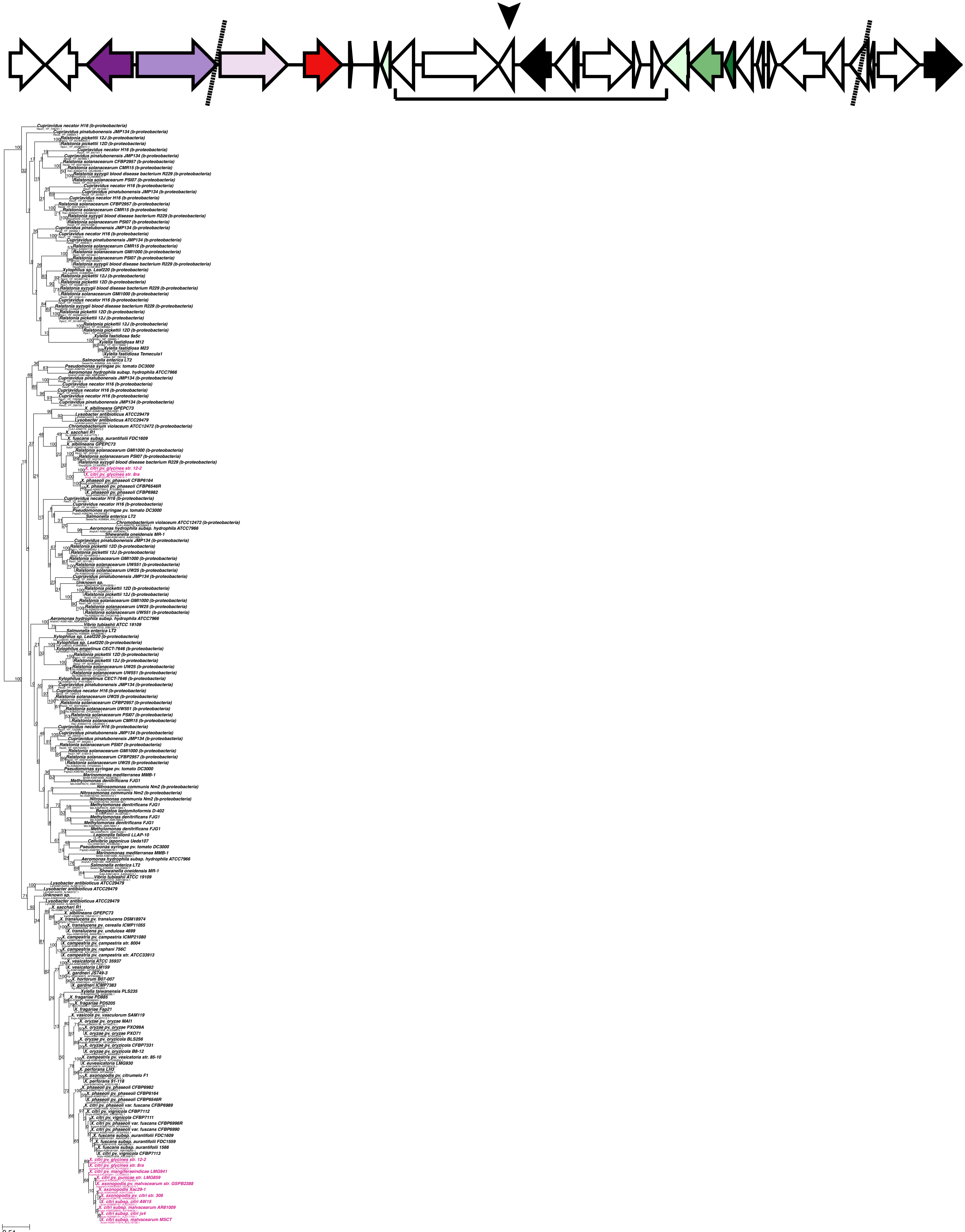

k) OG0005040 (function unknown)

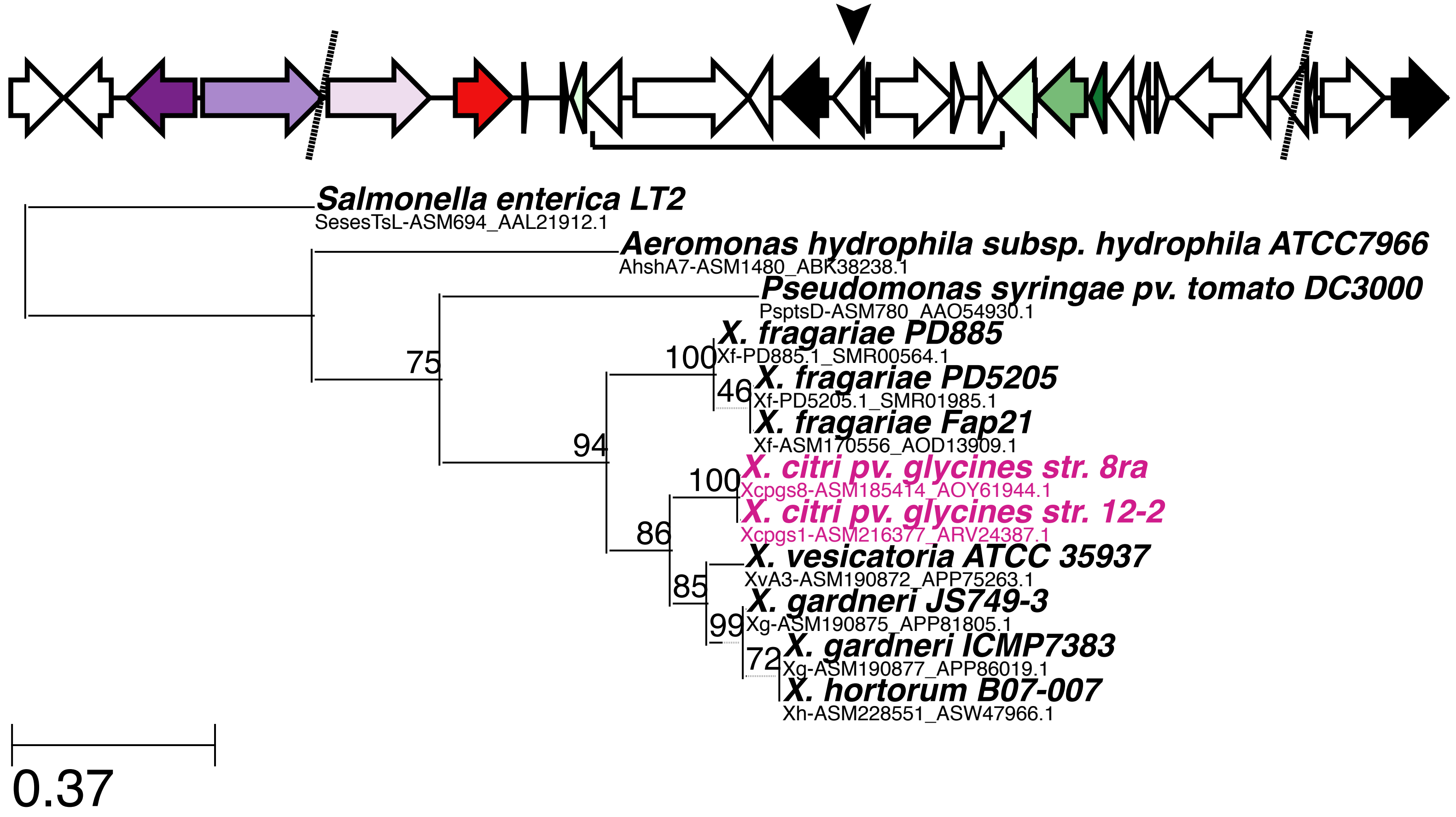

### I) OG0015184 (function unknown)

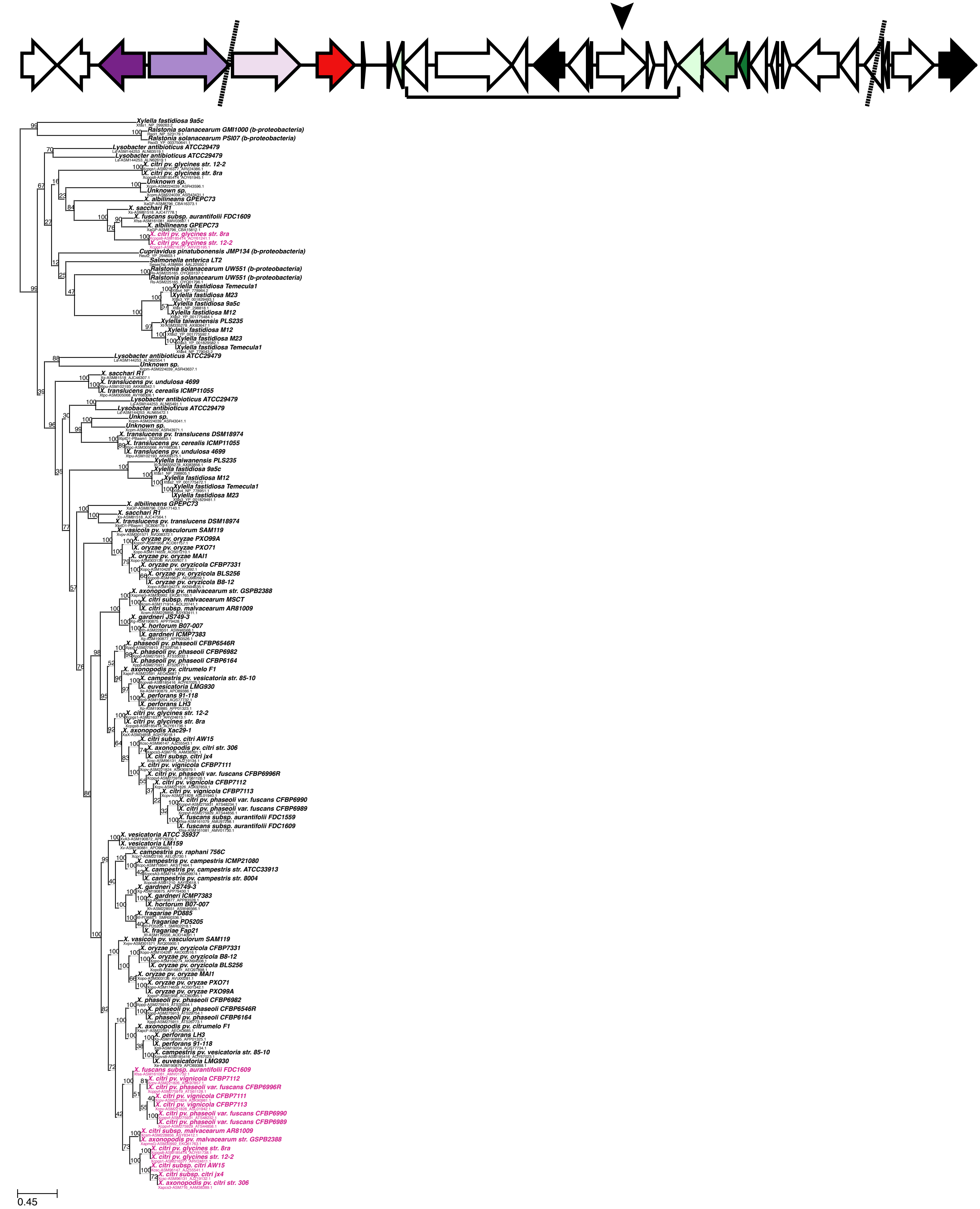

m) OG0004674 (function unknown)

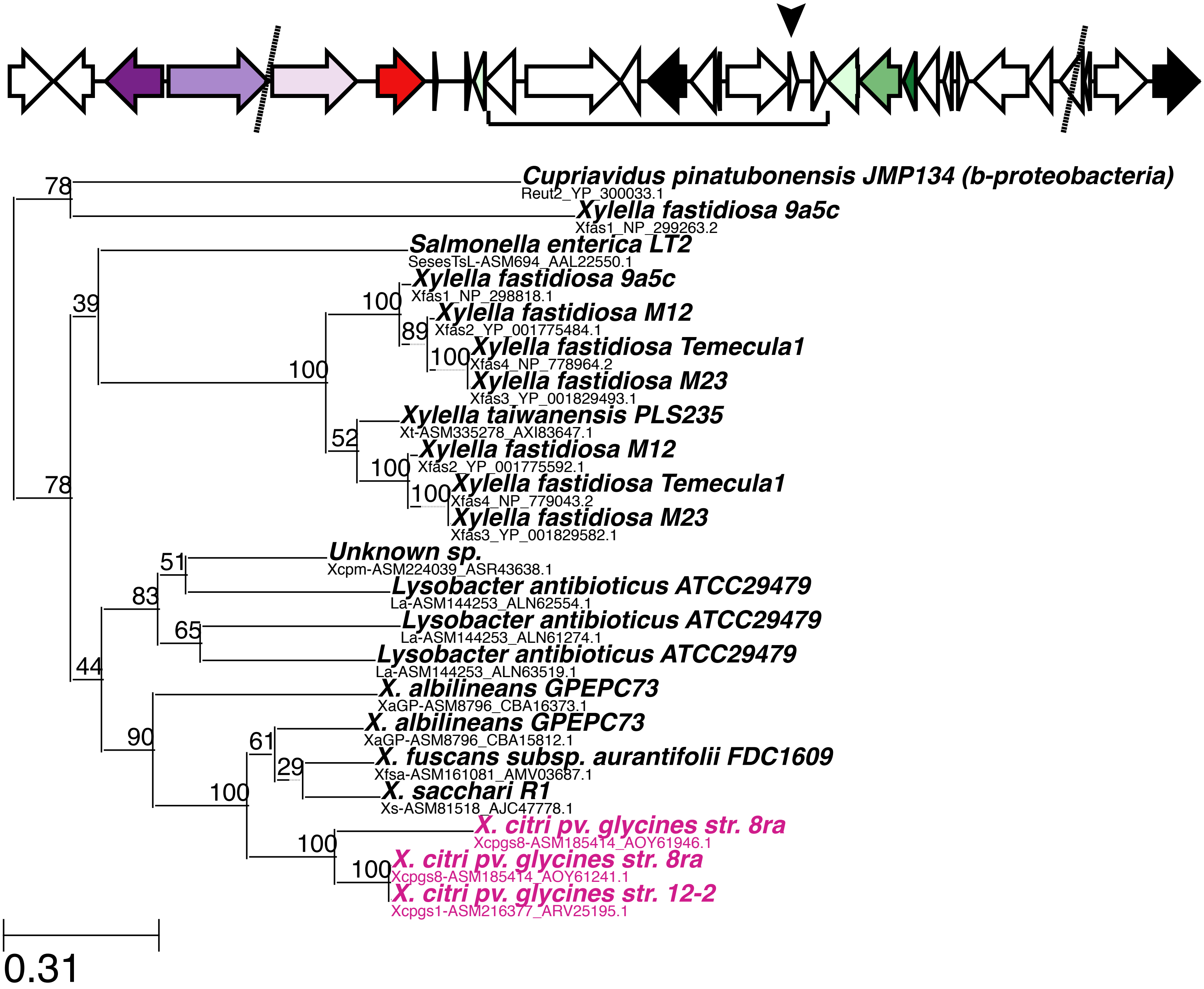

n) OG0005551 (CDP-diacylglycerol pyrophosphatase)

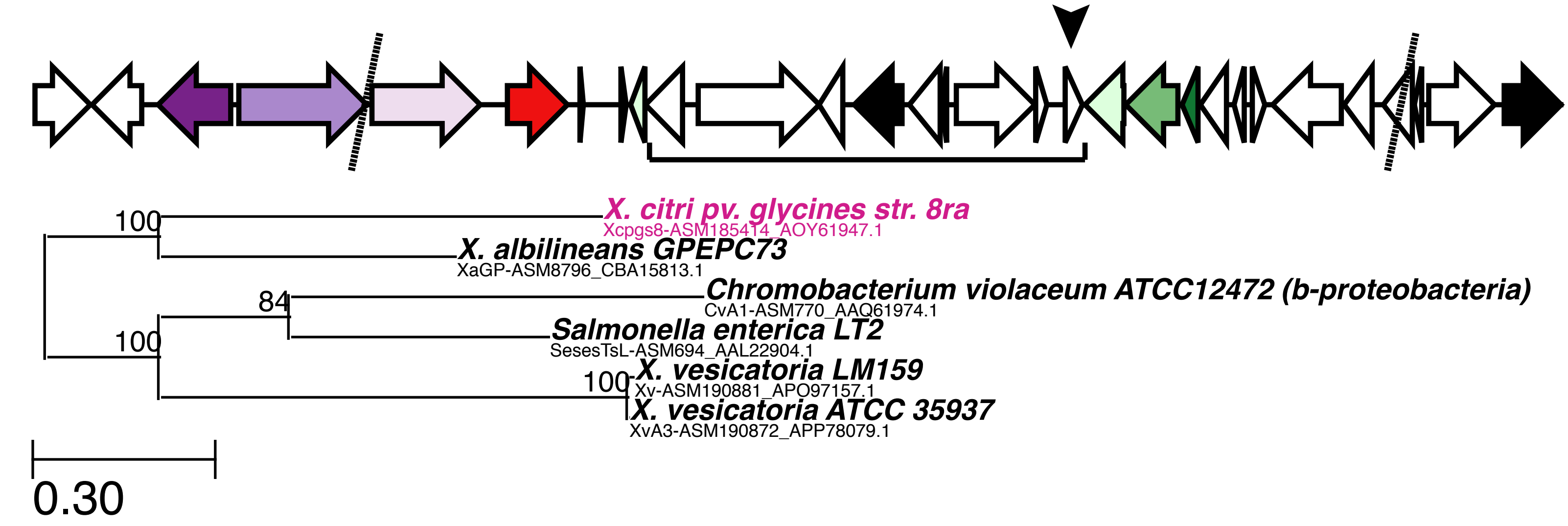

o) OG0003060 (peptidase m28)

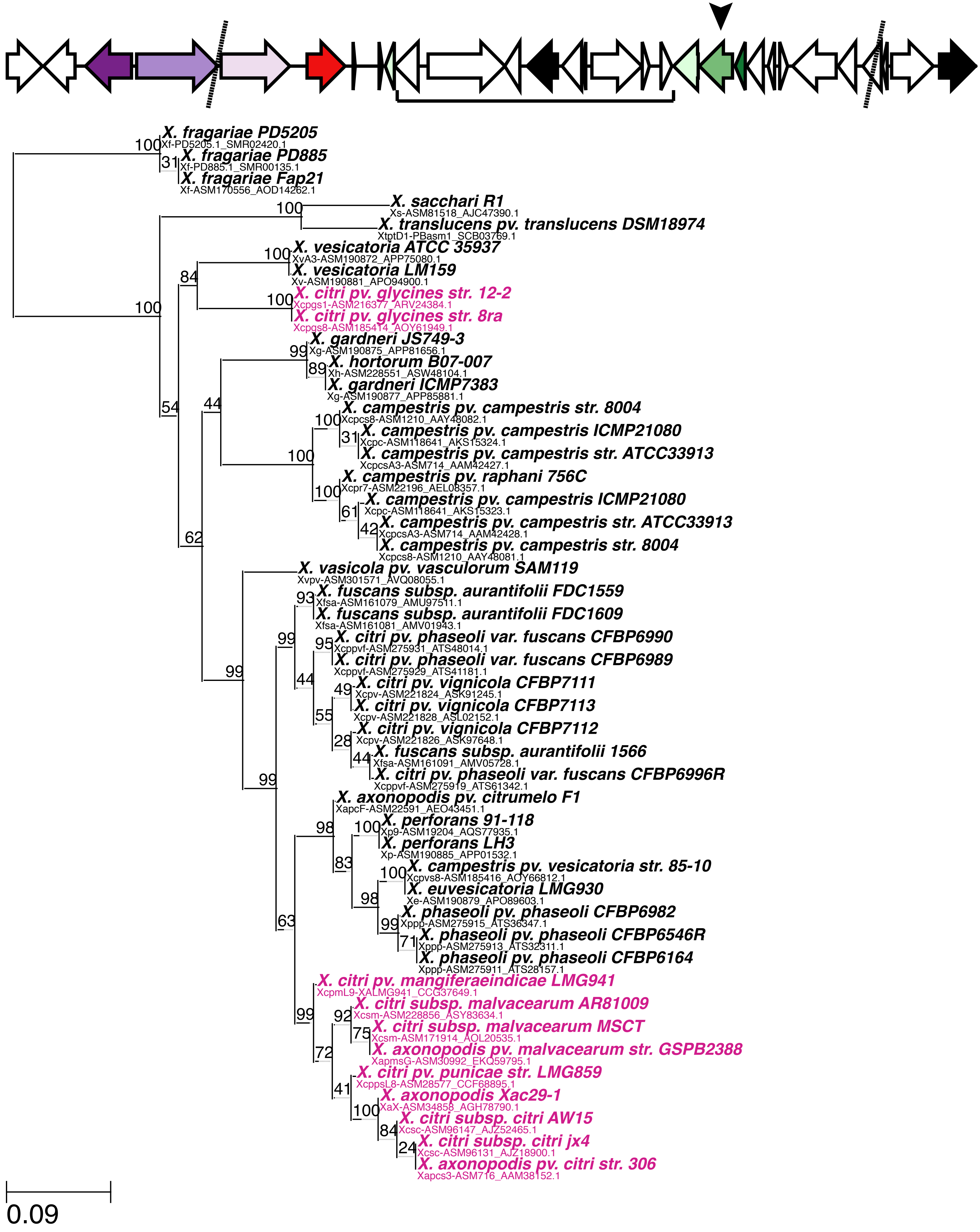

p) OG0001360 (large-conductance mechanosensitive ion channel)

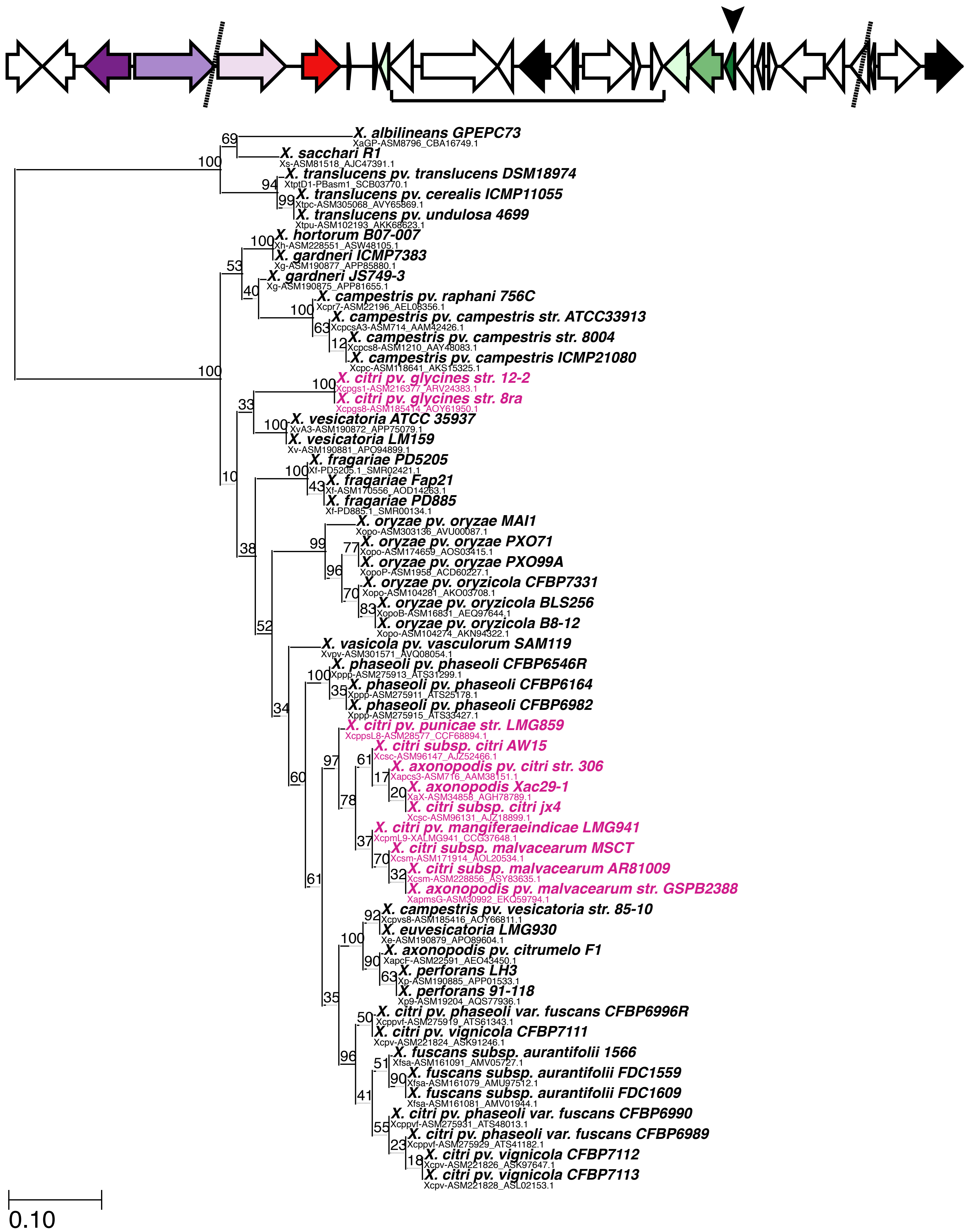

q) OG0000926 (fumarylacetoacetate hydrolase)

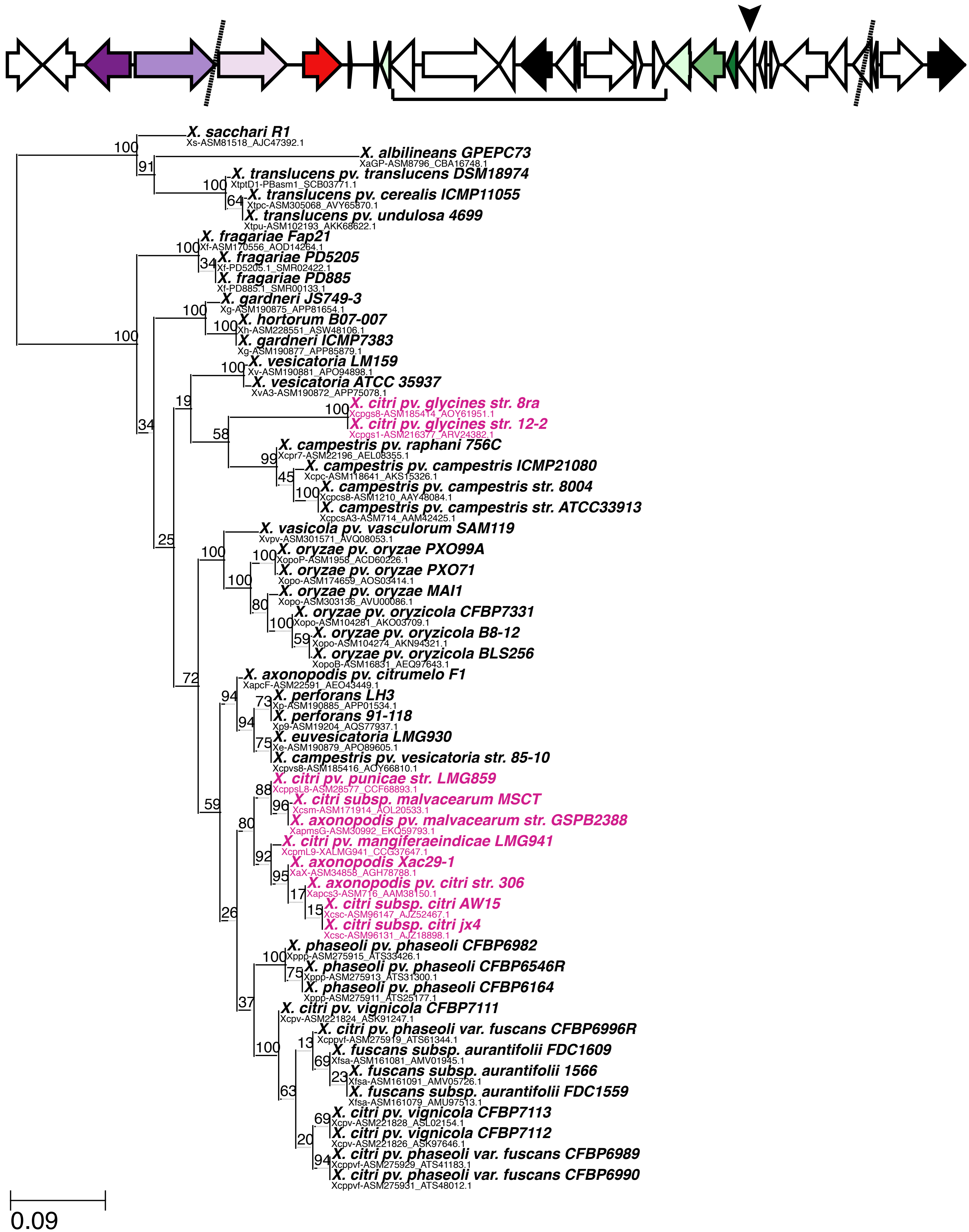

r) OG0002126 (function unknown)

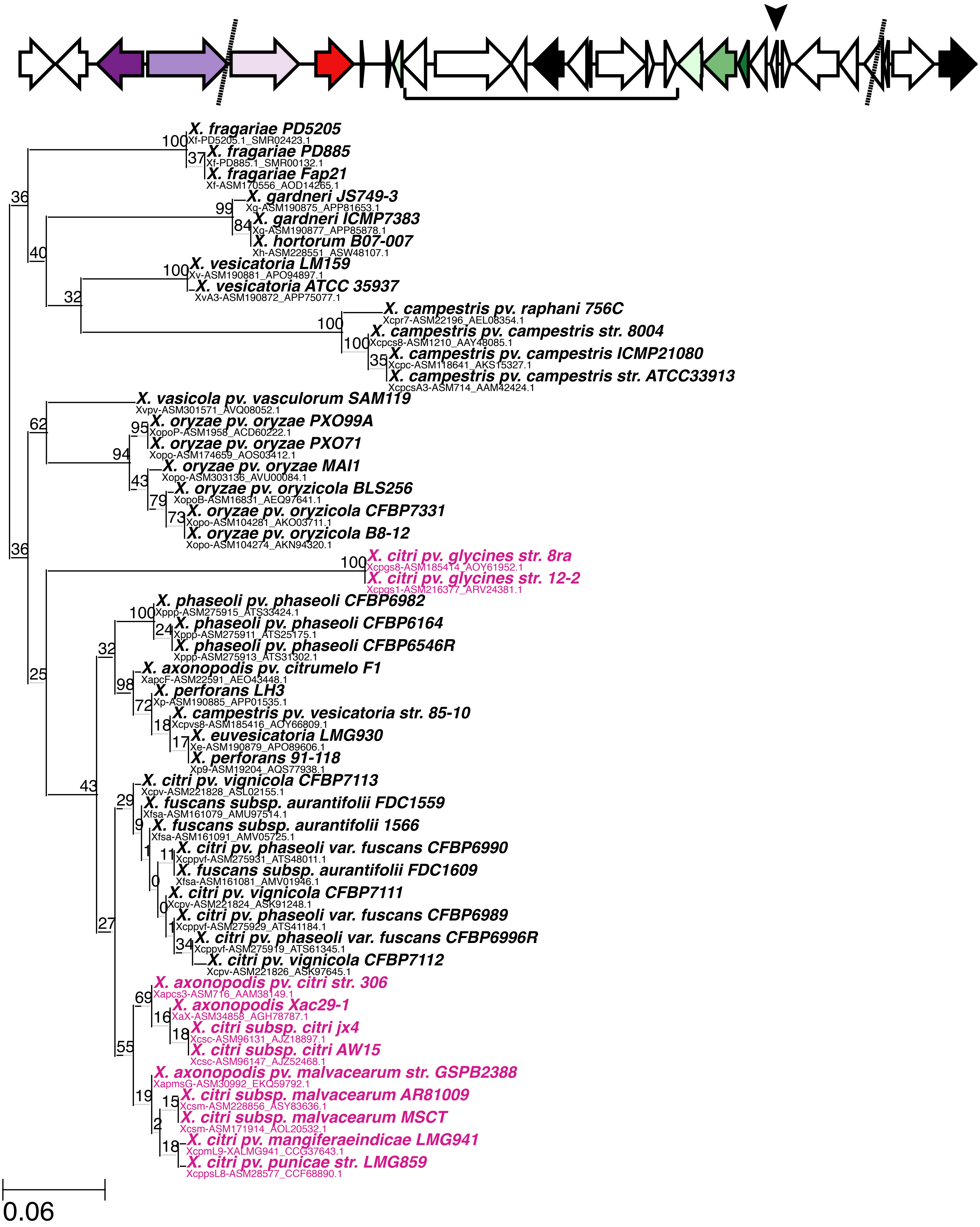

s) OG0001639 (Rieske [2Fe-2S] domain-containing protein)

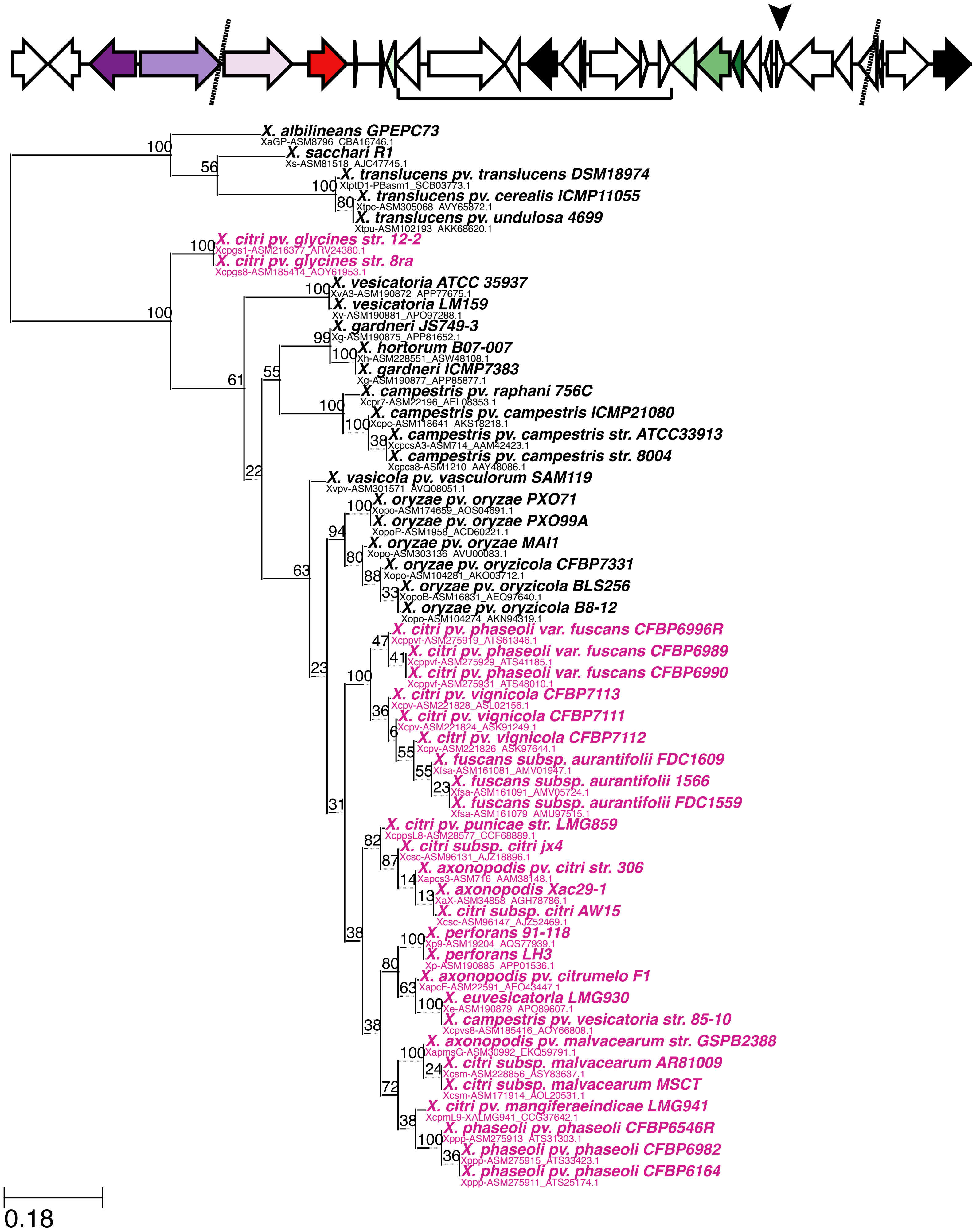

### t) OG0001483 (citrate transporter)

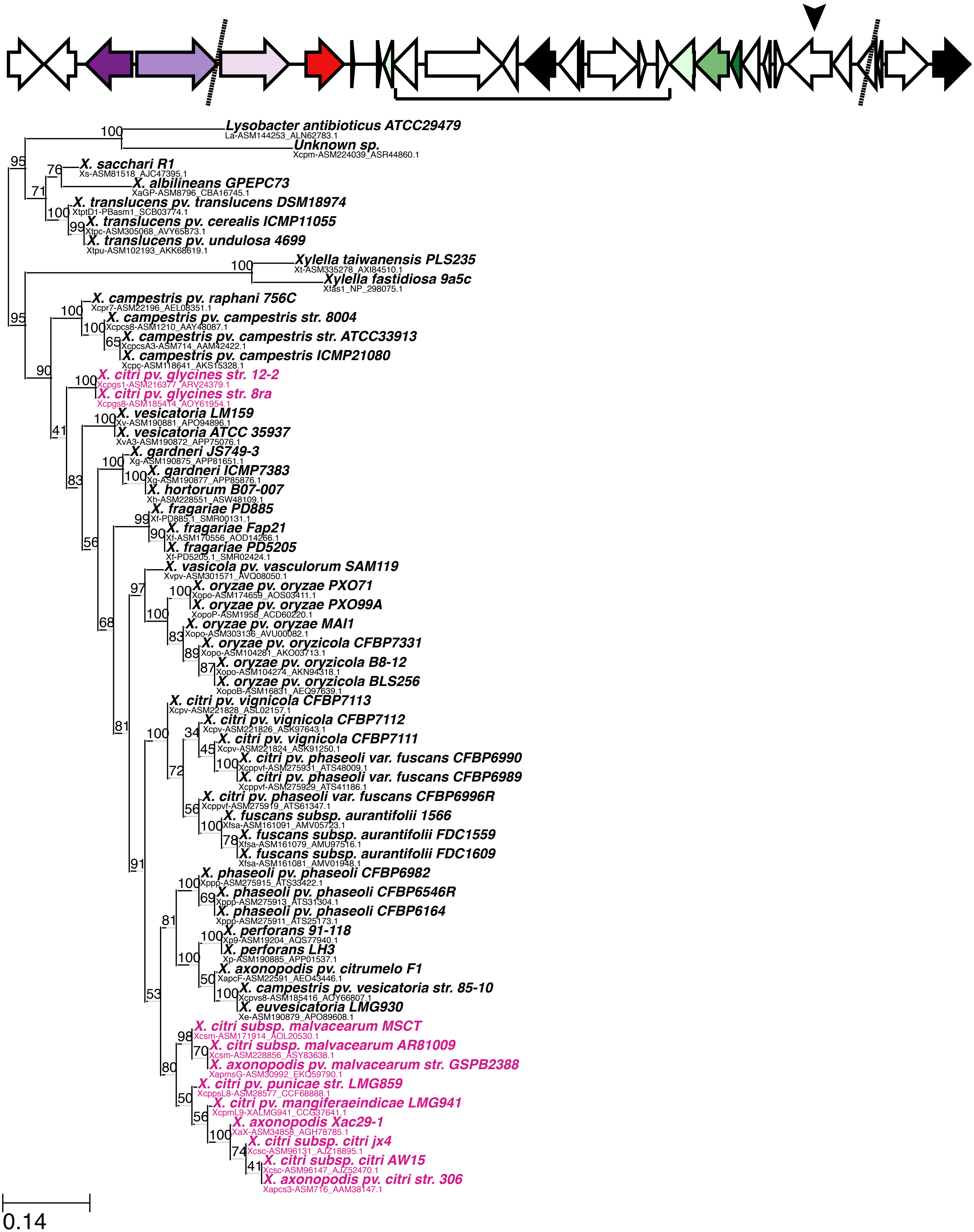

u) OG0001080 (putative methyltransferase)

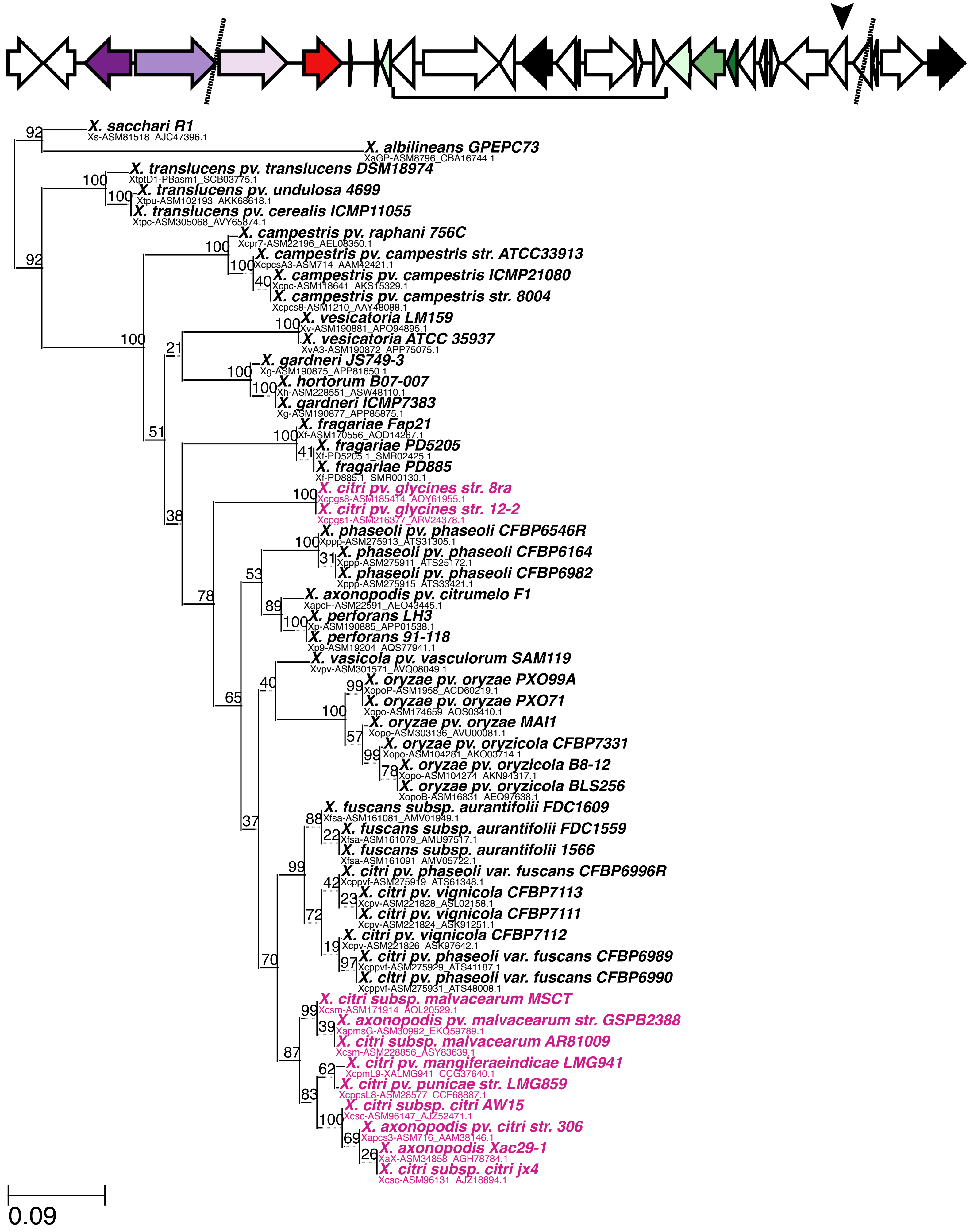

### v) OG0000801, partition 2 (thiazole biosynthesis protein thiG)

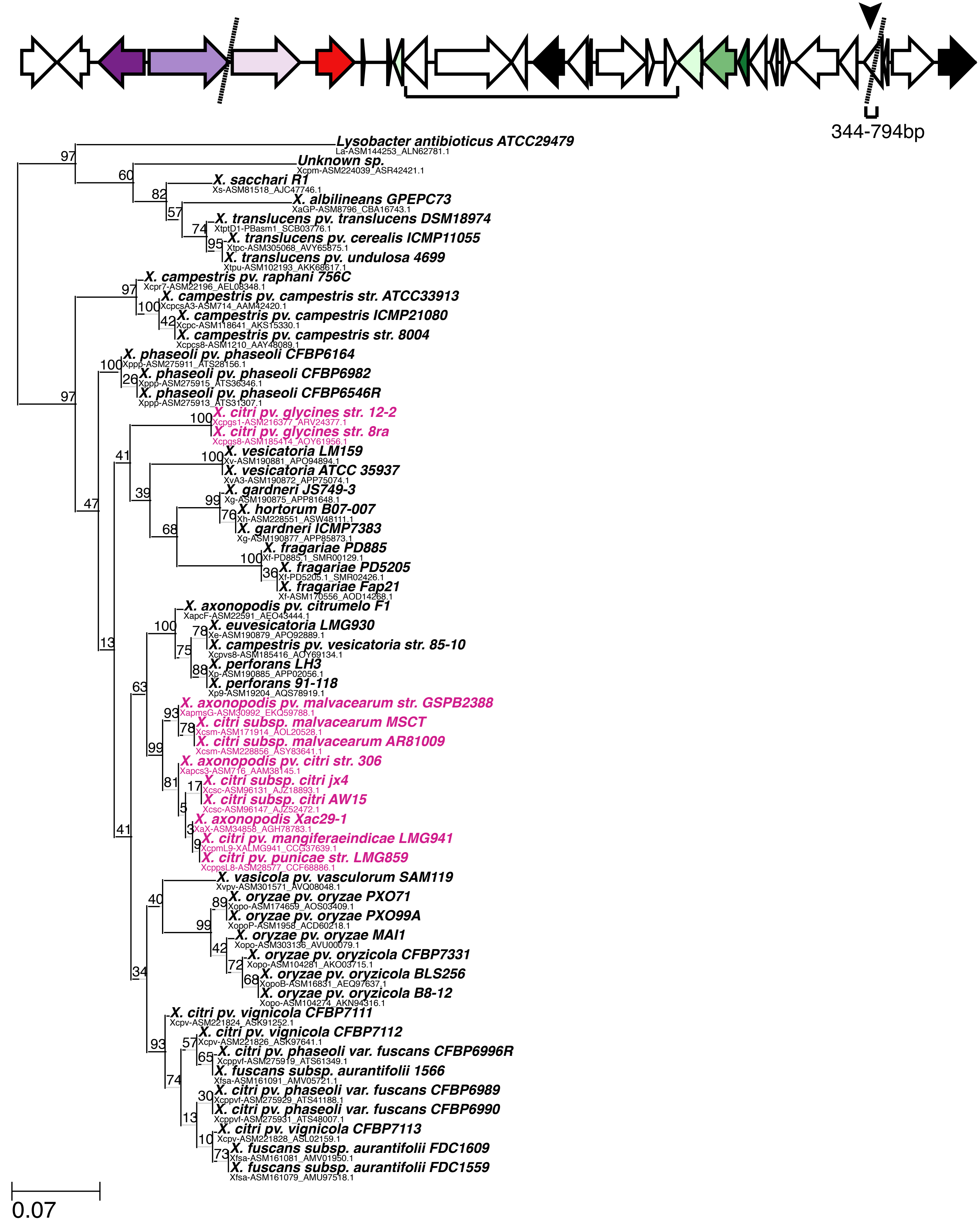

w) OG0000801, partition 1 (thiazole biosynthesis protein thiG)

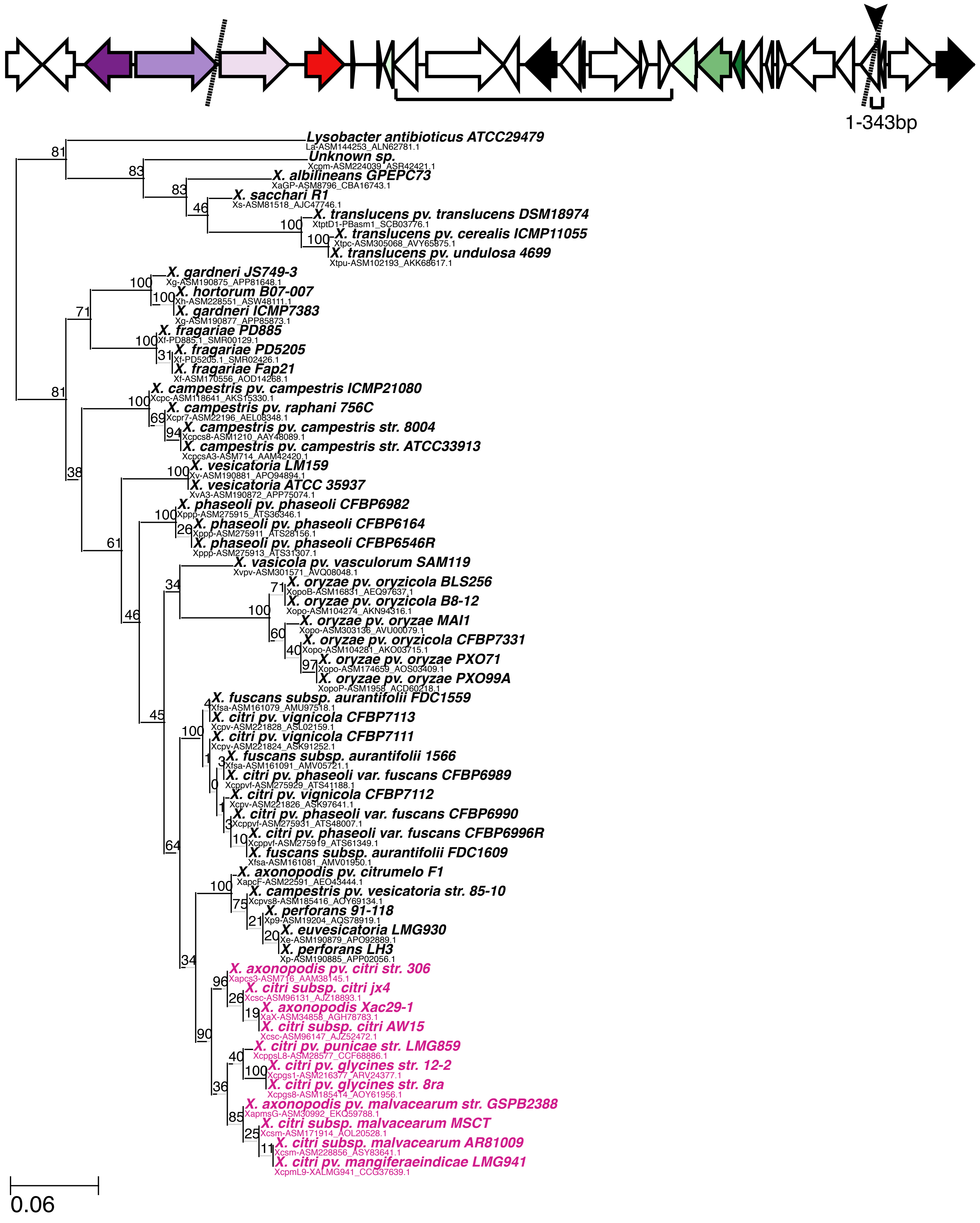

x) OG0001155 (thiazole biosynthesis protein thiS)

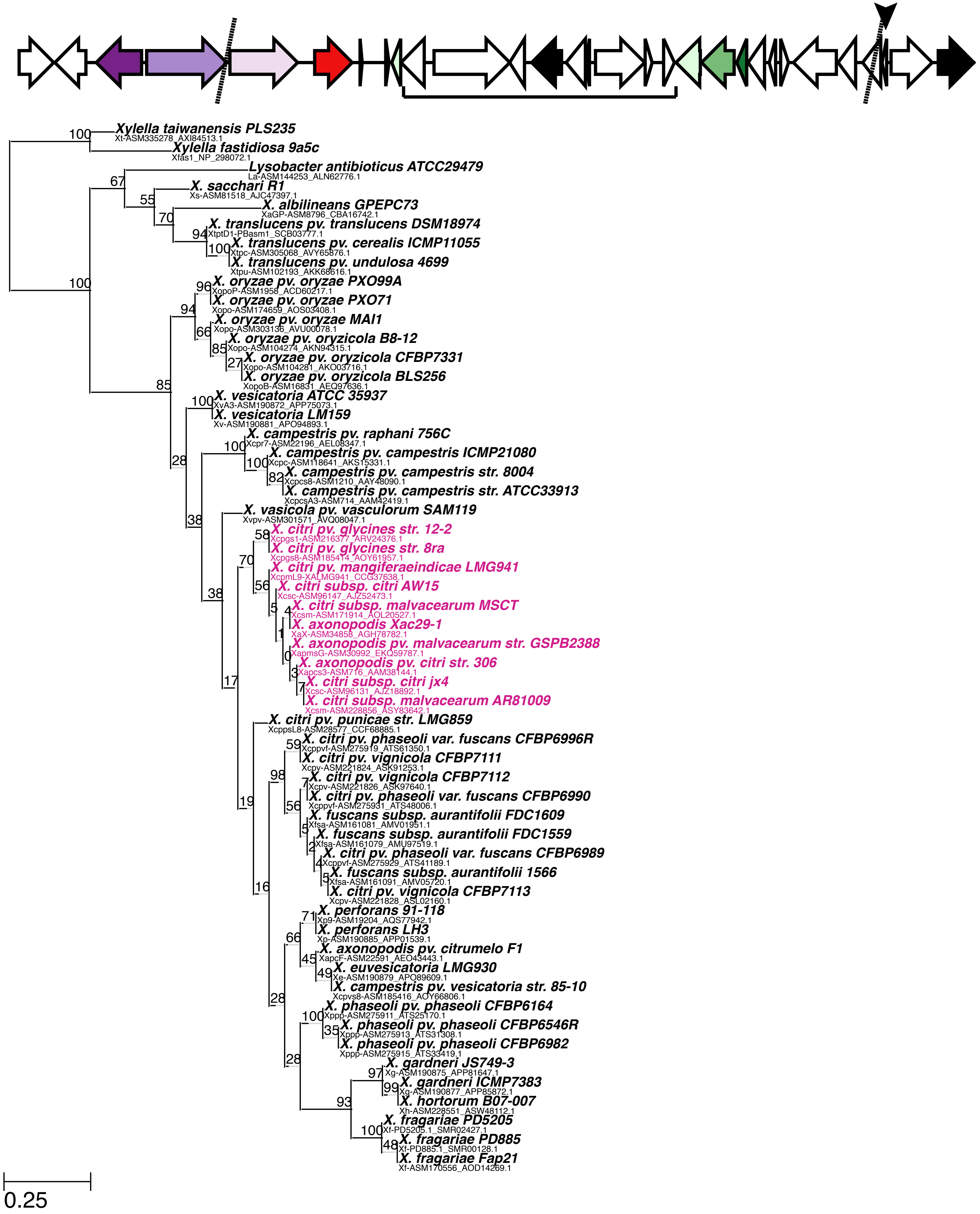

y) OG0001453 (esterase)

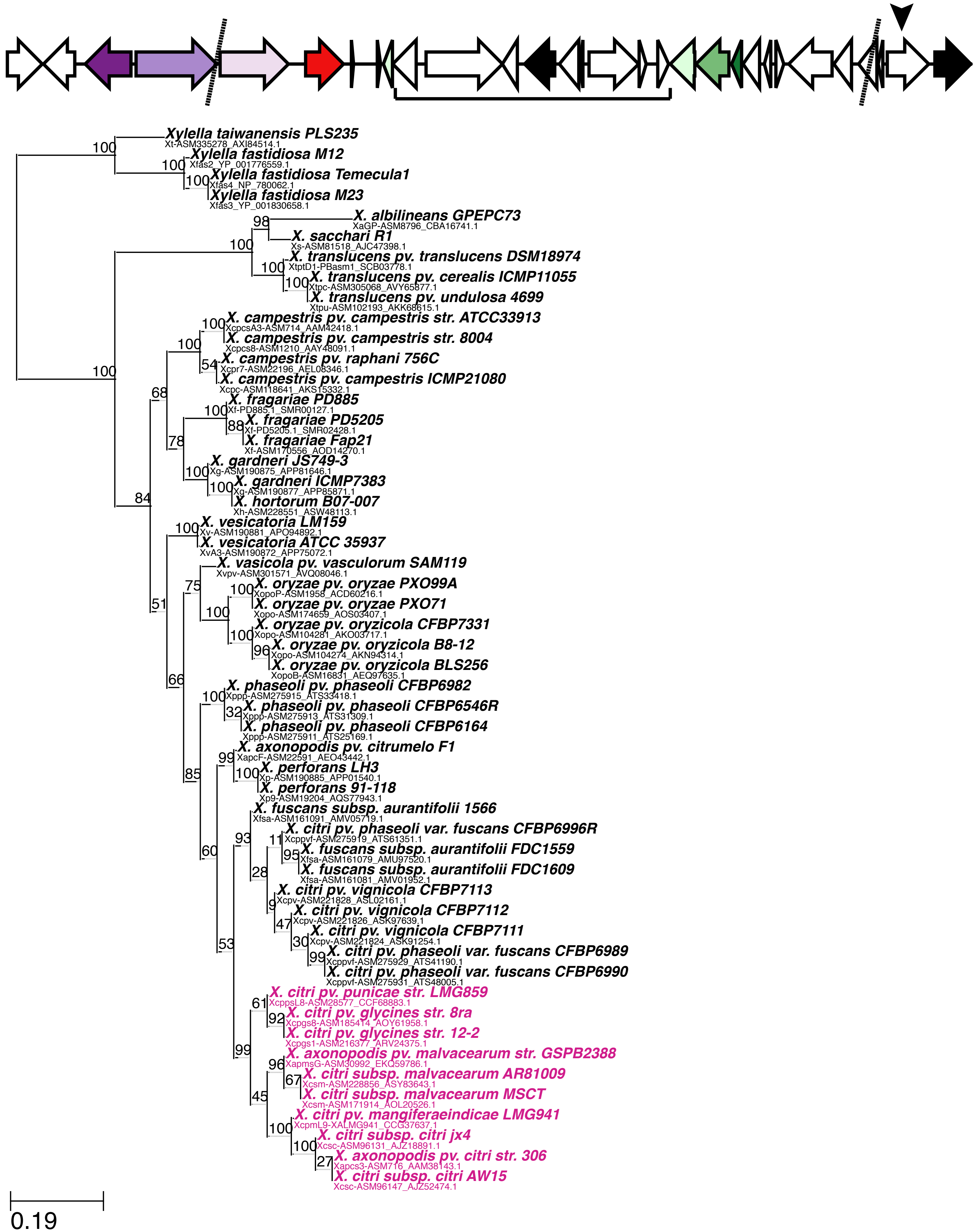
