## Supplemental Figure 8 for "Repeated gain and loss of a single gene modulates the evolution of vascular pathogen lifestyles"

a) OG0001552 (magnesium and cobalt transport protein)

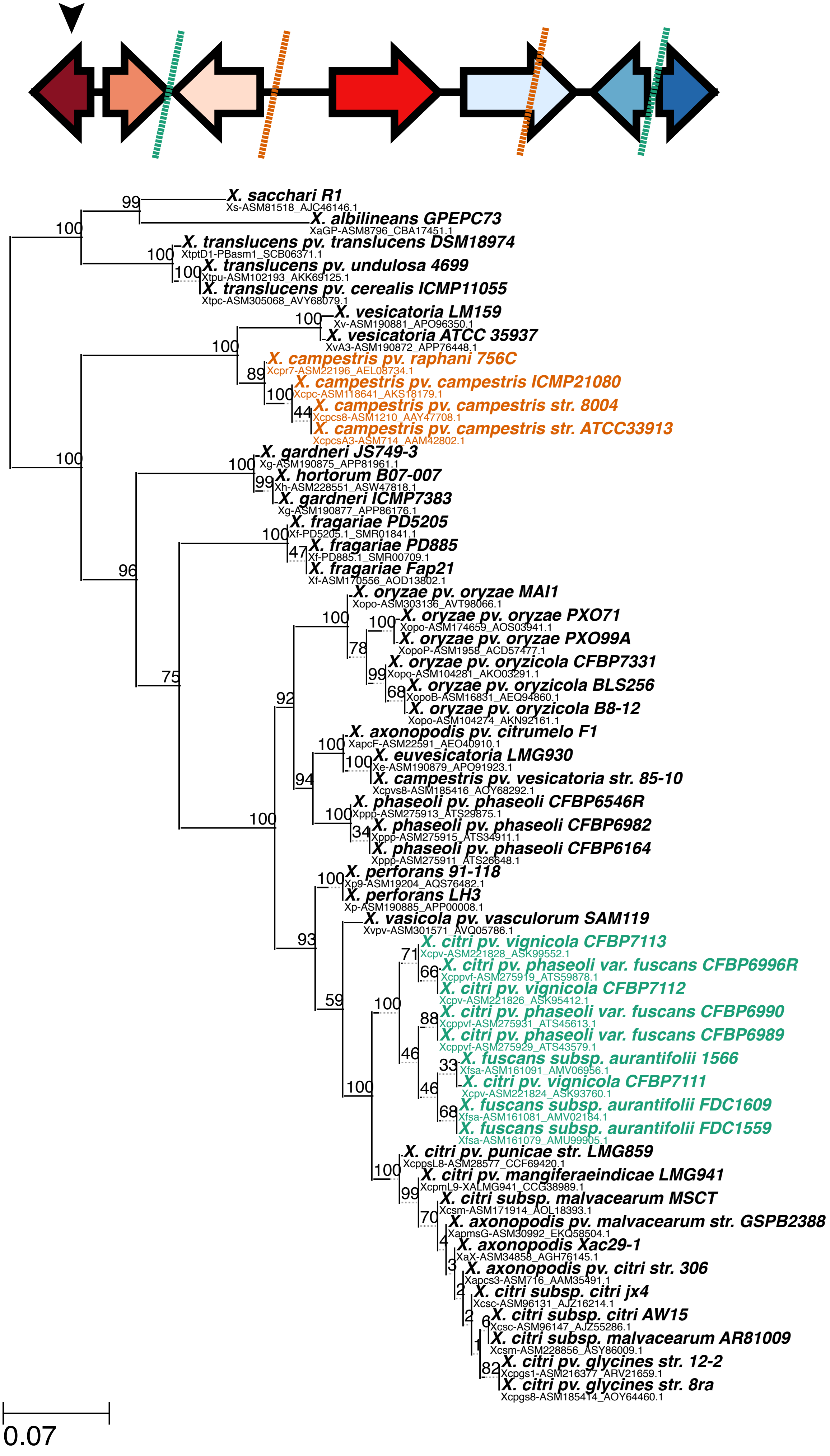

b) OG0002828 (function unknown)

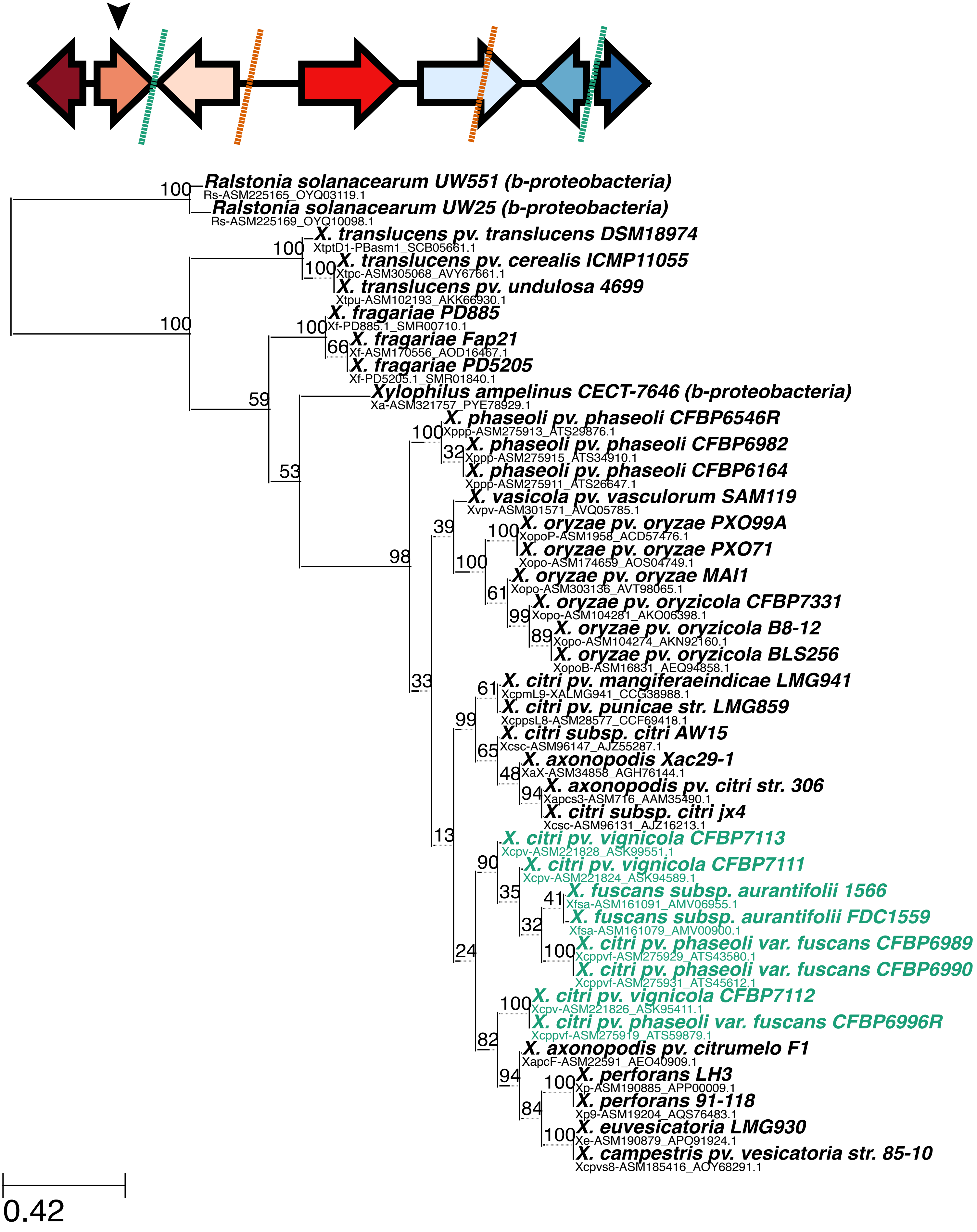

c) OG0000035 and 5' UTR, partition 2 (amino acid permease)

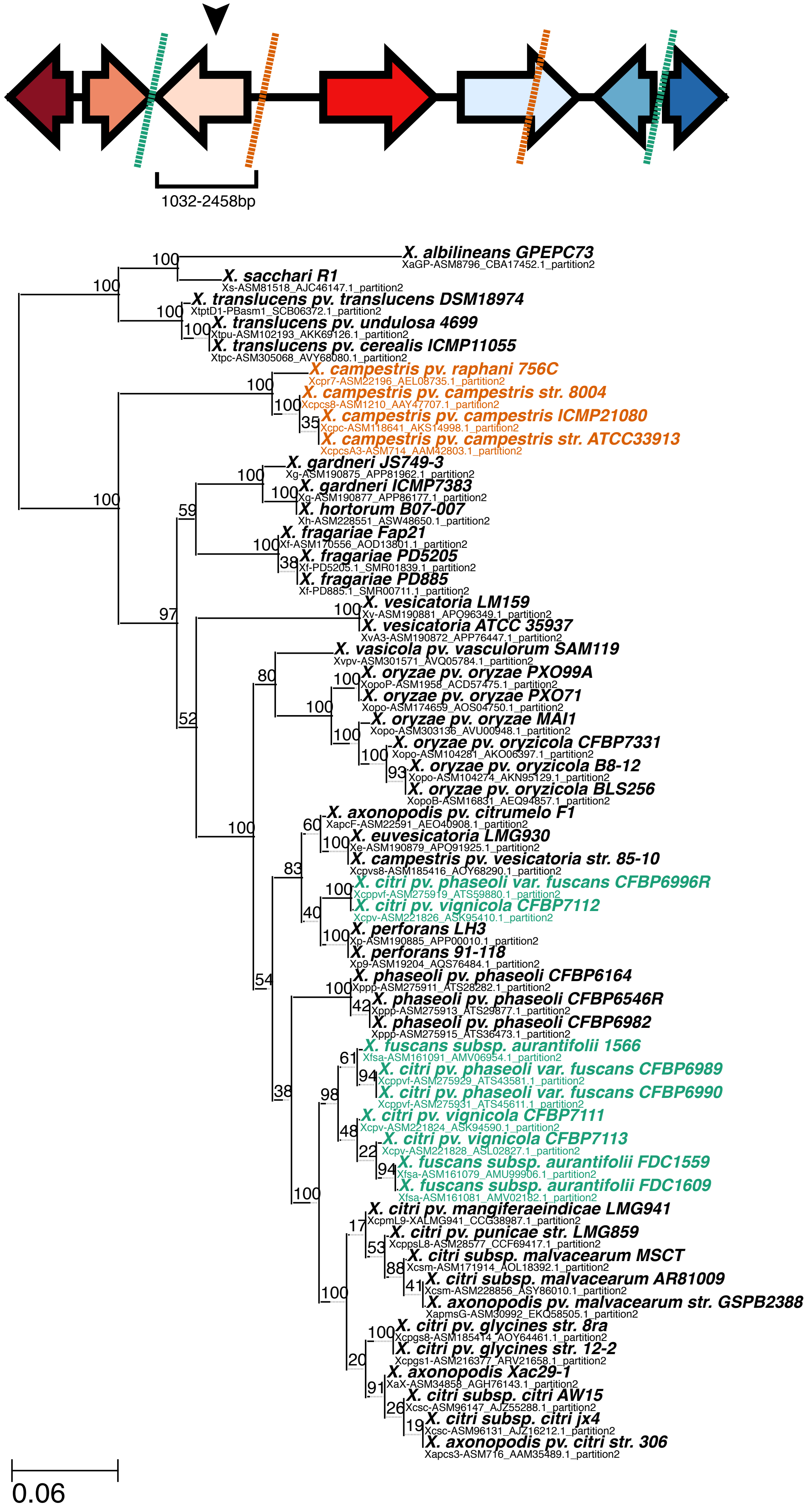

d) OG0000035 and 5' UTR, partition 1 (amino acid permease)

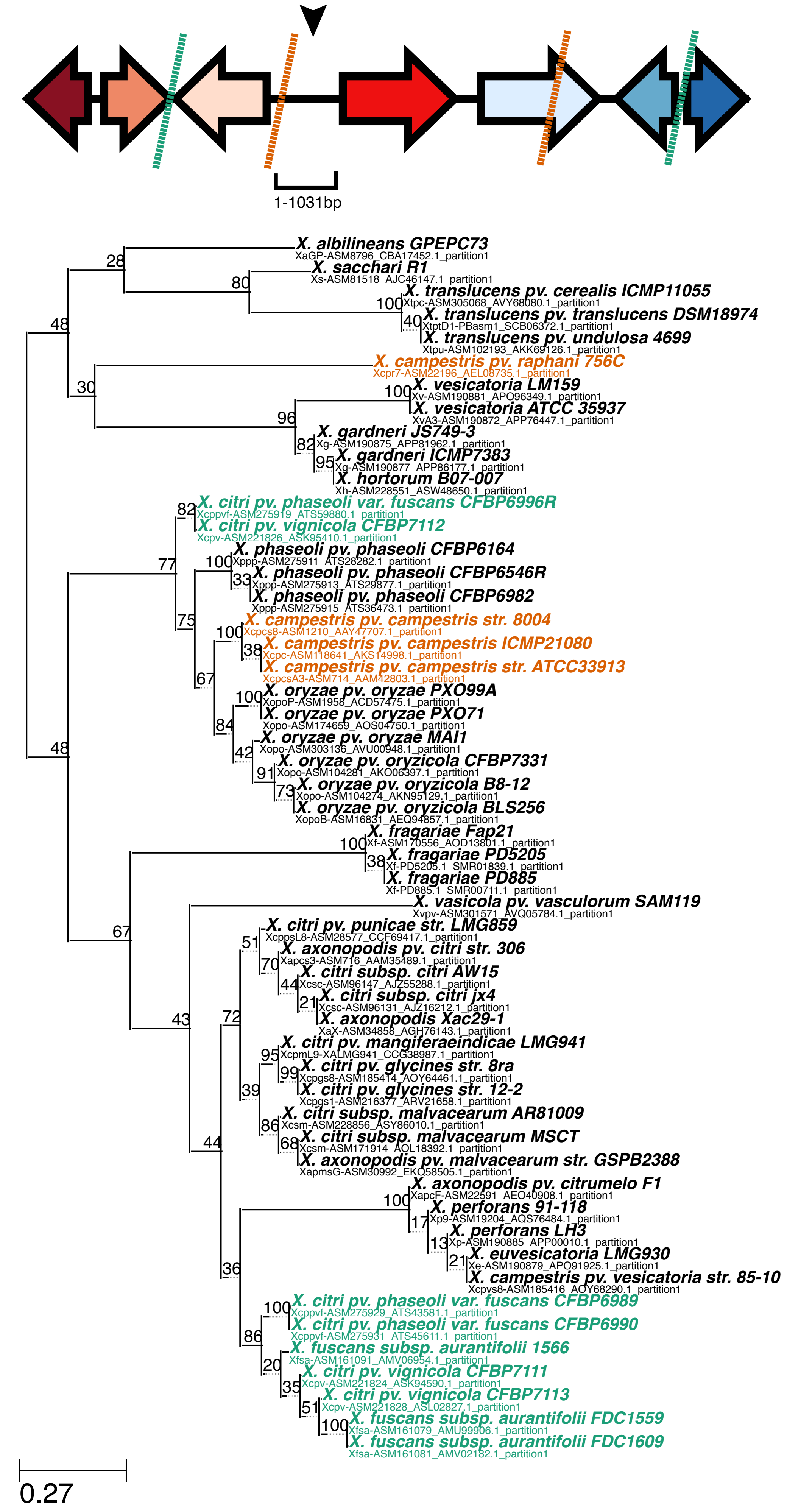

e) OG0000234, partition 1 (cellulase)

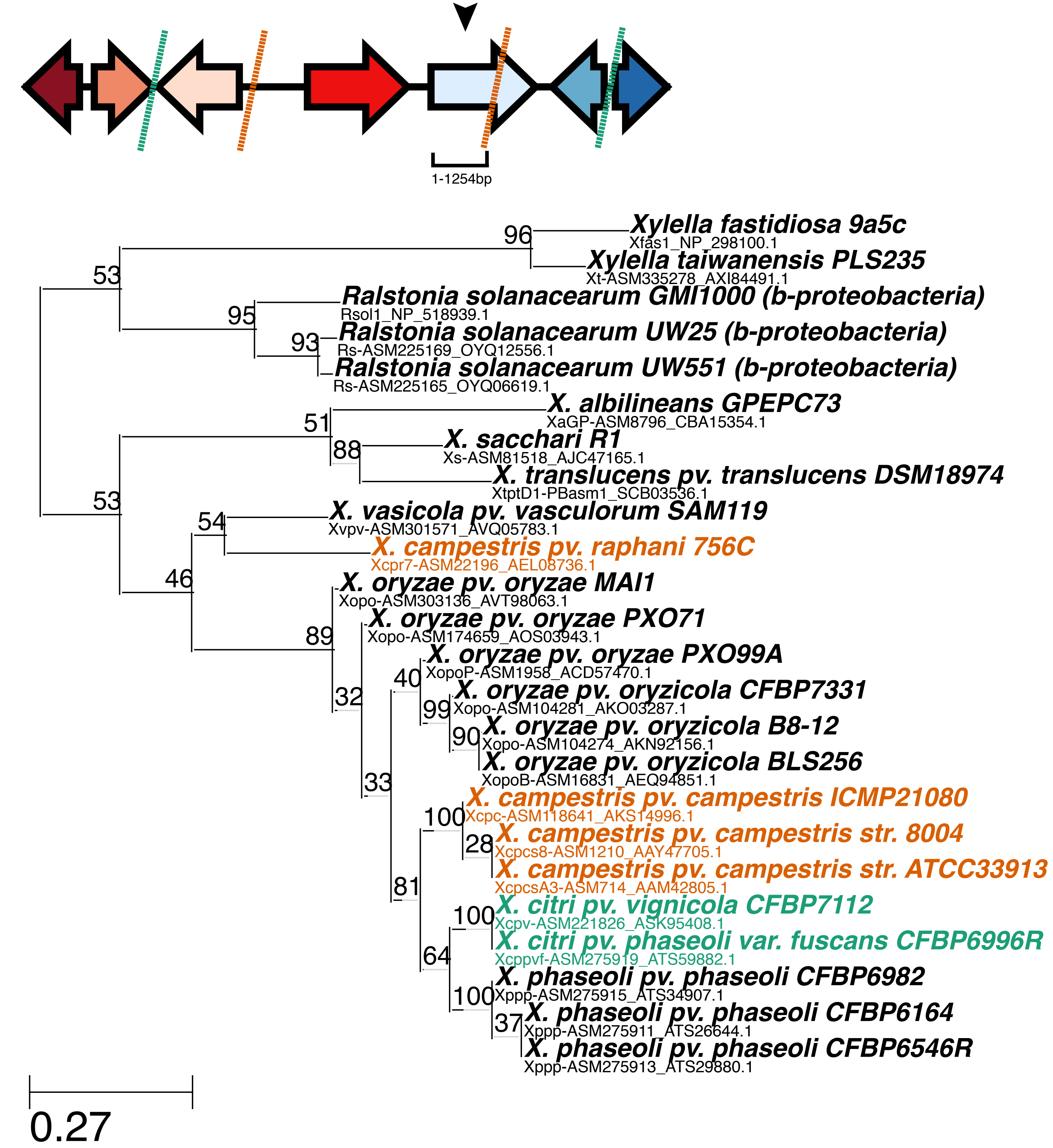

f) OG0000234, partition 2 (cellulase)

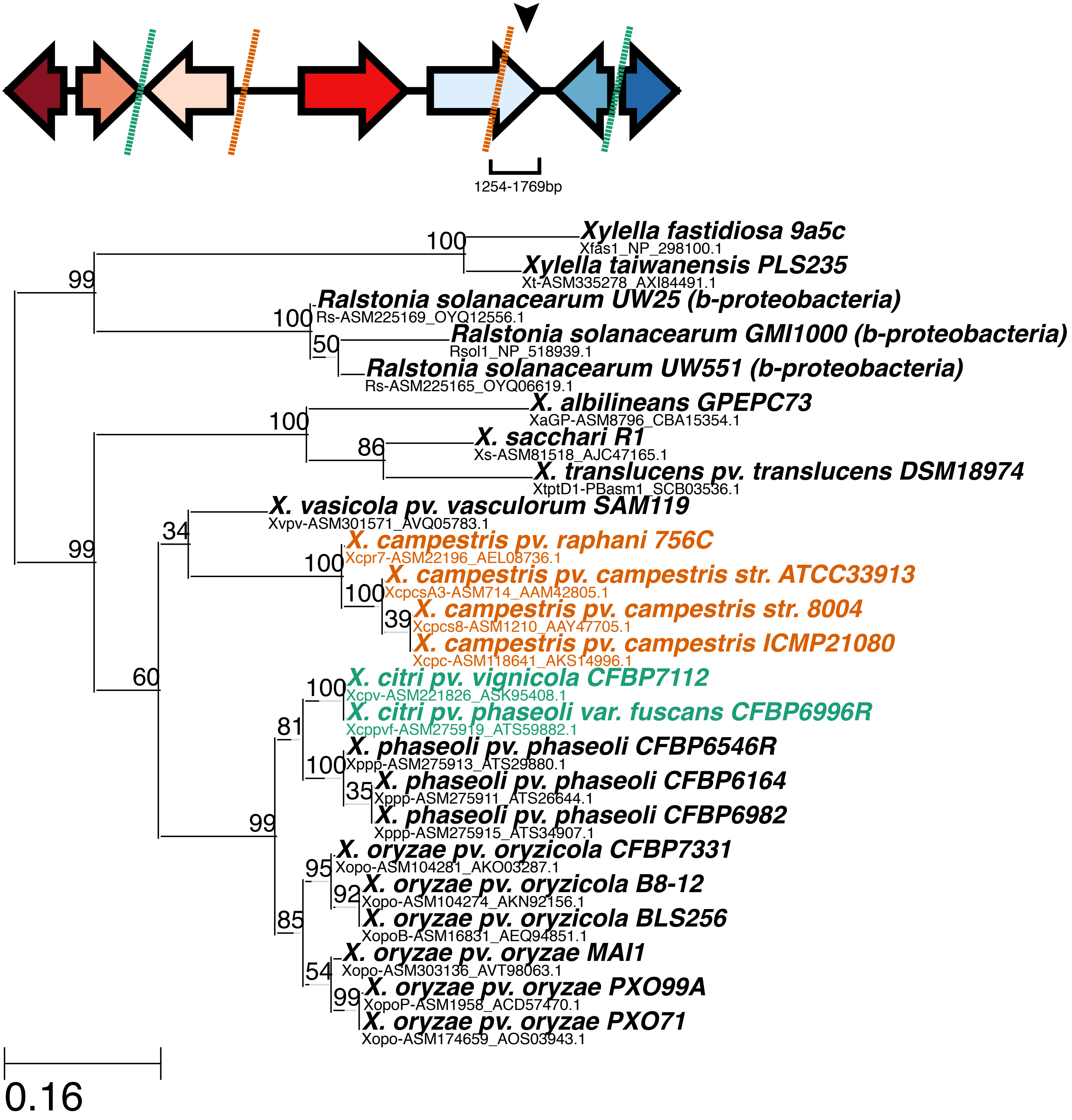

g) OG0002434 (integral membrane protein)

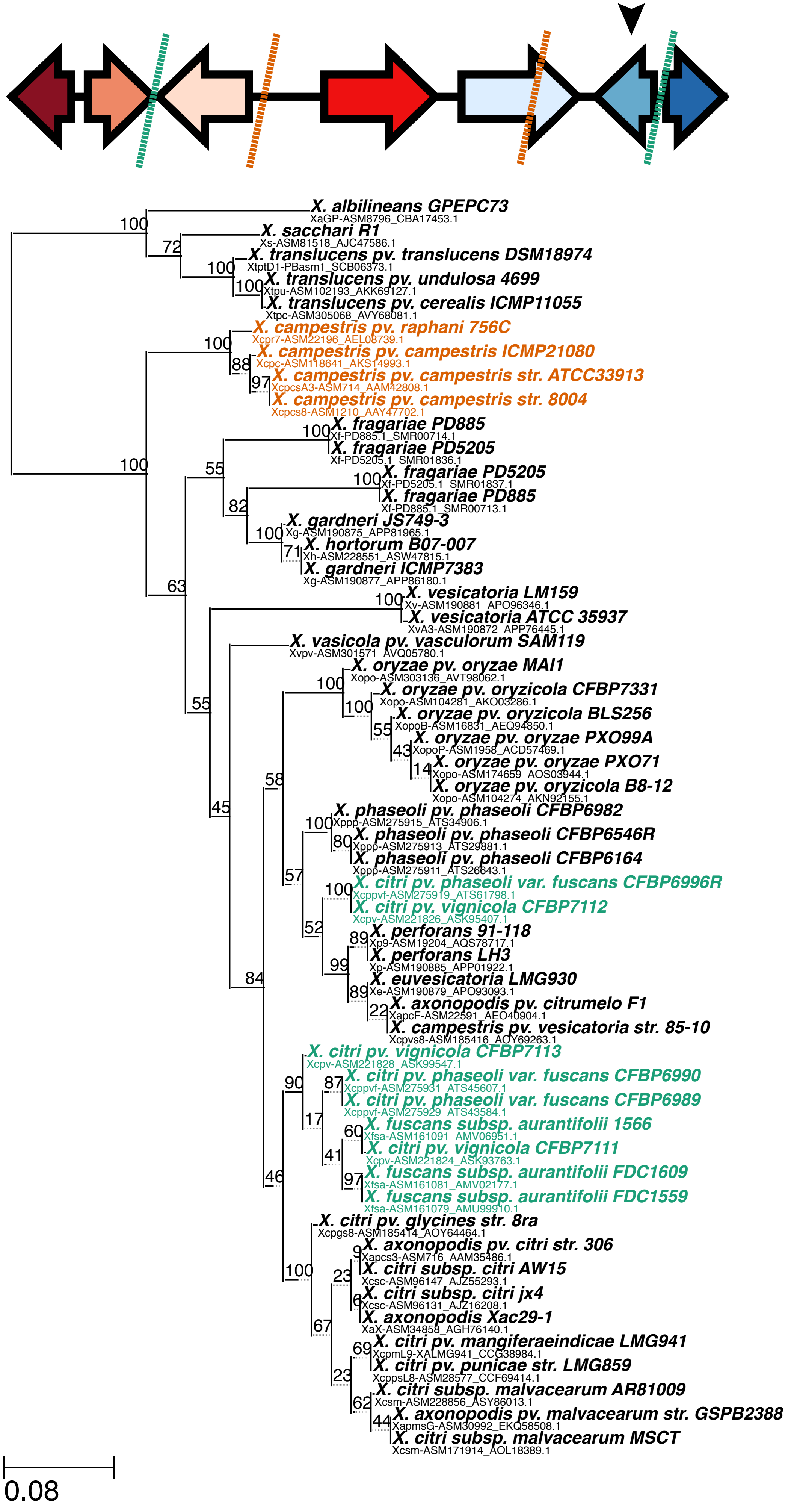

h) OG0002170 (methyltransferase)

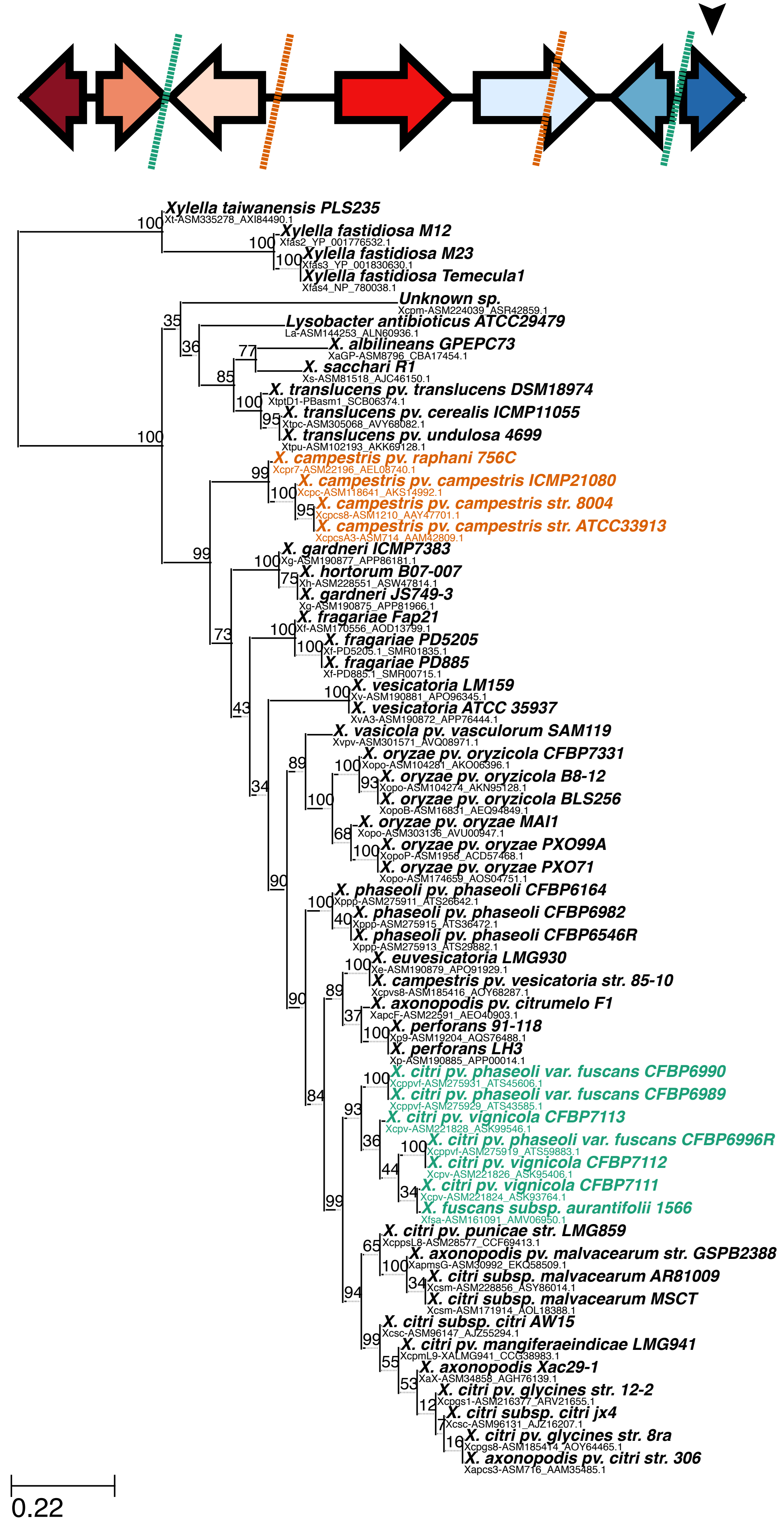
