## Supplemental Text with Figures for "Repeated gain and loss of a single gene modulates the evolution of vascular pathogen lifestyles"

**This PDF file includes:**

Materials and Methods

Figs. S1 to S8

Tables S1 to S9 (see additional file)

Materials and Methods

Bacterial strains and growth conditions

The bacterial strains used in this study are listed in Table S7. *Escherichia coli* strains were grown at 37°C in Luria-Bertani medium. *X.* *translucens* or *X. oryzae* cells were grown at 28°C on solid or liquid nutrient broth or peptone-sucrose rich media (*1*). When necessary, media were supplemented with gentamicin (15 µg/ml), kanamycin (25 µg/ml) or spectinomycin (50 µg/ml). See Table S7 for specific strains used in this study. *Xylella fastidiosa* subsp. *fastidiosa* str. Temecula1 wild-type (WT; Van Sluys et al., 2003) and *X. fastidiosa* subsp. *fastidiosa* str. Temecula1 Δ*cbsA* mutant were used in this study (Supplemental Table 9). Strains were cultured on PW (Davis et al., 1981) agar media, modified by removing phenol red and using 1.8 g/L of bovine serum albumin (BSA; Gibco Life Sciences Technology), for 7 days at 28ºC from -80ºC glycerol stocks, and subcultured onto fresh PW agar medium plates for another 7 days at 28^o^C before use. All assays were performed using the subcultured *X. fastidiosa* strains. PD3 (Davis et al., 1981) broth media and PBS buffer were used for suspending cells in liquid. *Escherichia coli* bearing the pUC4K plasmid was grown on LB (Luria-Bertani) medium (Bertani, 1951) at 37ºC. When needed, the antibiotic kanamycin (Km) was used at the concentration of 50 μg/mL.

Recombinant DNA techniques.

Total genomic and plasmid DNA were isolated by standard methods. *E. coli* and *Xanthomons* species were transformed as previously described (*2*). To construct complementation vectors of *cbsA*_Xtt_ and *cbsA*_Xoo_, the gene regions including the native promoters were PCR-amplified from *X. translucens* pv. tranlsucens str. UPB886. Each were cloned into pUC18miniTn7T to create pUC18miniTn7T::*cbsA*_Xtt_ and pUC18miniTn7T::*cbsA4*_Xoo_ (*3*). For gene expression, *X. translucens* pv. undulosa strains were transformed with miniTn*7* plasmids and pTNS1 to promote transposition and single gene insertion, and each was confirmed as described (*3*). We were unable to insert *cbsA* via miniTn*7* *X. translucens* pv. translucens strain UPB886. We therefore sequenced *X. translucens* pv. translucens ∆*cbsA* with long read PacBio sequencing. There were no notable differences in sequence between wild-type UPB886 and the ∆*cbsA* mutant. For visualization of bacteria by fluorescence microscopy, *Xanthomonas* bacteria (TableS7) were transformed with vectors for GFP expression (pNEO-GFP) (*4*). See Table S7 and S8 for specific strains and primers, respectively, used in this study.

The deletion of *cbsA* in *X. fastidiosa* strain Temecula1 (locus ID PD0529) was performed as described elsewhere (Kandel et al., 2018). Briefly, to obtain the targeting construct for site-directed mutagenesis, the upstream and downstream regions (905 and 968 bp, respectively) immediately flanking the *cbsA* gene were amplified using pairs of primers containing overlapping nucleotides with the Km resistance cassette present in the pUC4K plasmid (Table S9). The upstream and downstream regions of *cbsA* were fused to the Km resistance cassette through overlap-extension PCR, as detailed in Kandel et al. (2018). The purified PCR product was used for transforming WT strain through natural competence directly. Briefly, *X. fastidiosa* Temecula1 cells were suspended to OD_600_ of 0.25 (~10^8^ cells/mL) in PD3 broth, and 10 μL of this suspension was spotted together with 10 μL of the targeting construct on a PD3 agar plate. After five days of growth at 28ºC, cells were suspended into 1 mL of PD3 broth and plated into PW+Km agar for selection of mutants obtained through homologous recombination. Successful deletion of *cbsA* was confirmed through PCR (data not shown; primers shown in Table S9). In summary, non-amplification of an internal sequence of *cbsA*, and amplification of an internal sequence of the upstream region and the Km resistance cassette, confirmed deletion of *cbsA* in *X. fastidiosa* Temecula1 (WT was included as control). The obtained Δ*cbsA* mutant was stored as 25% glycerol stocks at -80ºC until use. The Gel/PCR DNA Fragments Extraction kit (IBI Scientific) was used for purification of PCR products and agarose gel fragments when needed. PCR reactions were performed using a standard protocol with the iProof High-Fidelity PCR kit (Bio-Rad) in a S1000 thermal cycler (Bio-Rad).

Plant growth conditions, inoculation methods and live imaging with confocal microscopy

Barley (*Hordeum vulgare* L. cv. Morex) were grown in growth chambers with cycles of 16 hours of light per day at 22-24^o^C. Rice (*Oryzae sativa*) were grown in growth chambers with 16 hours of light per day at 28^o^C 70% relative humidity or in the greenhouse. Plant seeds were directly germinated in potting mix. For barley, one leaf per plant was inoculated by leaf-clipping 7-10 days after seeds were planted with a water-based inoculum (OD_600_=0.1) or water as a control as previously described (*2*, *5*). For rice, one leaf per Nipponbare plants (three weeks old) were clip inoculated with *X. oryzae* or mutant (resuspension in water of OD600 0.1). Disease symptoms were assessed using at least five replications per condition with each experiment and each experiment was repeated at least three times. Symptom development was evaluated 21 or 15 days post-inoculation, respectively. *Xylella* Statistical was evaluated using an ANOVA or Student’s *t*-test.

For *X. fastidiosa* inoculation, plants were inoculated with PBS buffer (n=6), *X. fastidiosa* WT (n=9) or Δ*cbsA* (n=9). Disease severity and disease incidence were recorded weekly for 10 time points after first symptom appearance (8 weeks post inoculation). Briefly, disease incidence was considered as the percentage of plants showing at least one symptomatic leaf out of the total plants inoculated. Disease severity was calculated for each plant by counting symptomatic leaves and total number of leaves [(symptomatic leaves/total leaves) × 100] for each plant. The Area Under the Disease Progress Curve (AUDPC) was calculated by the midpoint rule method (Campbell and Madden 1990): AUDPC = Σ i^n-1^ [(y_i_ + y_i+1_)/2] (t_i+1_ –t_i_), where n = number of times disease assessment was performed, y = score of severity for each plant, and t = time of assessment.

For bacterial localization, barley plant leaves inoculated as above. Whole leaf tissue was imaged 5-14 days post inoculation with a Leica SP2 AOBS (Wetzlar, Germany) laser scanning confocal microscope with 40X oil objective. Barley leaves were cut directly adjacent to the inoculation zone for asymptomatic plants and immediately downstream of symptoms. Plant tissue was mounted onto a glass slide with water and covered with a glass cover slip. A 488 nm laser was used for GFP excitation and emitted fluorescence was collected between 505 and 540 nm. A 405 nm and 633 nm lasers were used for autofluorescence. Between 410 and 460 nm to define plant cell structures and between 650 and 700 nm for chlorophyll. Three to six plants were examined per treatment over three biological replicates.

**Supplemental Figure 1. Evolutionary relationship between vascular pathogenesis and a conserved cell wall degrading enzyme in *Xanthomonas* bacteria.** This figure is a modified Fig. 1 with detailed strain information for genomes analyzed. To explore the association of the vascular/non vascular lifestyle a set of publicly available complete and annotated genomes from different species in the Xanthomonadaceae family was analyzed. A pan genome SNP-based parsimony tree was built using kSNP3 (optimum kmer size = 21)(*6*). Genomes were classified as vascular (blue), non-vascular (yellow) or unknown (gray) based on available information in the literature. Ortholog groups for all annotated proteins were identified using Orthofinder (*7*), and a parsimony tree was generated based on pan-genome SNPs using KSNP3. Associations were identified between the presence/absence of each orthologue group in the analyzed genomes and the vascular/non-vascular trait according to the phylogeny using BayesTraitsV3. The likelihood that both traits (vascularity vs. gene presence) evolved dependently was compared to the likelihood they evolved dependently. Evidence of dependent evolution was assessed as Log Bayes Factors = 2(log marginal likelihood dependent model – log marginal likelihood independent model). Gene groups that were determined to evolve dependent on vascularity with very strong evidence (logBF >10) are shown, as well as the next top 12 genes below the threshold, genes are marked in red when present in a given strain. One gene group (OG0003492; CbsA) was commonly found in vascular strains, and the other (OG0002818; hypothetical) was more common in non-vascular genomes.

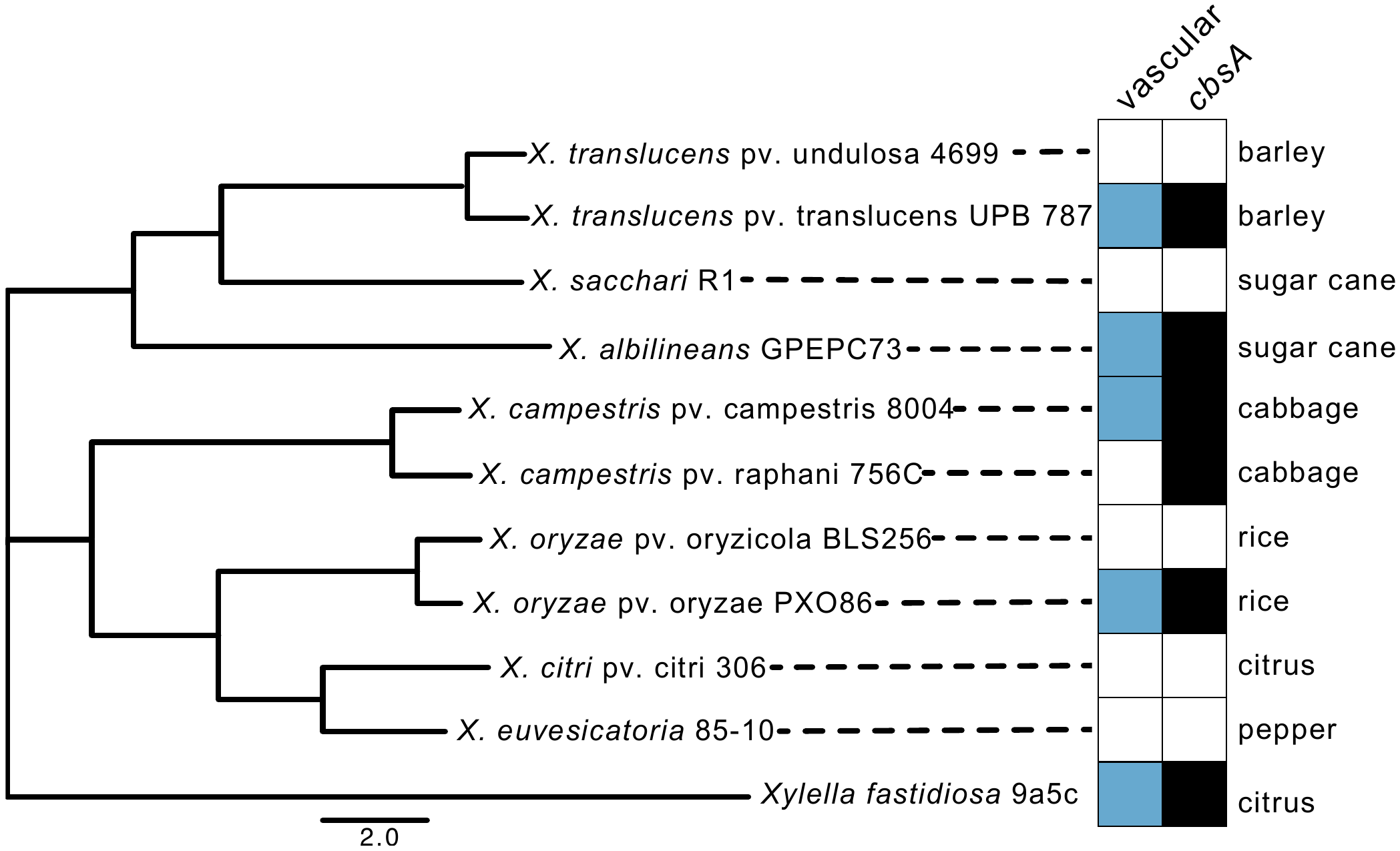

Supplemental Figure 2. Vascular, xylem pathogenesis strongly correlates with presence of *cbsA* but not host species. A phylogenetic tree was created based on representative Xanthomonad genomes from NCBI with Average Nucleotide Identity (ANI) (<http://enve-omics.ce.gatech.edu/g-matrix/>). Vascular, xylem colonizing bactetria are denoted in dark green. Light green boxes identify genomes with a *cbsA* homolog. Primary, characterized host for each pathogen is listed to the right of the boxes.

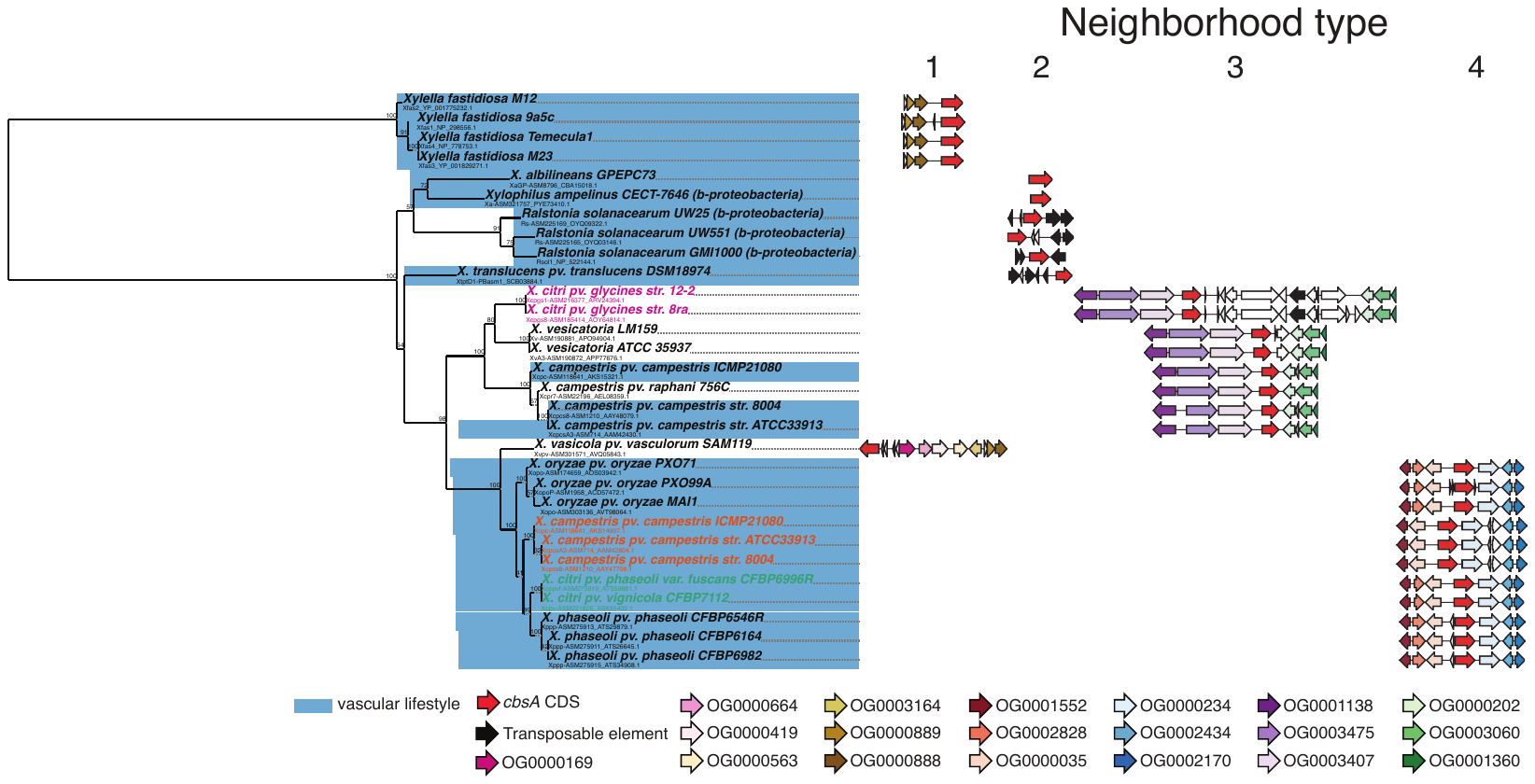

Supplemental Figure 3. The evolution and genomic context of *cbsA*. To the left is a nucleotide-based maximum likelihood phylogeny of *cbsA* homologs retrieved from the 86 genome database. Bootstrap support values (out of 100) are indicated above each bipartition. Each tip of the tree lists the full name of the isolate from which the sequence was retrieved in addition to the sequence’s accession number. To the right are schematics of the four distinct types of gene neighborhoods in which *cbsA* sequences are found. All schematics are drawn to scale within each column. Genes belonging to orthogroups of interest are color-coded (see legend at bottom), while all other intervening genes are left blank.

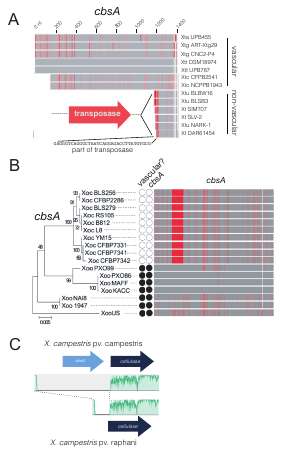

**Supplemental Figure 4. Specific inactivation events for *cbsA* homologs in *Xanthomonas* spp.** *cbsA* genes or genomic regions were aligned from vascular and non-vascular A) *Xanthomonas translucens*, B) *Xanthomonar oryzae* and C) *Xanthomonas campestris* with A&B) MAFFT alignment (www.benchling.com) and C) MAUVE. *cbsA* homologs were independently interrupted by three independent events: A) insertion, B) small deletion and C) complete gene loss. B) For Xanthomonas oryzae pathovars, the presence of CbsA was strongly correlated with vascular Xanthomonas oryzae pv. oryzae genomes but absent from non-vascular *X. oryzae* pv. oryzae. C) *X. campestris* pv. campestris is vascular, while *X. campestris* pv. raphanin is is non-vascular.

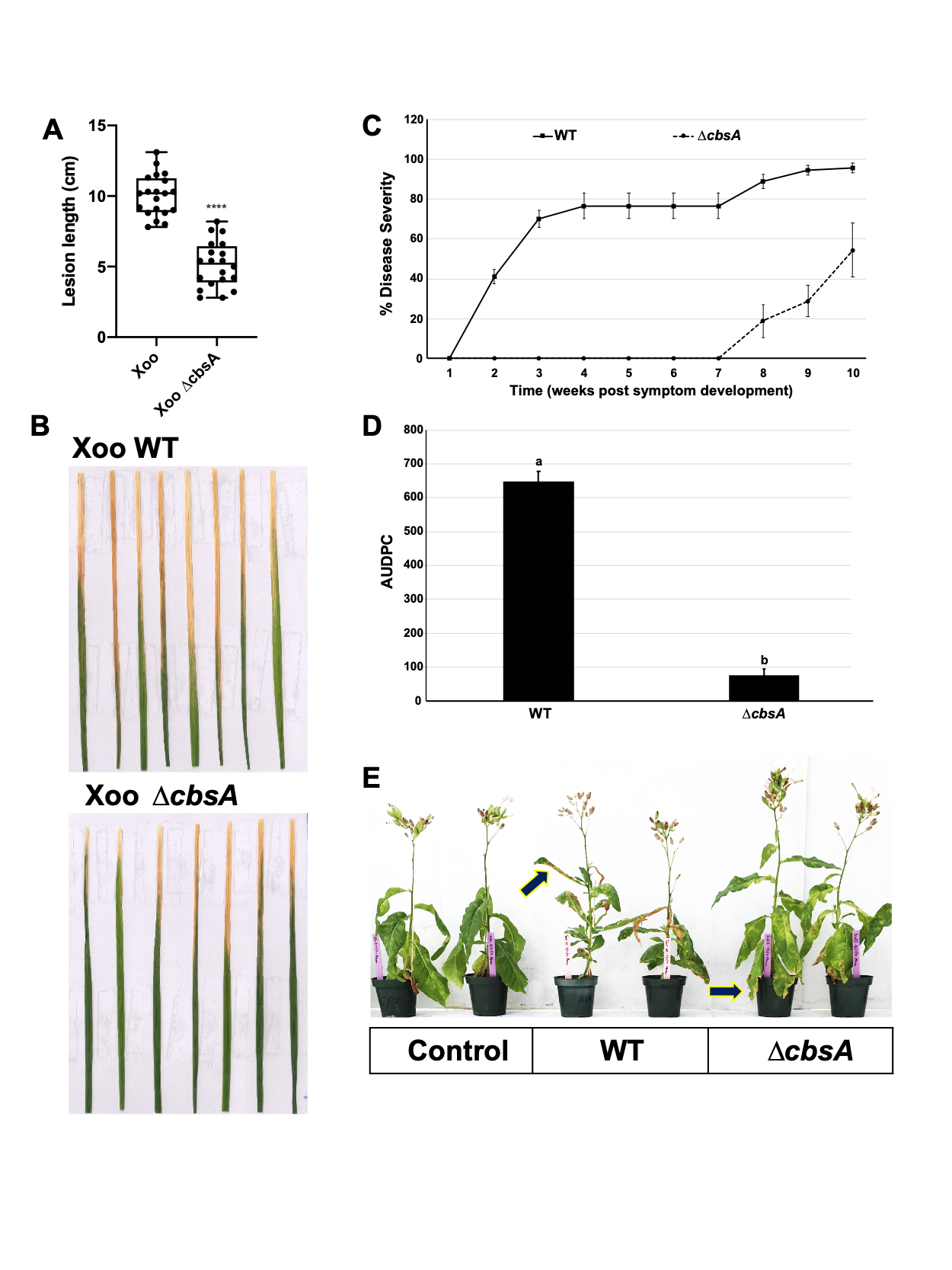

**Fig S5. Mutation of *cbsA* negatively affects virulence in *X. oryzae* pv. oryzae and *Xylella fastidiosa.*** A) Rice plants (cv. Nipponbare) were inoculated with *X. oryzae* pv. oryzae wild-type or ∆*cbsA* mutant. A-B) Lesion lengths were measured and imaged 15 days post inoculation. Lesion length was compared with Student’s *t-*test (*P*<0.0001)*.* C-E) Disease severity progression over time in inoculated tobacco plants. *Xylella fastidiosa* subsp. *fastidiosa* strain Temecula1 (WT) and mutant ∆*cbsA* were inoculated into *Nicotiana tabacum* L. cv. Petit Havana SR1 plants (PBS mock inoculation used as control). Leaf scorch symptoms were recorded for measurements of disease incidence and severity once a week during ten weeks after appearance of the first disease symptoms. At the final time point of evaluation, disease incidence in Temecula1 WT reached 100%, compared to mutant ∆*cbsA* reaching 66%; while disease severity reached 95% in WT and 54% in ∆*cbsA*. The mutant ∆*cbsA* showed delay of leaf scorch symptom development, with symptom appearance at the seventh week onwards and mostly restricted to lower leaves close to the inoculation point. Data represent means and standard errors from one experiment (n=9 for WT and ∆*cbsA*). D) Mean AUDPC per treatment group (WT and ∆*cbsA*). AUDPC was calculated using data from disease severity over ten weeks after first disease symptom appearance*.* AUDPC was lower for plants inoculated with ∆*cbsA*, in comparison to WT-inoculated plants. Data represent means and standard errors. Statistical significance was calculated using Tukey-Kramer HSD (*P*<0.05) (Statistical software JMP 15.0.1, 2015 SAS Inst. Inc., Cary, NC.). E) Representative image of leaf scorch symptoms in WT- and ∆*cbsA*-inoculated plants, as well as control plants (PBS-inoculated). Arrows in figures point to symptomatic leaves, which were distributed throughout the entire plant in WT-inoculated plants, and were mainly restricted to basal and middle leaves in ∆*cbsA*-inoculated plants.

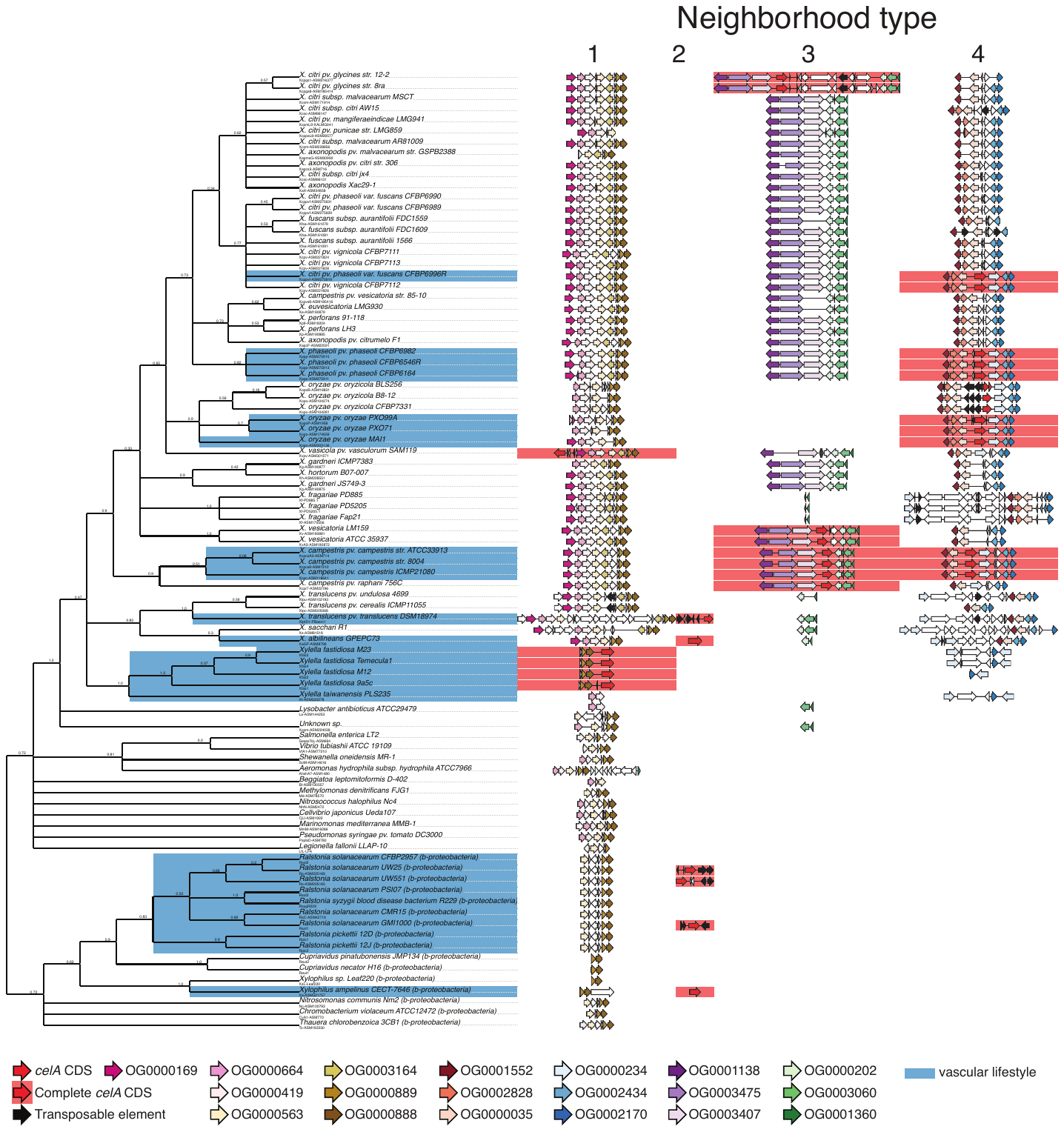
Fig. S6. The distribution of *cbsA* loci across beta- and gamma-proteobacteria (unedited version of Figure 1). Shown to the left is a majority rule consensus tree based on 81 maximum likelihood trees of single copy orthologs that summarizes species relationships among 86 bacteria examined in this study. Each bifurcation in the consensus tree is present in at least 50% of the single copy ortholog trees. Branch support values indicate internode certainty (ranging from 0-1), which quantifies the degree of conflict associated with a given bipartition across all 81 constituent trees. To the right of the tree is a graphic summarizing the distribution of the four distinct neighborhoods in which *cbsA* is found across each genome, in all cases whether *cbsA* is present or not. All neighborhood schematics are drawn to scale within each column. Genes belonging to orthogroups of interest are color-coded (see legend at bottom), while all other intervening genes are left blank.

See additional file attached (too large for supplemental document).

Fig. S7. Mid-point rooted, nucleotide-based maximum likelihood phylogenies of all genes in the type 4 *cbsA* neighborhood. Bootstrap support values (out of 100) are indicated above each bipartition. Each tip of the tree lists the full name of the isolate from which the sequence was retrieved in addition to the sequence’s accession number. Tree tips associated with sequences from *X. campestris* are colored orange, while tips associated with sequences from *X. citri pv. phaseoli, X. citri pv. vignicola* and *X. fuscans* are colored green. In topologies suggesting horizontal gene transfer (HGT), colored sequences were forced to be monophyletic in order to generate constrained topologies that would be expected under a scenario of vertical inheritance for phylogenetic hypothesis testing (Methods; Tables S4-5). A schematic of a *cbsA* gene neighborhood from *X. oryzae pv. oryzae* strain MAI1 is drawn above each tree, and a black vertical triangle indicates the current gene tree being displayed. Boundaries of the inferred homologous recombination events (Methods) are indicated by dashed lines, and are colored green for the HGT from the *X. phaseoli* clade to *X. campestris* and orange for the HGT from the *X. phaseoli* clade to *X. citri pv. vignicola CFBP7112* and *X. citri pv. phaseoli var. fuscans CFBP6996R*. a) OG0001552. b) OG0002828. c) OG0000035 and 5’ UTR, partition 2. d) OG0000035 and 5’ UTR, partition 1. e) OG0000234. f) OG0002434. g) OG0002170.

See additional file attached (too large for supplemental document).

Fig. S8. Mid-point rooted, nucleotide-based maximum likelihood phylogenies of all genes in the type 3 *cbsA* neighborhood, with midpoint rooting. Bootstrap support values (out of 100) are indicated above each bipartition. Each tip of the tree lists the full name of the isolate from which the sequence was retrieved in addition to the sequence’s accession number. Tree tips associated with sequences from *X. citri pv. glycines, X. citri pv. punicae, X. citri subsp. malvacearum, X. citri pv. mangiferaeindicae, X. citri subsp. citri* and *X. axonopodis pv. citri* are colored pink. In topologies suggesting horizontal gene transfer (HGT), colored sequences were forced to be monophyletic in order to generate constrained topologies that would be expected under a scenario of vertical inheritance for phylogenetic hypothesis testing (Methods; Table S6). A schematic of a *cbsA* gene neighborhood taken from *X. citri pv. glycines str. 8a* is drawn above each tree, and a black vertical triangle indicates the current gene being viewed. The boundaries of the inferred homologous recombination event (Methods) from *X. vesicatoria* to *X. citri pv. glycines* is indicated by black dashed lines. A black bracket indicates the boundaries of a 9-gene region that was likely inserted into the *cbsA* neighborhood after the HGT event. a) OG0003189. b) OG0003864. c) OG0001138. d) OG0003475. e) OG0003407. f) OG0006923. g) OG0000202. h) OG0012116. i) OG0006653. j) OG0004064. k) OG0005040. l) OG0015184. m) OG0004674. n) OG0005551. o) OG0003060. p) OG0001360. q) OG0000926. r) OG0002126. s) OG0001639. t) OG0001483. u) OG0001080. v) OG0000801, partition 2. w) OG0000801, partition 1. x) OG0001155. y) OG0001453.

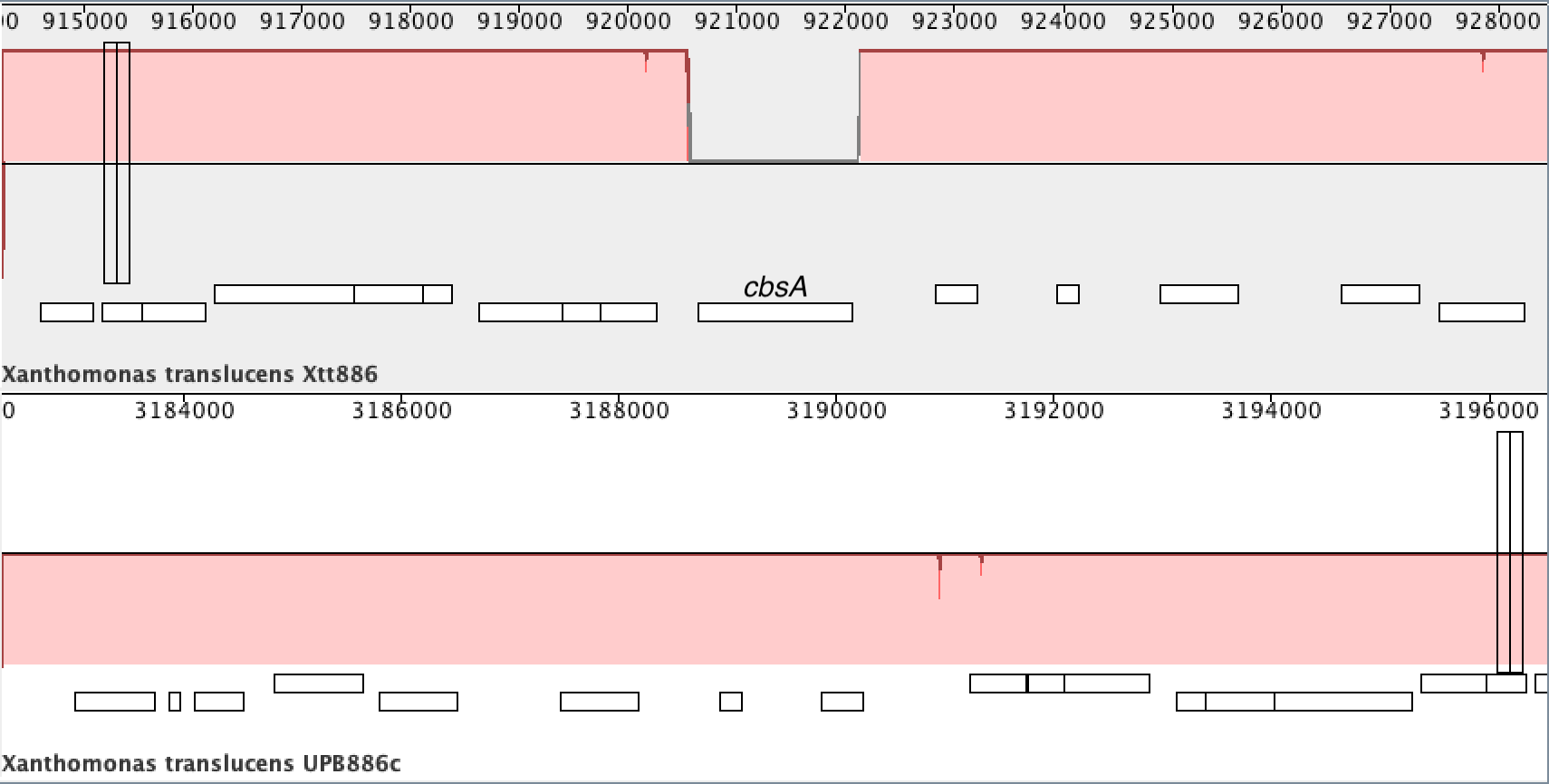

Fig. S9. Complete, whole genome sequencing validation of *X. translucens* pv. translucens ∆*cbsA*. We were unable to create a miniTn*7*::*cbsA* complementation of *X. translucens* pv. translucens UPB886 by transformation or conjugation. Therefore we performed whole genome sequencing to define the ∆*cbsA* mutation in UPB886. Genomic DNA from *X. translucens* pv. translucens ∆*cbsA* extracted with QIAGEN Genomic-tips 100G kit and sequenced by Psomagen, Inc using Pacbio RSII 20Kb SMRTbell. Assembly was done using Flye software with the parameters --pacbio-raw –g 5m (*8*). Genome annotation was done with Prokka (*9*). Genome comparisons and variant call was done using Mauve and NUCmer alignments (*10*). A MAUVE genome alignment of wild-type *X. translucens* UPB886 (Xtt886, top) compared to *X. translucens* pv. translucens ∆*cbsA* (UPB886c, bottom) demonstrates that the ∆*cbsA* gene was completely deleted by *sacB* mutagenesis for the *cbsA* loci (*11*, *12*).

Tables S1-9. See additional file attached for supplemental tables.
